# Regenerative capacity in *Drosophila* imaginal discs is controlled by damage-responsive, maturity-silenced enhancers

**DOI:** 10.1101/820605

**Authors:** Robin E. Harris, Michael J. Stinchfield, Spencer L. Nystrom, Daniel J. McKay, Iswar K. Hariharan

## Abstract

Like tissues of many organisms, *Drosophila* imaginal discs lose the ability to regenerate as they mature. This loss of regenerative capacity coincides with reduced damage-responsive expression of multiple genes needed for regeneration. Our previous work showed that two such genes, *wg* and *Wnt6*, are regulated by a single damage-responsive enhancer, which becomes progressively inactivated via Polycomb-mediated silencing as discs mature. Here we explore the generality of this mechanism, and identify numerous damage-responsive, maturity-silenced (DRMS) enhancers, some near genes known to be required for regeneration such as *Mmp1*, as well as near genes that we now show function in regeneration. Using a novel GAL4-independent tissue ablation system we characterize two DRMS-associated genes, *apontic* (*apt*), which curtails regeneration and CG9752/*asperous (aspr)*, which promotes it. This mechanism of suppressing regeneration by silencing damage-responsive enhancers at multiple loci can be partially overcome by reducing activity of the chromatin regulator *extra sex combs* (*esc*).

## Introduction

Tissue regeneration is a complex phenomenon that occurs in diverse taxa, and can result from a variety of non-exclusive mechanisms, including the generation of new tissue following the amplification of adult stem cells, de-differentiation of relatively mature cells to generate proliferating progenitor cells, and even rearrangement and remodeling of established tissue in the absence of proliferation (Tanaka and Reddien, 2011). Following tissue damage or loss, these processes, individually or together, promote the restoration of tissue size, structure and patterning, and are governed by coordinated programs of gene expression. However, in many organisms, regenerative capacity declines as an organism matures through development (Yun, 2015). Many examples of this phenomenon have been documented, such as the hind limbs of the anuran amphibian *Xenopus laevis*, which progressively lose the ability to recover from amputation as the tadpole develops through juvenile stages (Dent, 1962; Overton, 1963; Muneoka et al., 1986; Wolfe et al., 2000), or in mouse cardiac tissue, which can completely recover from partial surgical resection or induced infarction in newborn mice, while the same injury inflicted just 7 days later results in fibrosis and scarring (Porrello et al., 2011; Porrello et al., 2013). This striking loss of regenerative capacity over time is observed in diverse tissues of mammals (Reginelli et al., 1995; Porrello et al., 2011; Cox et al., 2014) including humans (Illingworth, 1974; King, 1979), as well as amphibians (Dent, 1962; Freeman, 1963; Beck et al., 2003; Slack et al., 2004) and even invertebrates (Smith-Bolton et al., 2009; Harris et al., 2016). Remarkably, many of these same tissues continue their program of developmental growth even after they lose the ability to regenerate, thus necessitating an explanation for how developmental and regenerative growth are regulated differently.

The imaginal discs of *Drosophila*, the larval primordia of adult structures such as the wing and eye, have been used extensively to study the genetic regulation of tissue growth and patterning. The ability of imaginal discs to regenerate was originally explored via classic transplantation studies (Ursprung and Hadorn, 1962), but more recently the use of genetic methods in which the discs are damaged *in situ* by the temporally and spatially limited expression of pro-apoptotic genes, have allowed large-scale experiments where the domain of tissue ablation can be regulated more precisely (Smith-Bolton et al., 2009; Bergantinos et al., 2010). Using these and other approaches, it was shown that imaginal discs readily regenerate at the beginning of the third larval instar (L3), but lose this ability over the course of L3 (Smith-Bolton et al., 2009; Halme et al., 2010; Harris et al., 2016). Multiple genes known to be upregulated in response to damage show less robust expression in more mature discs, which correlates with the loss of regenerative capacity. Recently it was shown that genome-wide changes in chromatin accessibility are associated with regeneration following genetically-induced cell death in wing discs (Vizcaya-Molina et al., 2018). However, these investigations were performed on discs at a single developmental stage when they still possessed high regenerative capacity, and therefore it remains to be seen how damage-induced changes to the epigenetic landscape might be altered in mature discs that have lost the ability to regenerate.

Wnt proteins play an important role in orchestrating regeneration in many organisms (reviewed in (Ricci and Srivastava, 2018). Using a genetic ablation system, we previously investigated the progressive decrease in damage-responsive *wingless* (*wg*) expression in wing-imaginal discs as they mature. Following damage, *wg* expression requires a damage-responsive enhancer, BRV118, located between *wg* and *Wnt6* (Schubiger et al., 2010; Harris et al., 2016). We showed that this enhancer contains a damage-responsive module (BRV-B) containing multiple binding sites for the JNK-responsive transcription factor AP-1 and that these sites are essential for its damage-responsive activity. An adjacent and separate element, BRV-C, has no enhancer activity on its own, but can silence the damage-responsive expression mediated by BRV-B in *cis* in a maturity dependent manner by promoting Polycomb-mediated silencing of the enhancer, characterized by highly localized H3K27 trimethylation. This localized epigenetic change, which spares more distant developmentally-regulated enhancers at the *wg*/*Wnt6* locus, provides a mechanism for selectively shutting off damage-responsive expression while preserving the ability of those genes to be expressed for normal development. Importantly, restoring *wg* expression in late L3 either by CRISPR/Cas9-mediated excision of the silencing element, BRV-C, or by expression of *wg* did not restore regeneration. This raises the possibility that multiple genes necessary for regeneration could be regulated similarly by damage-responsive enhancers that are also silenced in maturing tissues.

Using a genome-wide approach, we show here that a large number of genes, including many required for growth, are in the vicinity of regions of chromatin that are accessible in damaged discs in early L3, but are significantly less accessible in damaged discs of late L3. We show that some of these elements do indeed function as damage-responsive enhancers that are silenced as larvae mature. Using a GAL4-independent tissue ablation system that we have developed, we show that several genes associated with these elements are necessary for robust regeneration, thus demonstrating that the silencing of multiple such enhancers could account for the decrease in regenerative capacity as tissues mature. Proximity to such enhancers has also allowed us to identify novel regulators of regeneration. Finally, we show that modulating the activity of specific chromatin regulators that alleviate silencing at such enhancers can promote regeneration in mature discs.

## Results

### A damage-responsive and maturity-silenced (DRMS) enhancer is also present at the *Mmp1* locus

In this study we will be using the term <u>d</u>amage-<u>r</u>esponsive and <u>m</u>aturity <u>s</u>ilenced (DRMS) enhancers to denote enhancers that can promote gene expression in response to damage in immature (early L3) discs but not in mature (late L3 discs). To investigate the possibility that genes other than *wg* and *Wnt6* might be regulated by DRMS enhancers, we searched for modules with a similar bipartite organization to BRV118, the enhancer found in the *wg/Wnt6* locus. The damage-responsive (DR) module of BRV118, BRV-B, contains multiple AP-1 binding sites that are essential for its ability to respond to tissue damage, and which are also found in the corresponding enhancer regions of other *Drosophila* species (Figure 1A). In our previous analysis of the maturity silenced (MS) module, BRV-C, we showed that multiple elements in the module are required for silencing the enhancer in mature discs (BRV-C), since a series of deletions of the BRV-C fragment from one end results in progressive loss of its ability to silence damage-responsive expression by the adjacent BRV-B module (Harris et al., 2016). We also showed that Polycomb group (PcG) proteins are necessary for the silencing activity of BRV-C, suggesting that binding sites of the PcG DNA binding factor Pleiohomeotic (Pho) (Mohd-Sarip et al., 2002) might be important, one of which is present in BRV-C (Figure 1A). To identify other motifs that might be important for the function of both the DR and MS modules, we compared the sequences from four highly related *Drosophila* species and found several conserved regions (Figure 1A). Screening for stretches of identical DNA sequence 50bp or larger, we identified a single region (region 1) within BRV-B that is close to the three AP-1 binding sites that we previously showed are required for damage-responsive expression (Harris et al 2016). By comparison, the BRV-C module has multiple completely-conserved regions, including two regions over 100bp in length (regions 6/7 and 8), one of which (region 8) contains the conserved Pho binding site (Brown et al., 1998; Zhu et al., 2011). By conducting BLAST searches of the genome using sequences from these highly conserved regions, we found an exact copy of a 17 bp sequence from region 6/7 present within an enhancer previously identified in the *Matrix metalloproteinase 1* (*Mmp1*) locus (Figure 1B). We compared this 17bp sequence with a library of known *Drosophila* transcription factor binding sites using the TOMTOM motif comparison tool (Gupta et al., 2007) and found that within the 17bp sequence strongly matches an Sp1 binding site (Figure 1B). Sp1 binding sites are known to be required for the activity of Polycomb Response Elements (PREs) and have been identified in most molecularly characterized PREs (Brown et al., 2005), while an Sp1 family member, Ssps, has been shown to bind different PREs and contribute to silencing (Brown et al., 2005; Brown and Kassis, 2010). In the ∼5 kb surrounding this motif at the *Mmp1* locus are two conserved Pho binding sites and six AP-1 binding site (Figure 1B). In contrast to the Wnt enhancer, there are three AP-1 binding sites between the 17 bp motif and the PRE (Figure 1B), which could potentially indicate that there is not a clear separation between damage-responsive and silencing motifs at the *Mmp1* locus.

**Figure 1:**
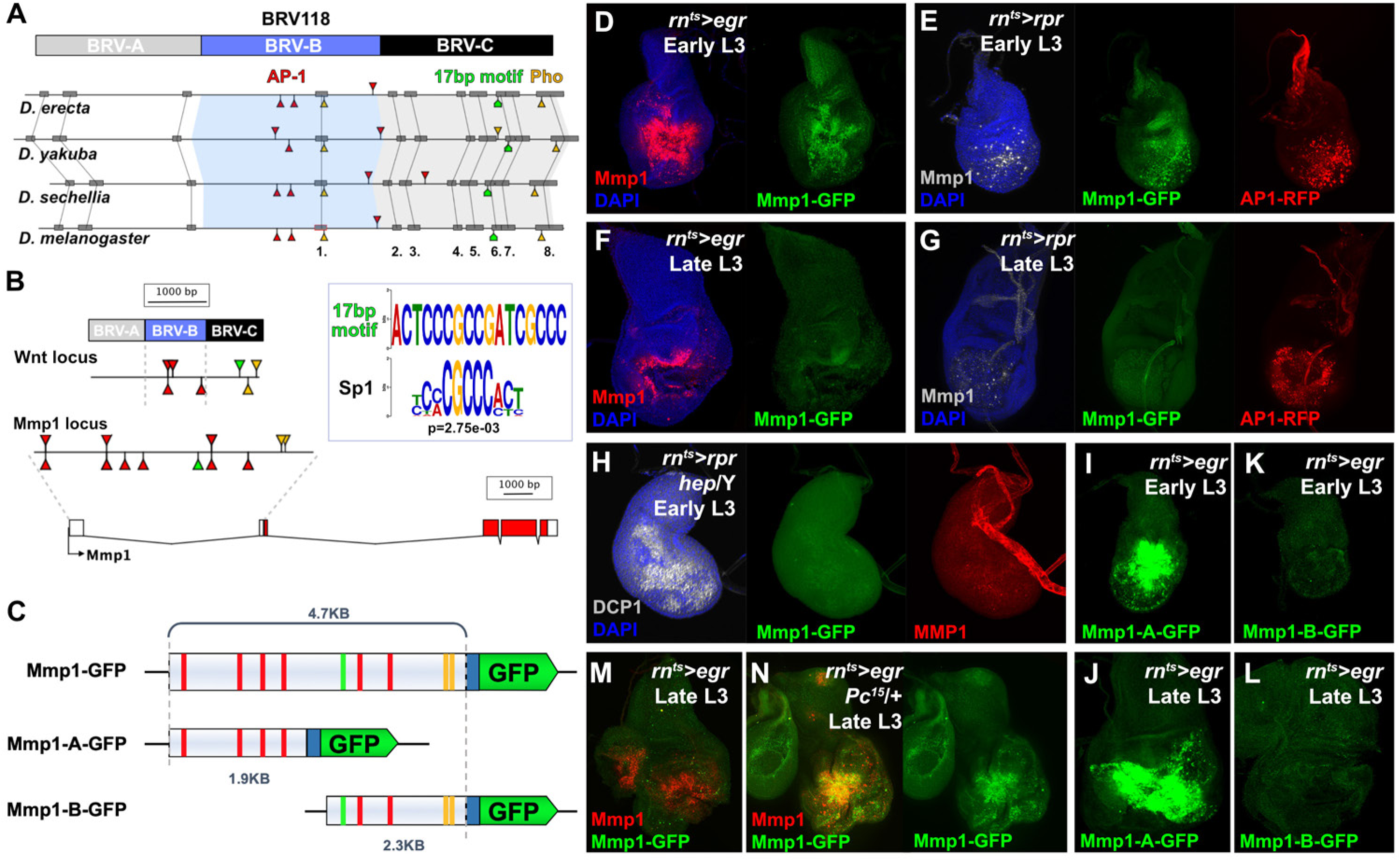
*Mmp1* is regulated by a bipartite damage-responsive enhancer with organizational similarity to DRMS^Wnt^, **(A)** Schematic illustrating the conservation of the BRV118 Wnt enhancer (top) in four *Drosophila* species. The main damage-responsive region, BRV-B (blue box), the maturity silencing region, BRV-C, (black box) and their equivalent sequences in other species are indicated. Matching sequence of 50bp or greater are shown as gray boxes and numbered. Also indicated are AP-1 binding sites (red arrowheads), Pleiohomeotic (Pho) sites (yellow arrowheads) and the conserved 17bp motif (green markers). Binding site orientation is indicated by appearance above or below the line, **(B)** Schematic comparing the BRV118 enhancer from the Wnt locus (top) with that of a putative enhancer at the *Mmp1* locus (bottom). AP-1 and Pho binding sites, and the 17bp motif consensus, are illustrated in both DNA sequences, as in (A). The 17bp motif and the Sp1 motifs are shown (inset), **(C)** Schematics of the *Mmp1-*GFP and related reporters. AP-1 binding sites (red bars), Pho binding sites (yellow bars) and the 17bp motif (green bars) are indicated. Blue box: *hsp70* minimal promoter, **(D)** Early L3 wing imaginal disc following ablation with *rn^ts^>egr* stained for Mmp1 (red) and DAPI (blue), and showing activity of the *Mmp1-GFP* reporter (green), **(E)** Early L3 wing disc following ablation with *rpr*, showing levels of Mmp1 (gray), the activity of the *Mmp1-GFP* reporter (green) and the *AP-1-RFP* reporter (red), **(F)** Late L3 wing disc following *egr* ablation, showing that both the damage-induced Mmp1 (red) and *Mmp1-GFP* reporter expression (green) is weaker than that of early L3 discs, **(G)** Late L3 wing disc ablated with *rpr* as in (E), showing expression of both *Mmp1* and the reporter are weaker in late L3 discs, while *AP-1-RFP* remains strongly activated on both days, DAPI: blue, **(H)** Early L3 hemizygous *hep^-^* mutant wing disc following *rpr* ablation, showing that neither Mmp1 (red) or the *Mmp1-GFP* reporter (green) is activated despite damage, indicated by dead cells (DCP1, gray), **(I-J)** Early L3 (I) and late L3 (J) wing discs bearing the *Mmp1-A-GFP* reporter (green) following *egr* ablation. The reporter is strongly activated, even in mature discs, **(K-L)** Early L3 (K) and late L3 (L) wing discs bearing the *Mmp1-B-GFP* reporter (green), showing no activity following *egr* ablation, **(M-N)** Mmp1 protein (red) and *Mmp1-GFP* expression (green) in late L3 *egr* ablated discs in a wild type (M) or *Pc^15^* heterozygous mutant background (N), showing increased levels of Mmp1 and GFP with reduced *Pc* gene function.

To investigate this region, 4.7KB of DNA upstream of the *Mmp1* coding sequence, which includes the 17bp motif, and the AP-1 and Pho binding sites, was cloned upstream of a minimal promoter and GFP coding sequence. This construct was used to generate a transgenic reporter, *Mmp1-GFP* (Figure 1C). The reporter showed little activity during normal development in undamaged discs, only weakly recapitulating *Mmp1* expression that normally occurs in the developing air sac in late L3 discs (Wang et al., 2010) (Figure 1 – Figure supplement 1A). In contrast, upon genetic ablation in early L3 discs (day 7) using *rn-GAL4*, *Gal80^ts^*, *UAS-eiger* (hereafter *rn^ts^>egr*, see Materials and Methods), the reporter is strongly activated in a pattern that resembles endogenous damage-induced Mmp1 protein (Figure 1D). A similar pattern of expression, albeit weaker, is observed when *reaper* (*rpr*) is used instead of *egr* to kill cells, and this expression is coincident with an AP-1 reporter (Figure 1E). Ablation in the absence of JNK activity using a *hep* mutant background fails to induce the reporter or *Mmp1* (Figure 1H). Thus, JNK signaling is a necessary input into the enhancer, as it is for damage-induced *Mmp1* expression. Conversely, ectopic activation of JNK signaling through expression of *hep^CA^* leads to strong reporter activation (Figure 1 – Figure supplement 1B). Physical wounding of these discs followed by *ex vivo* culture also results in activation of the *Mmp1* reporter at the wound edge, coincident with *Mmp1* expression (Figure 1 – Figure supplement 1C). Consistently, the *Mmp1-GFP* reporter also recapitulates the weaker expression of *Mmp1* in response to genetic ablation with either *egr* or *rpr* in late L3 discs (day 9, Figure 1F-G), despite a robust level of JNK activity, as indicated by the AP-1 reporter (Figure 1G). Together these data indicate that this region of the *Mmp1* locus contains an enhancer that is both damage-responsive and silenced with maturity.

To directly test whether this DRMS enhancer has separable damage-activated and maturity-silencing elements, two reporter lines, *Mmp1-A-GFP* and *Mmp1-B-GFP*, were generated using enhancer fragments (Figure 1C) and inserted into the same transgene landing site as the original *Mmp1-GFP* to make their activity directly comparable. *Mmp1-A* was strongly activated specifically in response to damage, more so than the full-length enhancer (Figure 1I-J). Moreover, in the absence of the *Mmp1-B* sequences it can be activated equally as strongly in both early and late L3 discs (Figure 1I-J). In contrast, *Mmp1-B*, which contains the 17bp motif and conserved Pho binding site yielded no enhancer activity in ablated young or old discs, despite containing three predicted AP-1 binding sites (Figure 1C, K-L). Further subdivision of the *Mmp1-A* region showed that the majority of damage-responsive expression is driven by a ∼1kb section of DNA bearing three high consensus AP-1 binding sites (Figure 1 – Figure supplement 2A-C). None of the generated reporters showed significant expression in the absence of ablation (Figure 1 – Figure supplement 2E-I). We also examined the activity of the *Mmp1-GFP* reporter in a PcG mutant background, which we previously showed de-repressed the enhancer in the *wg*/*Wnt6* locus (hereafter DRMS^Wnt^) in older damaged discs compared to wild type (Harris et al., 2016). Expression of *Mmp1-GFP* and Mmp1 protein in a *pc^15^*/+ ablated late L3 disc was significantly stronger compared to the wild type control (Figure 1M-N), indicating that Polycomb-mediated epigenetic silencing is necessary to limit *Mmp1-GFP* activation in mature tissues. Thus, the functional organization of this enhancer is very similar to the one we have characterized at the *wg*/*Wnt6* locus, with clearly separable damage-responsive modules and silencing modules. Also like DRMS^Wnt^, its age-dependent silencing is dependent upon PcG function. Due to the similarity of organization with DRMS^Wnt^, we will henceforth refer to this region as DRMS^Mmp1^. Thus, we have shown that another gene that has been demonstrated to function in both growth and tissue remodeling is regulated by a DRMS enhancer.

### Identification of DRMS enhancers genome-wide using ATAC-seq

While DRMS^Mmp1^ and DRMS^Wnt^ share sequence motifs, it is possible and even likely that other such enhancers rely on different mechanisms to be activated or silenced, and thus potentially lack these motifs. To search for such enhancers without a bias for any specific sequence motif, we employed ATAC-seq, a genome-wide assay of chromatin accessibility (Buenrostro et al., 2015), which utilizes the frequency of integration of a transposon as a measure of open chromatin. We compared chromatin profiles of *rn^ts^>egr* ablated wing discs from both early and late L3 larvae (reflecting times of high and low regenerative capacity, respectively), as well as from identically-staged unablated discs (Figure 2A). We performed three full biological repeats for each condition, yielding a final total of 14,142 open chromatin peaks after merging overlapping peaks from each condition (see Materials and Methods for full details of quality control and data analysis parameters). To assess the overall similarity of chromatin landscapes between the different samples, we performed Pearson correlation analysis on CPM-normalized counts from DEseq2 (Figure 2 – Figure supplement 1A), which shows a high correlation between replicates (r>0.9) and demonstrates that the data cluster first by developmental stage and then by whether or not the tissue had been damaged. Thus, changes in the chromatin landscape are greater between discs at different developmental time points than that of similarly aged discs that differ in their damage status. This could either be because maturation during L3 impacts chromatin status more than the effect of damage, or because only a subset of cells in the disc are strongly affected by damage while all cells are affected by disc maturation. We noted that one of the three late L3 damaged samples clusters with the early L3 samples. The reason for this is unclear. However, we have done separate analyses both with and without this sample and our conclusions are not appreciably altered (not shown). We have therefore not removed this sample from any of the analyses shown in this manuscript.

**Figure 2:**
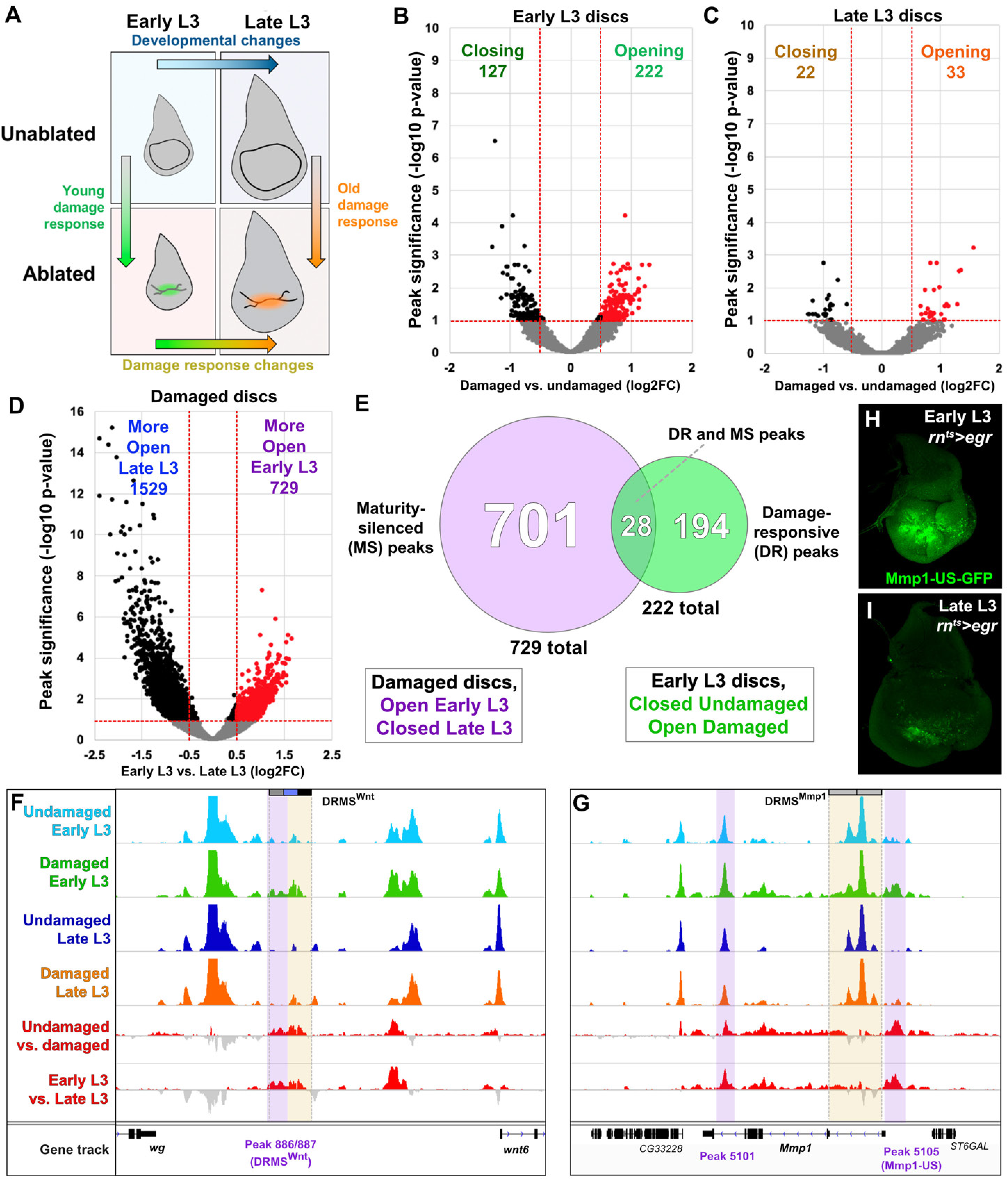
ATAC-seq of regenerating discs identifies damage-responsive and maturity-silenced regions, **(A)** Sample comparisons used in the ATAC-seq analysis, **(B-D)** Volcano plots identifying regions that are more or less accessible (opening or closing peaks) upon damage in early L3 discs (B), late L3 discs (C) or comparing damaged discs in early and late L3 (D). Lines indicate significance threshold (p<0.1) and fold change cutoff (log2FC>0.5). Red data points in (B) and (C) highlight regions opening upon damage at each stage. The number of peaks in each category is indicated on the graph, **(E)** Venn diagram showing genomic regions that are maturity silenced (MS, purple), damage-responsive (DR, green) and those that are both (DR and MS, intersection). MS peaks are defined as accessible (open) in damaged early L3 discs and less accessible (closed) in damaged late L3 discs. DR peaks are defined as closed in undamaged early L3 discs and open in damaged early L3 discs, **(F-G)** ATAC-seq chromatin accessibility traces (z scores) at the Wnt (*wg*/*Wnt6*) locus (F) and the *Mmp1* locus (G), for the four conditions indicated. Traces showing the difference between early L3 undamaged versus damaged discs, and early L3 damaged versus late L3 damaged discs, is shown in the bottom two traces (subtracted z scores, red and gray). As for other genomic traces, the peaks detected by computational analysis are indicated by purple boxes for MS peaks or green boxes for DR peaks, labelled with unique peak ID numbers underneath. The yellow boxes indicate the experimentally validated DRMS^Wnt^ (BRV118, Harris *et al*. 2016) (F) and the DRMS^Mmp1^ (characterized *Mmp1* enhancer of Figure 1) (G). Peaks 886/887 overlap the DRMS^Wnt^ enhancer, while peaks are not found in the DRMS^Mmp1^ enhancer, **(H-I)** Characterization of the *Mmp1-US* enhancer. Early L3 (H) and late L3 (I) *rn^ts^>egr* ablated discs bearing GFP reporter of the *Mmp1-US* MS region (peak 5105), showing damage responsive, maturity silenced behavior. Reporter expression in undamaged discs shown in Figure 1-Figure Supplement 2.

As our analysis was done using whole discs, any signal from the damaged portion of the disc is likely to be diluted by the chromatin profile of the rest of the disc. Therefore, we chose to analyze peaks that showed a log2FC>0.5 and an adjusted p<0.1. Using these cutoffs, our ATAC-seq data shows that 349 regions change upon damage in early L3 compared to undamaged controls, with 222 becoming more accessible and 127 becoming less accessible (Figure 2B and Supplemental table 1). In contrast, only 55 are differentially accessible upon damage in late L3 discs, with only 33 being more accessible and 22 being less so (Figure 2C and Supplemental table 2). Thus, the chromatin landscape, at least as assessed by this criterion, is more responsive to damage in immature (early L3) than in mature (late L3) discs, which in principle could contribute to the reduction in regenerative capacity. In this work we will focus on the regions that become more accessible upon damage.

We also looked for changes between damaged early L3 discs and damaged late L3 discs (Figure 2D and Supplemental table 3). We found 2258 differences: 729 regions become less accessible and 1529 regions become more accessible. The same comparison in undamaged discs from the two time points identifies 1638 regions becoming more accessible and 944 regions becoming less accessible (Figure 2 – Figure Supplement 1B and Supplemental table 4). Together these data imply that the chromatin landscape is remodeled more extensively by developmentally regulated signals than by damage. It is also possible that some damage-induced chromatin changes are obscured by the chromatin landscape of undamaged cells of the disc.

To identify possible DRMS enhancers we looked for regions that increase in accessibility upon damage in early L3 compared to undamaged discs (damage-responsive peaks, DR), and regions that are less accessible in late L3 damaged discs than early L3 damaged discs (maturity silenced, MS). Peaks that have both properties potentially represent novel DRMS enhancers (DR and MS peaks). Using these criteria, there are 222 DR peaks and 729 MS peaks, with 28 peaks present in both groups (Figure 2E and Supplemental tables 1, 3 and 5). However, DRMS enhancers could also already be accessible in early L3 (open even in the absence of damage), but are unused by damage signals such as JNK in the absence of injury. Such regions could fall within the remaining 701 MS peaks that represent chromatin that is already open in damaged early L3 discs and closed in discs damaged in late L3 (Figure 2E).

We examined whether the experimentally characterized DRMS enhancers at the *wg/Wnt6* and *Mmp1* loci might include peaks within either of these two categories. We found peaks in both loci that fall within the 729 MS peaks that are observed in damaged early L3 discs but not damaged late L3 discs (Figure 2F and G). These peaks were not in the set of 28 peaks that are also induced by damage. At the Wnt locus, two peaks that are in the MS category, peak 886 (log2FC=1.04, padj=0.02) and peak 887 (log2FC=1.17, padj=0.02), both map within the BRV118 (DRMS^Wnt^) fragment. The genomic traces of the Wnt locus suggest that these peaks may also be damage responsive (Figure 2F) consistent with activity of the *BRV118-GFP* reporter. Indeed, it has previously been shown that this region does increase in chromatin accessibility upon damage (Vizcaya-Molina et al., 2018). However, for both of these peaks the difference between undamaged and damaged in early L3 is insufficient to meet our statistical criteria for being classified as DR. Similarly, examination of the *Mmp1* locus shows that the chromatin at the characterized DRMS^Mmp1^ enhancer becomes more accessible upon damage in early L3 discs, but not in late L3, as assessed by a broadening of the peak (Figure 2G). However, a substantial proportion of this region also maintains significant accessibility regardless of damage or age, precluding its identification as a region of differential accessibility in either category (DR or MS) by our peak detection method.

Our analysis did, however, identify two MS peaks at the *Mmp1* locus: peak 5101 (log2FC=0.51, padj=0.07), which is found in a 3’ intron in *Mmp1* and peak 5105 (log2FC=0.98, padj=0.04), which is close to and upstream of the transcriptional start site. We generated a GFP reporter to the region spanning peak 5105 (*Mmp1-US-GFP*) and observed damage-responsive, maturity-silenced expression correlating with its reduced accessibility in mature discs indicated by ATAC-seq (Figure H-I). As for the *Mmp1-GFP* reporter, *Mmp1-US-GFP* showed no activity in the absence of damage (Figure 1 – Figure Supplement 2D). Again, this *bona fide* DRMS enhancer contained an MS peak but not a DR peak. The two regions identified by ATAC-seq that we have subsequently validated as DRMS enhancers so far, *wg/Wnt6* peaks 886/887 and *Mmp1* peak 5105, fall within the set of 729 MS peaks that show differential accessibility between early and late L3 discs but not among the 28 peaks that increase accessibility in response to damage. Thus, enhancers which respond to damage and are shut off with maturity can be readily detected using our criteria for maturity-dependent silencing, but the sensitivity of our method seems too low to define these regions as damage-responsive. This is possibly a result of using whole discs for the ATAC-seq, since non-blastema cells of the disc that have open chromatin at this locus could mask additional damage-specific accessibility changes that occur solely in regenerating cells. As such, it is possible that many more DRMS enhancers might be found in the set of 729 MS peaks even though those peaks do not survive our statistical criteria for being damage-responsive. Similarly it is conceivable, that some of the 222 DR peaks could be less obvious in older discs but not meet our statistical cutoffs.

### Characterizing DRMS enhancers and their regulatory targets genome-wide

We looked at the distribution of the 923 peaks that are DR and/or MS on a genome-wide basis (Figure 3A and Figure 3 – Figure Supplement 1A-B). Kharchenko *et al*. have previously catalogued the relative representation of different chromatin states in the mappable genome (Kharchenko et al., 2011). Relative to the mappable genome, the total open chromatin detected by ATAC-seq is relatively enriched for active promoters/transcriptional start sites, actively transcribed introns and underrepresented in actively transcribed exons. When the DR and/or MS peaks are compared to open chromatin, there is a strong reduction in the representation of promoters/transcriptional start sites and increased representation of actively transcribed introns and “other open chromatin” (Figure 3A and Figure 3 – Figure Supplement 1B). This is consistent with these peaks localizing to enhancers, which in *Drosophila* are typically found in intergenic regions or the first intron.

**Figure 3:**
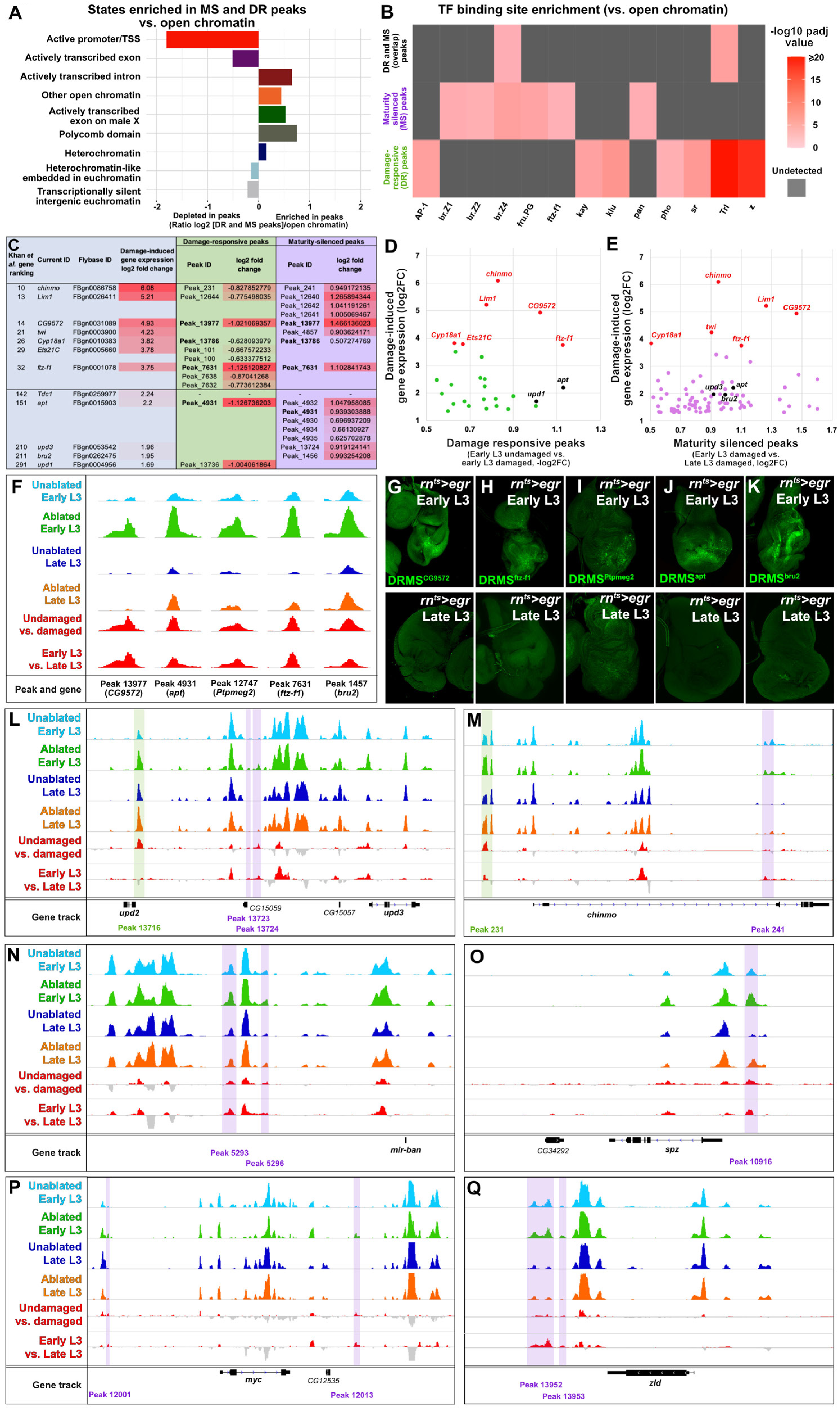
Genes expressed during regeneration are associated with DR and MS peaks. **(A)** Ratiometric graph showing the distribution of DR and MS peaks across the genome relative to total open chromatin detected by ATAC-seq, categorized into the chromatin states defined by ChIP-seq data of chromatin modifications from Kharchenko et al., 2011. DR and MS peaks detected in our ATAC-seq experiments are enriched at open chromatin found in introns and in other open chromatin, but depleted at promoters and exons, **(B)** Enrichment of candidate transcription factor binding sites in MS or DR peaks (full list of transcription factors tested is available in Supplemental methods table 2). Two motifs for AP-1 are tested, one based on validated sites identified in BRV118 (AP-1) and another using the Fly Factor Survey consensus for Kayak (Kay) as for the other transcription factors tested. Both sets of peaks have enrichment for several transcription factors previously associated with damage-responsive signaling. Adjusted P value for enrichment is indicated by strength of shading (capped at –log_10_ padj of 20). Gray indicates enrichment is not significant over background, **(C)** Table of genes shown to be upregulated in blastema cells (from Khan et al. 2017) that are associated with DR and/or MS peaks. Shown are the top 25 most strongly expressed genes (with measurable changes in gene expression, top 33 overall) during regeneration that have DR or MS peaks (top), and other genes with strong expression and significant DR or MS peaks described in this work (bottom). For each gene the fold change in gene expression is given (log2FC, Khan et al. data), and the type of peak (MS or DR) and peak identity associated with the gene is shown. The four Peak IDs in bold were among the 28 peaks found in both DR and MS groups, **(D-E)** Scatterplot showing the relationship between the change in chromatin accessibility of DR peaks (D) and MS peaks (E) versus damage-induced expression (Khan et al. data), showing that several of the most strongly expressed genes have the greatest increase in accessibility upon damage (D) and diminished accessibility with maturity (E). Highlighted data points are those in the top 25 upregulated genes (red), or other genes (black) shown in the table in (C), **(F)** ATAC-seq chromatin accessibility traces of the peaks (and associated genes) indicated, chosen for *in vivo* validation based on their strong DR and MS signatures and gene expression in blastema cells, **(G-K)** Early L3 (top) and late L3 (bottom) *rn^ts^>egr* ablated discs bearing GFP reporters of the peaks indicated in (F). Each reporter has damage-responsive expression in early L3 discs, which is reduced in damaged late L3 discs, consistent with chromatin accessibility of the region, **(L-Q)** ATAC-seq chromatin accessibility traces of at the loci of (L) the JAK/STAT ligands *upd2* and *upd3*, (M) *chinmo*, (N) the microRNA *bantam*, (O) the Toll pathway ligand *spz*, (P) the growth regulator *Myc*, and (Q) the transcription factor *zelda*, showing MS peaks (purple boxes) and DR peaks (green boxes), detected by differences in chromatin accessibility, as in Figure 2F-G.

Next we sought to identify DNA sequences found in the DR peaks and/or MS peaks that might imply regulation by specific DNA-binding proteins. We first assembled a list of 28 candidate transcription factors with known consensus sequences that have been identified in published studies of regeneration (listed in Supplemental methods table 2). A directed search in the DR and MS peak DNA sequences revealed enrichment of binding motifs in DR peaks, MS peaks and in the peaks that fall in both categories compared to the set of all 14,142 peaks identified in our analysis under all sets of conditions (Figure 3B). The DR peaks are most enriched for binding motifs for Zeste (Z) and Trithorax-like (Trl), also known as GAGA factor. Both Zeste and Trl function to regulate chromatin organization and promote gene expression (Mulholland et al., 2003; Bejarano and Busturia, 2004; Kostyuchenko et al., 2009). Also enriched are the binding motifs for the AP-1 (Kayak) transcription factor, Pho and two related zinc-finger proteins, Klumpfuss (Klu) and Stripe (Sr). The enrichment of AP-1 motifs is consistent with activation by the JNK pathway and the Pho motif suggests that a subset of these peaks could be targets of PcG-mediated silencing. Klu and Sr belong to a family of genes that are orthologs of the mammalian early growth response genes (EGR), which are known to function in wound healing and regeneration in different organisms (Wu et al., 2009; Lai et al., 2015). Recently, an EGR protein was shown to be a master regulator of whole-body regeneration in the acoel worm *Hofstenia miamia* through its role as a pioneer factor that modifies chromatin accessibility to regulate expression of many genes required for regeneration (Gehrke et al., 2019). The presence of binding sites for *Drosophila* orthologs of this factor in DR peaks suggests that some of the mechanisms that regulate chromatin accessibility during regeneration are conserved between *Hofstenia* and *Drosophila*.

The motifs enriched within the set of MS peaks are mostly non-overlapping with those in the DR peak group (Figure 3B). Notable is enrichment for the binding sites of three isoforms of the ecdysone-responsive transcription factor Broad which has previously been implicated in promoting a non-regenerative state in discs (Narbonne-Reveau and Maurange, 2019). Also enriched are binding motifs for an isoform of Fruitless and the nuclear receptor Ftz-F1. To our knowledge a role for these proteins in regulating regenerative capacity has not been demonstrated so far. Also enriched is the binding-motif for the Wnt-responsive transcription factor TCF/Pangolin indicating that, in addition to reduced damage-responsive *wg*/*Wnt6* expression, some targets downstream of Wnt signaling may also become less accessible in mature discs.

We identified the closest two genes for each of the 222 DR and 729 MS peaks, and the 28 peaks that are found in both sets (Supplemental tables 6-8), since the majority of known enhancers are thought to regulate their immediately flanking genes (Kvon et al., 2014). Gene ontology (GO) analysis using only the closest genes in the MS peak group shows that the most represented categories are those related to imaginal disc-derived wing morphogenesis and transcription (Supplemental table 9), suggesting these are likely candidates for genes regulating wing disc regeneration. By comparison, GO term analysis of the genes nearest the 222 DR peaks is primarily enriched only for gene transcription (Supplemental table 10).

We also compared our list of genes adjacent to both DR and MS peaks to previously published data of gene expression in regenerating discs. Khan *et al.* used a highly similar protocol, albeit using *rpr* rather than *egr* to ablate the wing pouch, and unlike other approaches, used sorted blastema cells to analyze damage-induced gene expression in L3 wing discs (Khan et al., 2017). Using this data set we found that, of the 502 most upregulated genes in regenerating discs, 85 (17%) have one or more associated MS peak and 27 (5%) have one or more DR peak, while 14 (3%) have both (Supplemental table 11). This includes 7 of the top 25 genes with measurable changes in gene expression, 5 of which have both MS and DR peaks (Figure 3C and Supplemental table 11). Several of the most strongly damage-induced genes are also associated with the most significant changes in both DR and MS peak accessibility (Figure 3D-E), showing that many of the regions with the greatest increases in chromatin accessibility upon damage, and the greatest loss of accessibility with maturity, are associated with genes that are most strongly upregulated upon damage. Furthermore, 19 of the 85 (22%) MS peak-associated genes and 5 of the 27 (19%) DR peak-associated genes from the Khan *et al*. dataset have more than one peak (Figure 3C), consistent with single genes being regulated by multiple DRMS enhancers, as we have found for *Mmp1*.

To directly assess whether regions containing the relevant peaks do indeed function as DRMS enhancers *in vivo*, we generated transgenic reporter lines by cloning DNA spanning the relevant peak upstream of a basal promoter driving GFP. We chose from peaks that are categorized as both DR and MS, and which are strongly upregulated in regenerating cells according to the Khan *et al.* dataset (Figure 3C, Supplemental table 11). Based on these criteria we tested regions adjacent to the genes *CG9572, ftz transcription factor 1* (*ftz-f1*), *Protein tyrosine phosphatase Meg2* (*Ptpmeg2*) and *apontic* (*apt*) all of which contain peaks that are in the overlapping set of 28 DR and MS peaks (Figure 3F and Supplemental table 8). We also tested a peak adjacent to *bruno 2* (*bru2*), even though this peak was only in the set of DR peaks, because the ATAC-seq traces strongly suggested some degree of maturity-dependent silencing, despite not meeting the statistical cutoff for an MS peak (Figure 3F), and because it is upregulated in blastema cells (Figure 3C). All five reporters were tested under both ablated and unablated conditions in early and late L3 discs, and each showed damage-induced expression in early L3 discs of varying strength (Figure 3G-K). Importantly this expression was consistently reduced in late L3 ablated discs (Figure 3G-K), and none of the regions tested showed activity during development in undamaged L3 discs (Figure 3 – Figure supplement 2A-E). Thus, the DNA corresponding to these peaks are indeed *bona fide* damage-responsive enhancers that are silenced with maturity, similar to the DRMS^Wnt^ and DRMS^Mmp1^ enhancers. More broadly, the damage-responsiveness of these transgenic reporters support the utility of chromatin accessibility profiling in the identification of cis-regulatory regions involved in shaping the transcriptional response to damage.

Having shown that this group of peaks is associated with genes known to be expressed in blastema cells, and can act as DRMS enhancers *in vivo*, we went on to characterize other potential regulatory targets. Functional annotation of the MS peak genes yields groupings that comprise pathways already implicated in regeneration, including Hippo, Hedgehog and Wnt signaling (Supplemental table 12). While it is known that enhancers are capable of regulating the expression of more distant genes, methods for predicting those interactions are still unreliable. The 1458 genes that are closest to the 729 MS peaks represent 1143 unique genes, with individual genes being associated with 1-8 peaks (Supplemental table 6). In addition to the Wnt pathway, the JAK/STAT pathway (Katsuyama et al., 2015; La Fortezza et al., 2016) and the Hippo pathway (Grusche et al., 2011; Sun and Irvine, 2011) have been shown to be required for regeneration in imaginal discs. Moreover, we have previously shown that the activity of each of these two pathways is markedly different in early and late L3 discs following damage (Harris et al., 2016). The genes encoding the ligands for the JAK/STAT pathway, *upd1*, *upd2* and *upd3* are clustered together on the X chromosome. We find three DR and two MS peaks in this locus, the peaks closest to *upd2* and *upd3* are shown in Figure 3L. There are also peaks close to three transcriptional targets of the JAK/STAT pathway, *chinmo*, *zfh2* and *Socs36E* (Supplemental table 6 and 7). The expression of *chinmo* correlates with regenerative capacity as it decreases in late L3 and increased expression of *chinmo* in late L3 has been shown to augment regeneration (Narbonne-Reveau and Maurange, 2019). We find a DR peak upstream of *chinmo* and an MS peak in the intron just upstream of the group of exons shared between the different transcripts (Figure 3M).

Expression of the microRNA *bantam* (*ban*) is activated by Yorkie, the transcriptional regulator downstream of the Hippo pathway. There are two MS peaks in the *ban* locus (Figure 3N). Interestingly, multiple upstream components of the Hippo pathway have MS peaks that are present in early L3 but not late L3 (*fat* (*ft*), 1 peak; *dachsous* (*ds*), 3 peaks, *dachs* (*d*), 1 peak; *four-jointed* (*fj*), 1 peak; *kibra*, 3 peaks; *Tao*, 1 peak, Supplemental table 6). *kibra* also has a DR peak (Supplemental table 7). Currently, there is little evidence that the transcription of these genes changes in response to regeneration. However, the observation that regions of chromatin adjacent to these genes are more accessible in early L3 than in late L3 could suggest that their transcription might be altered in response to damage in early L3 but less so in late L3.

The Toll ligand *spätzle* (*spz*) has an MS peak (Figure 3O) as do two other *spz*-related genes, *spz3* and *spz6.* Interestingly, several TLR-family genes, *Toll-4*, *Toll-7* and *Tollo* (2 peaks), also have MS peaks (Supplemental table 6), raising the possibility that this pathway might have a hitherto unsuspected role in regenerative growth. We have previously shown that the Myc protein is a key driver of regenerative growth (Smith-Bolton et al., 2009; Harris et al., 2016) and that Myc expression is reduced following ablation in late L3. We have detected two MS peaks near the *Myc* gene (Figure 3P). Finally, the pioneer transcription factor Zelda, is thought to function in concert with other transcriptional regulators to activate gene expression. There is a region 3’ to the *zelda* gene that becomes noticeably less accessible in mature discs (Figure 3Q). Although these peaks did not fall within the DR category using our statistical cutoffs, based on the traces shown in Figure 3L-Q, we note that the majority of these MS peaks show some level of induction following damage. In this respect, they are similar to the peaks at the Wnt and *Mmp1* loci that we have experimentally validated as DRMS enhancers but were not detected as DR peaks. Thus, a number of genes that are likely to function in regeneration, including components of pathways that are less active following damage in late L3, are near peaks that potentially indicate DRMS enhancers.

In summary, we have shown that genes known to be involved in regeneration as well as several genes not previously implicated as regulators of regeneration (*CG9572*, *apt*, *Ptpmeg2*, *ftz-f1* and *bru2*) are adjacent to experimentally-validated DRMS enhancers. To test whether specific genes are indeed necessary for regenerative growth, we have developed a GAL4-independent tissue ablation system - DUAL Control that enables us to manipulate gene activity during the phase of regenerative growth following ablation.

### A novel combinatorial expression system, DUAL Control, allows genetic manipulation of regenerating tissues

Proximity to DR or MS peaks could be a way of identifying novel genes that have an important role in tissue regeneration. It would then be necessary to test whether reducing the function of the gene would impact regeneration. The tissue ablation system that we have been using thus far causes cell death by rendering GAL4 active for 40 h at 30°C during L3, and thus activating expression of a pro-apoptotic gene under UAS control. Therefore, any other gene expressed under the control of UAS elements would also be expressed only during the time of ablation and be limited to the cells targeted for ablation. In order to take advantage of the extensive collections of UAS-driven RNAi lines and similar UAS-based tools available in *Drosophila* to study the genetic regulation of tissue regeneration, we developed a novel genetic ablation system that is independent of GAL4/UAS (Figure 4A), such that UAS-driven transgene can be expressed after the ablation phase, and importantly, in cells of the regeneration blastema (Figure 4B). Our system takes advantage of the bacterial transcriptional regulator LexA and its binding motif LexAOp, which have previously been used in *Drosophila* as an independent alternative to GAL4/UAS (Lai and Lee, 2006; Pfeiffer et al., 2010). To permit both temporal and spatial control over ablation, we first generated separate transgenes that express the DNA binding domain (DBD) of LexA under the control of a *spalt* (*salm*) enhancer (Jory et al., 2012) to provide spatial control, and a transcriptional activator p65 domain (Schmitz and Baeuerle, 1991) under the control of the *hsp70* heat shock promoter to provide temporal control (Figure 4A). Each domain bears a complementary leucine zipper (Ting et al., 2011), which allows formation of the full chimeric LexA::p65 only in cells that express both components. We combined these two transgenes with either *lexAOp-rpr* or *lexAOp-egr*, resulting in a system that allows tissue-specific ablation of the medial wing pouch in response to a heat shock, and which is entirely independent of GAL4/UAS. Since this system permits the simultaneous and independent use of both LexA and GAL4, each with separate spatial and temporal control, we have named this system <u>Du</u>ration <u>A</u>nd <u>L</u>ocation Control, or DUAL Control.

**Figure 4:**
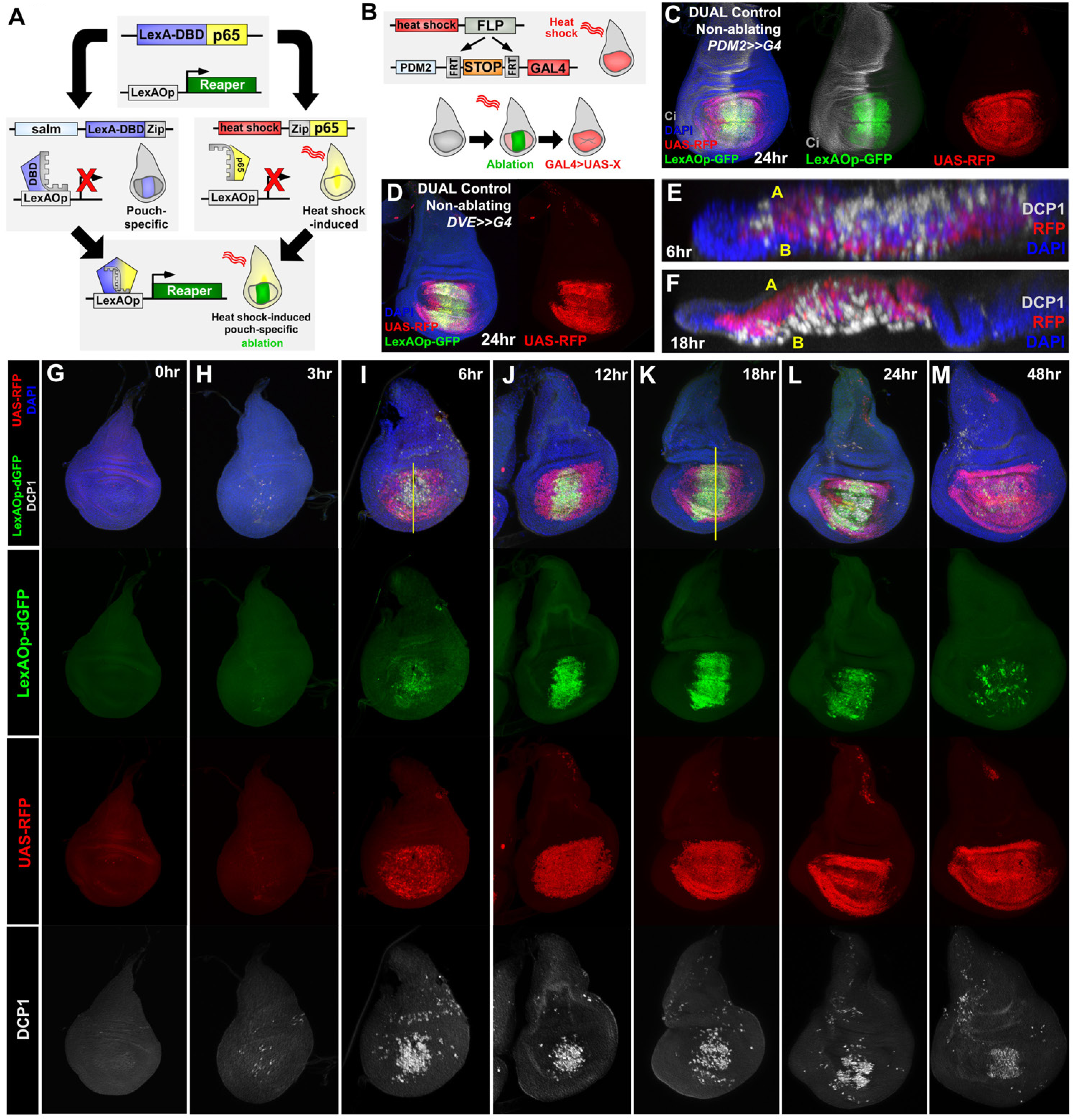
DUAL Control: a novel genetic ablation system to manipulate gene expression in regenerating discs, **(A)** Schematic of the DUAL Control genetic ablation system, based on split LexA. See manuscript text for details. *salm*: *spalt* enhancer, Zip: Leucine zipper domain, p65: transcriptional activator domain, **(B)** Schematic of the *PDM2>>GAL4* driver used in DUAL Control to manipulate gene expression in regenerating cells. Heat shock-induced FLP removes the transcriptional stop cassette through FRT recombination, allowing a single heat shock to activate both ablation (green) and GAL4 expression (red) in different cell populations, **(C)** Non-ablating DUAL Control with *PDM2>>GAL4* crossed to a double fluorescent tester stock, bearing *UAS-RFP* (red) and *lexAOp-GFP* (green), heat shocked at early L3 (day 3.5 at 25°C) and imaged 24 hr post heat shock (PHS). Cells that can be ablated (*salm* domain) are marked by GFP (green), while the surrounding cells that can be genetically manipulated (*PDM2* domain) are marked by RFP (red). The ablation domain straddles the compartment boundary, indicated by Ci staining (gray), **(D)** As in (C), using non-ablating DUAL Control with the stronger *DVE>>GAL4* driver. The expression of the *DVE* domain is similar to *PDM2*, **(E-F)** Sections through discs seen in (I) and (K), showing apoptotic cells (DCP1, gray) and pouch cells (RFP, red). The majority of dead cells and debris present within the disc proper at 6 hr PHS (E) is extruded basally from the disc epithelium by 18 hr PHS (F). A: Apical surface of the disc proper epithelium, B: Basal surface of the disc proper epithelium, DAPI, blue, **(G-M)** Time course of DUAL Control ablation with *rpr* on day 3.5 discs bearing *lexAOp-dGFP* (fast-degrading) and *UAS-RFP*, imaged at the indicated number of hours PHS. GFP (green), RFP (red) and cell death (DCP1, gray) can be detected at 6 hr PHS, and persist until 24 hr PHS. At 12 hr PHS the rate of new dead cell production decreases, while DCP1 positive cells and GFP label persist. At 24 hr PHS, GFP expression begins to decline and mostly subsides by 48 hr PHS. GAL4 expression (RFP) is consistent from 6 hr PHS to pupariation throughout the regeneration period. DAPI, blue. Yellow lines in (I) and (K) indicate cross-sections in (E) and (F).

We established a DUAL Control stock that also includes a pouch-specific GAL4 under the control of either a *PDM2* or *DVE* enhancer (Jory et al., 2012), both of which target expression to regions of the wing pouch surrounding the region that would be ablated. Although both enhancers drive in similar spatial patterns within the pouch (Figure 4C-D), our experiments show that *DVE* has consistently stronger expression (data not shown). These GAL4 lines were generated with a flip-out cassette to allow heat shock-induced FLP-mediated activation (*PDM2>>GAL4* or *DVE>>GAL4,* Figure 4B). Thus, a single heat shock can simultaneously induce LexA-driven ablation of the medial pouch and activate GAL4 expression in the surrounding cells (Figure 4C-D), which have been shown to generate the regenerated pouch (Herrera et al., 2013; Verghese and Su, 2016). A full description of the DUAL Control genotype can be found in Materials and Methods.

To characterize this new system, we crossed a non-ablating DUAL Control *>>DVE* stock to a double-fluorescent tester stock, *lexAOp-dGFP*; *UAS-RFP*, and heat shocked early L3 larvae (day 3.5 at 25°C). dGFP denotes a destabilized green fluorescent protein. We found that both the LexA component (*lexAOp*-driven ablation) and the GAL4 component (UAS-driven transgenes) of the system can be activated with a single 37°C heat shock of duration as short as 10 minutes. We tested different heat shock durations (Figure 4 - Figure Supplement 1A-D) and found that a single 45-minute heat shock was optimal, which was therefore used in all subsequent DUAL Control experiments. Using a time course, we examined the dynamics of LexA and GAL4 activity over time in discs ablated with *rpr* using DUAL Control *DVE>>GAL4* (Figure 4G-M). GFP, which is expressed in cells that also express *rpr*, can be detected within 6 hours post-heat shock (PHS, Figure 4I). At this time point the majority of the *salm* domain stains positively for activated caspase, which is mostly observed within the disc proper (Figure 4E). After 18 hours PHS the majority of cell corpses have been extruded basally from the disc (Figure 4F). At 24 hours the epithelium has mostly regained a normal appearance (Figure 4L), and by 48 hours PHS the associated debris is minimal, while regeneration is complete. (Figure 4M). Despite using a fast-degrading GFP reporter (dGFP) with a half-life of only a few hours (Lieber et al., 2011), we found that fluorescence induced by LexA::p65 persists in the disc after the initial heat shock, including in cells within the ablated *salm* domain (Figure 4J-L). This is likely due to these cells activating the *LexA-DBD* transgene as they take on distal pouch identity, which functions together with residual p65 to express *lexAOp-GFP*. These cells do not undergo apoptosis, however, indicated by the lack of new DCP1 staining, suggesting that this level of LexA::p65 activity is enough to drive GFP expression but not ablation.

By comparison, RFP expression resulting from heat shock induced *DVE>>GAL4* is observed at 6 hours PHS (Figure 4I) and is present in most cells of the wing pouch by 12 hours PHS (Figure 4J), persisting until pupariation. Finally, we observed that at 6 hours PHS, when the *salm* domain had undergone significant ablation, RFP was present in the surrounding cells, and persisted throughout the recovery period (Figure 4I-M), demonstrating that cells that drive the regenerative growth can be targeted for manipulation using DUAL Control.

### Reducing activity of individual DMRS-regulated genes using DUAL Control limits regeneration

Using DUAL Control we examined the effect of inducing ablation at different developmental time-points. Larvae were heat shocked at early and late L3 stages (days 3.5 and 4.5 at 25°C) and the extent of regeneration was measured by assaying the size of the resulting adult wings (Figure 5A). Ablation occurred comparably in discs at both developmental time-points using either *rpr* or *egr* as the pro-apoptotic stimulus, although ablation with *egr* resulted in lower levels of activated caspase (Figure 5 – Figure supplement 1A-D). The adult wings that develop from *rpr* ablated discs display a series of phenotypes, which we categorized into discrete groups: “wild type” for those indistinguishable from unablated wings (and therefore likely to be fully regenerated), “nicked” for those with some margin loss, “partial notch” or “full notch” for those with significant loss of both margin and wing blade tissue, and “ablated”, describing those that had lost the entire distal wing (Figure 5A). When ablated with *rpr* in early L3, around 95% of wings produced were in the “wild type” or “nicked” category, indicating that regeneration was mostly complete (Figure 5A). By comparison, ablation in late L3 yielded many more “full notch” or “ablated” wings (Figure 5A), confirming that imaginal discs lose the ability to regenerate in late L3. Ablation with *egr* resulted in weaker adult phenotypes (Figure 5 – Figure supplement 1E), consistent with the reduced level of caspase apparent in discs. We therefore used ablation with *rpr* for subsequent wing scoring assays.

**Figure 5:**
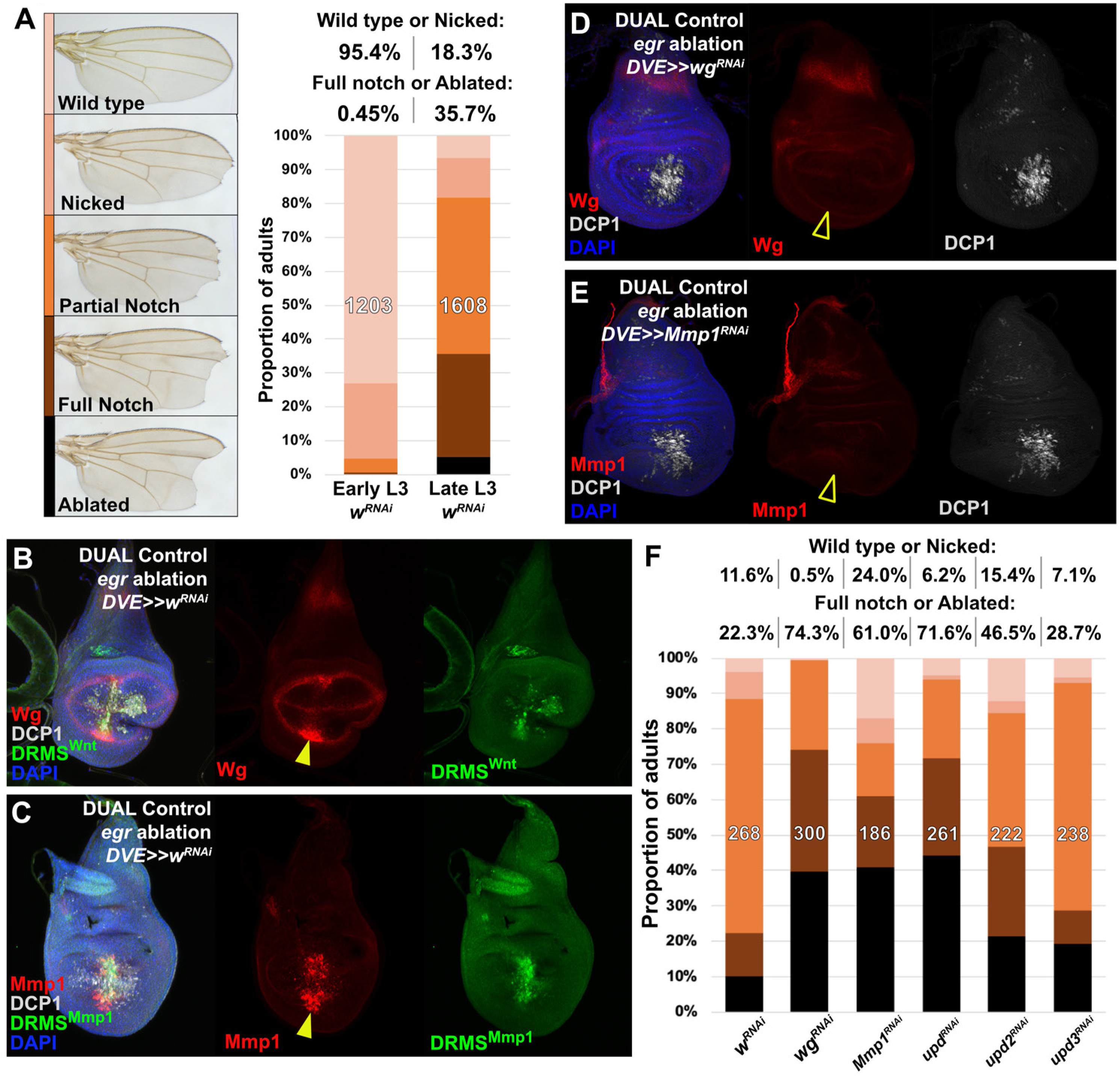
Manipulation of genes previously known to function during regeneration using DUAL Control, **(A)** DUAL control used to ablate discs with *rpr* in early (day 3.5) or late (day 4.5) L3 discs, assayed for regeneration by wing size. Classification of adult wing phenotypes following ablation and regeneration (left). Graphs illustrate the proportion of adults that eclose with the indicated wing phenotypes, showing the loss of regenerative capacity that occurs between early and late L3 (right). In this and subsequent figures, the number of flies scored is labelled on the graph of each genotype, and the percentage of each genotype characterized as “Wild type or Nicked” and “Full notched or Ablated” is indicated above. *UAS-w^RNAi^* is used as a control for regeneration scoring, **(B-C)** Early L3 discs bearing the DRMS^Wnt^-GFP reporter (B) or the DRMS^Mmp1^ GFP reporter (C), ablated with *egr* using DUAL Control and imaged 24 hr later. Ablation activates expression of both *wg* and *Mmp1* (red, arrowheads), and both DRMS reporters (green), **(D-E)** RNAi knockdown of damage-induced *wg* (D) and *Mmp1* (E) expression (red) in DUAL Control *egr* ablated discs. Note that developmental expression of *wg* in the hinge and notum, and *Mmp1* in the tracheal tubes is unaffected by the knockdown, which is limited to the regenerating pouch tissue (open arrowheads). DCP1: gray, DAPI: Blue, **(F)** RNAi knockdown of identified DRMS-target genes *wg*, *Mmp1*, and JAK/STAT ligands *upd1*, *upd2*, and *upd3* which have both DR and MS peaks in their vicinity in disc ablated with *rpr* using DUAL Control *DVE>>GAL4* in mid-L3 shows their requirement for regeneration, indicated by adult wing appearance.

We have previously identified multiple genes that are strongly expressed following ablation in early L3 but not in late L3 (Harris et al. 2016). Since regenerative capacity correlates with the expression of these genes, we could use DUAL Control to determine if expression of these genes in the tissue surrounding the ablated region was indeed necessary for regeneration. Therefore, we targeted for knockdown or overexpression, genes associated with MS and/or DR peaks: *wg*, *Mmp1*, the growth regulator *Myc*, the JAK/STAT ligands *upd2* and *upd3*. (Figure 2F-G, Figure 3L,P). *wg* and *Mmp1*, both of which have experimentally-validated DRMS enhancers, are detected following ablation with *rpr* (Figure 5 – Figure supplement 2A-B), and more so with *egr* due to its stronger activation of the JNK pathway (Figure 5B-C). The GFP reporter of each DRMS enhancer is also activated by DUAL Control ablation (Figure 5B-C). Targeting either gene for knockdown with RNAi using this system strongly decreases the damage-induced expression of both genes, although the amount of cell death appears unaffected (Figure 5D-E). The extent of regeneration, as assessed by the change in adult wing phenotype, is reduced (Figure 5F). These defects are dependent upon ablation and regeneration because knockdown of *Mmp1* for the same duration in the absence of ablation yields little to no effect on adult wings (Figure 5 – Figure supplement 2C), while knockdown of *wg* without ablation produces patterning defects localized to the distal wing edge (Figure 5 – Figure supplement 2D) that are clearly distinguishable from the wing tissue loss that follows ablation. Thus, *wg* is required for both regrowth following damage and repatterning. We also manipulated expression of *Myc*, a potent growth regulator shown to be sufficient to improve regeneration of late L3 discs (Smith-Bolton et al., 2009; Harris et al., 2016), and which is associated with two MS peaks (Figure 3P). Knockdown of *Myc* using RNAi directed to regenerating tissue with either DUAL Control *PDM>>GAL4* or *DVE>>GAL4* showed a dramatic reduction in regeneration (Figure 5 – Figure supplement 2E). Consistently, overexpression of *Myc* improves adult wing size and morphology (Figure 5 – Figure Supplement 2E). An E2F reporter (*PCNA-GFP*) shows that these phenotypes likely result from changes in damage-induced proliferation in response to altered levels of *Myc* (Figure 5 – Figure supplement 2F-I). JAK/STAT signaling is also an important regulator of regenerative growth, and we have identified MS peaks associated with *upd2* and *upd3* and DR peaks near *upd2* and *upd1* (Figure 3L). Knockdown of either of the JAK/STAT ligands *upd2* or *upd3* appears to reduce regeneration. However, knockdown of *upd1* appears to have the greatest effect (Figure 5F). Since the effects of the three *upd* genes could be additive on pathway function, we disrupted pathway components that are thought to function downstream of all three genes. Knockdown of other pathway elements including the transcription factor *Stat92E* and receptor *domeless* (*dome*) also results in less-complete wings (Figure 5 – Figure supplement 2J), confirming the likely requirement for JAK/STAT signaling in regenerating tissue surrounding the ablation domain. Together, these data show that several genes either associated with experimentally validated DRMS enhancers or with peaks identified in this study are functionally necessary for regeneration, and demonstrate that DUAL Control is a powerful tool to assay and manipulate gene function in regenerating tissue, circumventing the limitations of our previous ablation system.

### Apontic (Apt) limits regeneration via regulation of JAK/STAT signaling

Having established a robust assay to examine the effect of manipulating gene activity on regeneration following ablation, we explored the potential for identifying novel regulators of regeneration by investigating genes that were in close proximity to DRMS enhancers. We chose to investigate two genes – *apontic* (*apt*) and *CG9572*, and have not characterized *ftz-f1*, *Ptpmeg2* or *bru2* further. Five MS peaks identified in our screen, of which one is also a DR peak (peak 4931), are found within an intron of *apt* (Figure 6A). We have shown that DNA spanning peak 4931 does indeed function as a DRMS enhancer *in vivo* (Figure 3J). Additionally, based on the data of Khan *et al*., it is transcriptionally upregulated in blastema cells (Figure 3C and Supplemental table 11). *apt* encodes a b-Zip transcription factor (Eulenberg and Schuh, 1997; Gellon et al., 1997). It was originally identified as a dominant enhancer of the semi-lethality of heteroalleleic combinations of *Deformed* (*Dfd*) alleles; *Dfd* encodes a homeobox-containing protein that patterns portion of the maxillary and mandibular segments (Gellon et al., 1997). Independently, *apt* (also known as *trachea defective*, *tdf*) was shown to be necessary for the initial invagination and migration of cells in the embryonic tracheal placode (Eulenberg and Schuh, 1997). Subsequently, *apt* was shown to limit the number of anterior follicle cells in the ovary that are specified as migrating border cells (Starz-Gaiano et al., 2008). In border cell selection and in the male germline, decreasing *apt* function results in increased expression of JAK/STAT targets, suggesting *apt* functions, at least in this context, to restrict JAK/STAT signaling (Starz-Gaiano et al., 2008; Monahan and Starz-Gaiano, 2016), which is known to be required for wing disc regeneration (Katsuyama et al., 2015) (Figure 5F and Figure 5 – Figure supplement 2J). More recently, *apt* has been shown to promote expression of *cyclin E* and *hedgehog* in imaginal discs (Liu et al., 2014; Wang et al., 2017). Using an antibody that recognizes the Apt protein (Liu et al., 2003), we found that in unablated discs Apt is expressed at high levels in the cells of the squamous cells of the peripodial epithelium and tracheal tubes of developing wing discs throughout L3, and expression is not obviously above background in cells of the disc proper (Figure 6 – Figure supplement 1A-B). However, optical section imaging of ablated discs shows that Apt can be detected in the disc proper following damage in early L3 (Figure 6B), but not in late L3 (Figure 6C). Thus, the presence DR and MS peaks within the *apt* gene is consistent with its stage-specific damage-responsive expression.

**Figure 6:**
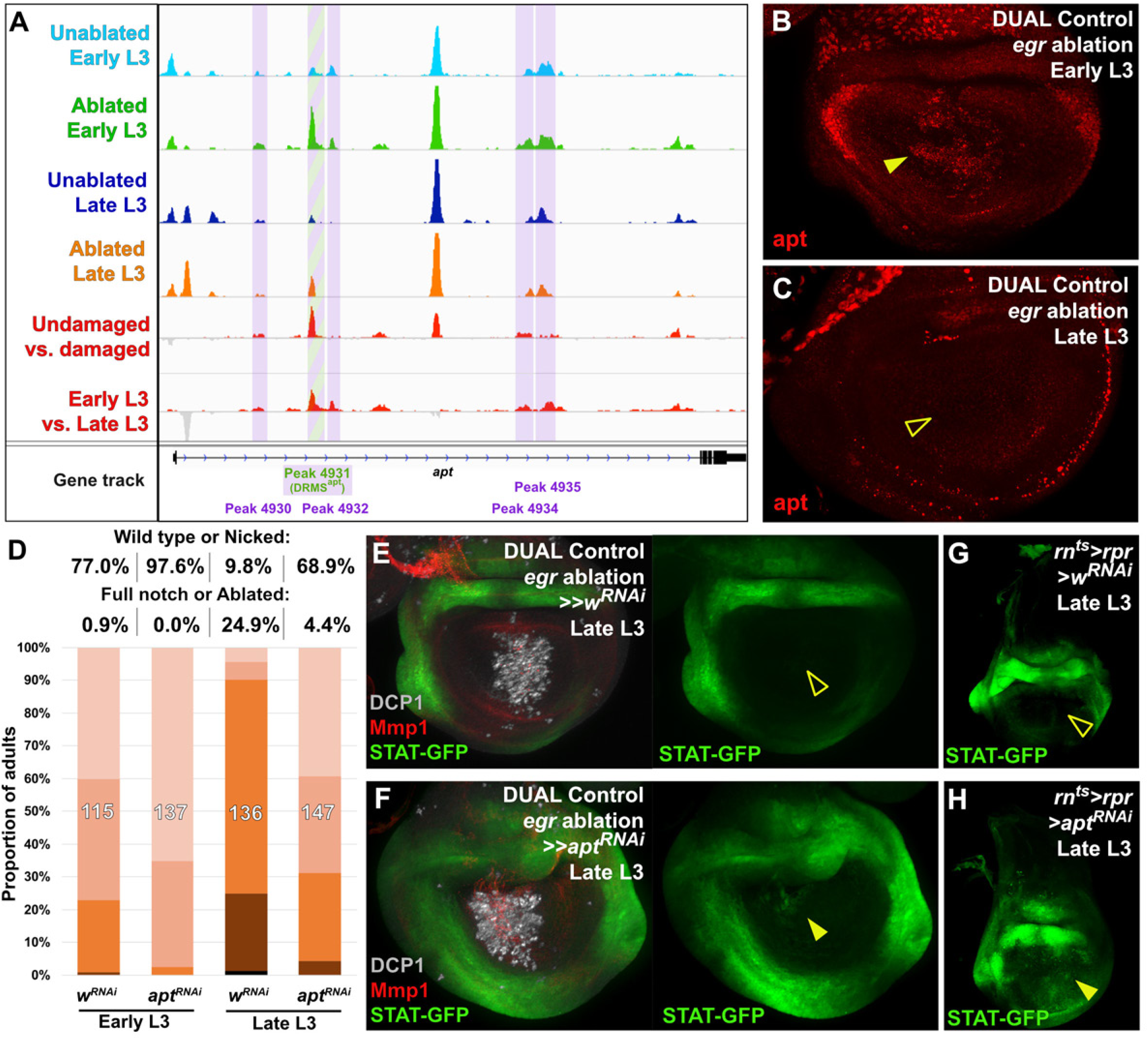
DRMS enhancer-regulated gene *apontic* (*apt*) regulates regeneration by modulating JAK/STAT signaling, **(A)** ATAC-seq chromatin accessibility traces at the *apt* locus, showing the four detected MS peaks (purple boxes) and single peak that is both DR and MS (green and purple box) – the DRMS^apt^ enhancer was experimentally validated (see Figure 3J), **(B-C)** Apt protein (red) detected in the disc proper in early L3 (B) and late L3 (C) discs ablated with *egr* using DUAL Control. *Apt* is expressed in blastema cells upon ablation in early L3 discs (arrowhead), but is absent in late L3 (open arrowhead). Developmental expression of *apt* persists in cells of the peripodial epithelium and trachea at both time points, **(D)** RNAi knockdown of damage-induced *apt* with DUAL Control *DVE>>GAL4* discs ablated with *rpr* increases regeneration in both early and late L3 discs, as assessed by wing size, **(E-H)** Late L3 discs bearing a *Stat92E* reporter (*STAT-GFP)* ablated with DUAL Control *DVE>>G4* using *egr* (E-F) or *rn^ts^>rpr* (G-H) in the presence of *apt* RNAi (F and H) or control RNAi (E and G). Ablation in control samples shows no increase in reporter activity in the regenerating cells of late L3 discs (E and G, open arrowheads), while knockdown of *apt* in the pouch results in increased Stat92E reporter expression in cells surrounding the zone in the medial disc where ablation occurs using DUAL Control (F, arrowhead), or in the majority of the pouch when ablated with *rn^ts^*, which targets the entire pouch (H, arrowhead). GFP expression from earlier developmental expression persists in the hinge region of all samples. DCP1: gray, Mmp1:red.

To test whether *apt* is necessary for regeneration, we reduced its expression using DUAL Control by expressing an *apt* RNAi transgene in regeneration-competent early L3 discs. In view of the demonstration that *apt* can promote wing growth by promoting Hedgehog expression (Wang et al., 2017), we were surprised to find that the adult wings were more complete than in ablated controls not expressing the *apt* RNAi (Figure 6D). We then examined the effect of *apt* knockdown in late L3 discs and found that, once again, adult wings were more complete than in controls (Figure 6D). These observations indicate that *apt* knockdown either protects discs from damage during the ablation phase or promotes regeneration. Since the levels of cleaved caspase DCP1 appeared similar in the presence and absence of *apt* knockdown (Figure 6E-F), this suggests that *apt* normally acts to limit regeneration following damage. As *apt* is known to act in the germline to limit JAK/STAT signaling, we examined whether it might affect regeneration by influencing JAK/STAT activity. We observed the expression of a fluorescent Stat92E reporter (*STAT-GFP*) in late L3 discs ablated with DUAL Control in the presence of *apt* knockdown. Normally at this stage, STAT activity is minimal in regenerating cells of ablated discs (Figure 6E). However, with *apt* knockdown, we found an increase in the activity of this reporter specifically in blastema cells (Figure 6F). This is more clearly visible with *rn^ts^>egr* ablation, which damages a larger area of the disc (Figure 6G-H). These data suggest that rather than being necessary for regeneration, activation of *apt* expression in young, but not old discs, might be a mechanism of tempering the extent of regeneration when it occurs.

### *asperous* (*CG9572*) is a novel regulator of regenerative capacity

The uncharacterized gene *CG9572* is one of the most strongly upregulated by damage in regenerating cells (Figure 3C) and has a peak just upstream of the transcriptional start site that is both damage-induced and maturity silenced (Figure 7A). Moreover, using a reporter gene, we have shown that the DNA spanning this peak functions as a DRMS enhancer (Figure 3G). *CG9572* is predicted to encode a 441 amino acid protein of unknown function. In order to better characterize CG9572, we performed protein blast (blastp) and alignment scoring, which revealed strong sequence similarity to the Jagged protein (Figure 7 - Figure supplement 1A), a membrane bound ligand for Notch in vertebrates (Lindsell et al., 1995). A Jagged ortholog in *Drosophila* has not been described. The peptide sequence is predicted to contain seven EGF-type repeats that each displays a characteristic spacing of cysteine residues (Figure 7 - Figure supplement 1B-C). EGF-like repeats are found in all Notch ligands that have been described to date (reviewed by (Kovall et al., 2017)). Similar to Jagged, CG9572 also has a 14-amino acid hydrophobic stretch close to its N-terminus that is likely to function as a signal peptide (Figure 7 - Figure supplement 1D). However, unlike Jagged or other Notch ligands it lacks a second hydrophobic stretch that would serve as a transmembrane domain (Figure 7 – Figure supplement 1D). Thus, at least based on its sequence, the predicted CG9572 protein has similarity to a Jagged-like Notch ligand, but unlike Jagged or other known Notch ligands, is likely to be secreted. Due its similarity to mammalian Jagged, we have called this protein Asperous (Aspr). When a tagged version, UAS-*aspr::HA*, was expressed in the posterior compartment using *en-GAL4*, HA staining appeared to be cytoplasmic (Figure 7B), localizing towards the apical surface of the disc proper (Figure 7C) and with occasional punctae observed in the anterior compartment. When the stronger *hh-GAL4* driver was used, punctae were observed throughout the anterior compartment consistent with the Aspr protein being secreted from cells (Figure 7D). Overexpression of *aspr* in the whole wing pouch using *rn-GAL4* does not result in an observable phenotype in the adult wing (Figure 7 – Figure supplement 2A), but expression in the posterior compartment causes abnormal folding in the pouch at the boundary with wild type cells (Figure 7 – Figure supplement 2C).

**Figure 7:**
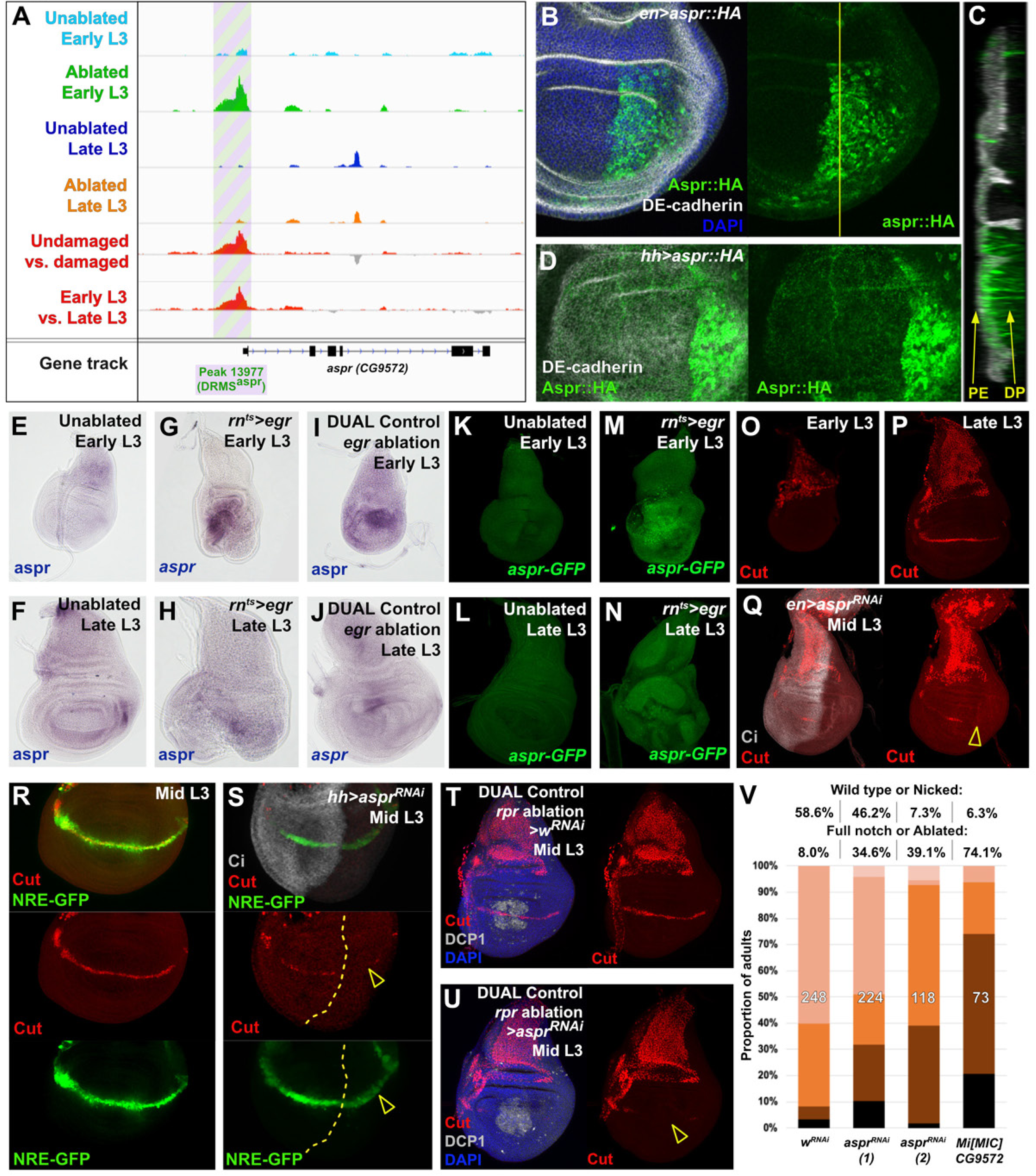
Uncharacterized gene *CG9572*/*asperous* is a novel regulator of wing disc regeneration. **(A)** ATAC-seq chromatin accessibility traces at the *asperous* (*aspr)* locus, showing a peak that is both DR and MS and was experimentally validated as a DRMS enhancer (Figure 3G)(DRMS^aspr^, green and purple box), **(B)** Ectopic expression of an epitope tagged Aspr (Aspr::HA, green) in the posterior compartment using *en-GAL4*. The protein is cytoplasmic and is mostly excluded from nuclei. DE-cadherin (gray). DAPI: blue. Yellow line indicates plane of image in (C), **(C)** Z-section though the disc shown in (B), showing mostly apical localization of Aspr::HA in the expressing cells (green), and presence of Aspr::HA between the disc proper (DP) and peripodial epithelium (PE), suggesting extracellular localization. DE cadherin shows cell membranes (gray), **(D)** High magnification imaging of an apical disc section of a disc expressing Aspr::HA (green) under the control of *hh-GAL4*, showing punctae of staining away from the expressing cells at a level between the peripodial membrane and the disc proper epithelium, **(E-F)** RNA *in situ* hybridization detects weak *aspr* expression in the ventral and lateral areas of the disc in early L3 (E) and late L3 (F) discs, but is mostly absent from the pouch, **(G-J)** RNA *in situ* hybridization detecting *aspr* in early L3 (G and I) or late L3 (H and J) discs following ablation with *egr* using *rn^ts^>* (G-H) or DUAL Control (I-J). In both cases, *aspr* is upregulated in blastema cells in the pouch upon ablation in early L3 discs, but only weakly in late L3, **(K-N)** Expression of an *aspr* GFP MiMIC reporter line (green) in early L3 (K and M) and late L3 discs (L and N) during normal development (K-L) and following ablation with *rn^ts^>egr* (M-N). Consistent with the RNA *in situ* expression data, *aspr* reporter activity is mostly absent during normal development, being activated by damage in early L3 discs, with reduced activity in late L3 discs, **(O-P)** Expression of *cut* during normal development in early L3 (O) and late L3 (P) wing discs. Cut protein (red) is detected in cells of the notum, including myoblasts and trachea, at both developmental time points, and becomes upregulated at the D-V boundary in late L3 in response to Notch signaling, **(Q)** Knockdown of *aspr* in the posterior compartment with *en-GAL4* driving *aspr* RNAi delays onset of *cut* expression, as shown by a lack of Cut at mid L3 (open arrowhead) when it usually extends across the entire posterior compartment. Ci: Gray, **(R-S)** Cut protein (red) and a Notch reporter *NRE-GFP* (green) in mid-L3 wing discs, showing expression of both extending across the entire D-V boundary in wild type discs (R), while discs expressing *aspr* RNAi in the posterior compartment using *hh-GAL4* have delayed expression of Cut and weaker NRE activity (S, open arrowheads). Ci (gray) demarcates A-P compartment boundary (yellow dotted line), **(T-U)** Mid L3 discs ablated with *rpr* using DUAL Control *DVE>>GAL4* and driving a control RNAi (T) or *aspr* RNAi (U). Cut protein (red) is quickly reestablished following *rpr* ablation, extending across the D-V boundary (T). Knockdown of *aspr* in the entire pouch with *DVE>>GAL4* prevents reestablishment of Cut during regeneration (U, open arrowhead). DCP1: gray, DAPI: blue, **(V)** Knockdown of *aspr* using two different RNAi lines, or heterozygosity for a presumed *aspr* mutant (*Mi[MIC]CG9572*), in discs ablated with *rpr* using DUAL Control *DVE>>GAL4* reduces regeneration, as assessed by wing size.

As the annotated transcriptional start site of *aspr* is close to a peak that is both MS and DR (Figure 7A), we examined its expression following damage in discs of different maturity using RNA *in situ* hybridization, and a gene-trap cassette insertion line (Mi[MIC]CG9572[^MI02471^]), which bears an eGFP gene that can be used to monitor *aspr* expression (Venken et al., 2011). In undamaged L3 wing discs, *in situ* hybridization shows *aspr* is expressed at low levels in the ventral and lateral areas of the disc at low levels (Figure E-F). Similarly, the gene trap shows little to no expression (Figure 7K-L). Upon damage in early L3 discs, *aspr* is upregulated strongly in the region of the blastema, as shown in discs ablated by both *rn^ts^>egr* (Figure 7G,M) and DUAL Control ablation with *egr* (Figure 7I). In damaged discs from late L3 larvae, *aspr* has much weaker damage-induced expression (Figure 7H,J,N). Knockdown of *aspr* with two different RNAi lines in the developing wing pouch using *rn-GAL4* in undamaged discs has little effect on adult wing size or patterning (Figure 7 – Figure supplement 2B and data not shown). However, we found that knockdown of *aspr* in mid L3 discs using *en-GAL4* delays the onset of expression of the Notch target *cut* at the prospective wing margin in the posterior compartment (Figure 7O-Q), suggesting *aspr* might promote Notch signaling during normal development. This is also shown by the weak reduction in fluorescence of a Notch reporter, NRE-GFP (Zacharioudaki and Bray, 2014) (Figure 7R-S). These data suggest that Aspr is a secreted regulator of Notch signaling in the wing, which is strongly activated in regenerating tissue upon damage. To address whether *aspr* is necessary for regeneration, we used DUAL Control to reduce its expression in regenerating cells in mid-L3 discs using the two different RNAi lines following *rpr* ablation of the wing pouch. At this stage Cut expression at the margin is beginning to be established (Figure 7R), while ablation at an earlier stage leads to a delay in development and hinders Cut expression. Upon *aspr* knockdown the presence of Cut in these discs is markedly reduced (Figure 7T-U), suggesting that ablation combined with the loss of *aspr* may limit Cut expression that has already been initiated. The extent of regeneration is also decreased (Figure 7V). This effect on regeneration was also observed with *rn^ts^>egr* ablation (Figure 7 – Figure supplement 2D), where expression in the regenerating tissue is likely to be less than achieved with DUAL Control since the RNAi is expressed in the same cells as *egr* during the ablation phase. These experiments suggest that *aspr* promotes specification of the wing margin during development. We also tested the MiMIC line that we used as a GFP reporter for *aspr* expression, which is likely to also be an *aspr* mutant due to the mutagenic cassette in the insertion that is designed to disrupt gene expression (Venken et al., 2011). The insertion is in the first intron of the coding region, downstream of the transcriptional start site of all three aspr transcripts (Figure 7 – Figure supplement 2E). Semi-quantititive PCR of *rn^ts^>egr* ablated discs from MiMIC hemizygous animals (Mi[MIC]CG9572[^MI02471^]/Y) showed strongly reduced levels of *aspr* mRNA compared to ablated wild type discs (Figure 7 – Figure supplement 2F), consistent with this line being a transcriptional mutant. *aspr* hemizygous animals have no obvious developmental defects (data not shown) but show a strongly decreased ability to regenerate when ablated with DUAL Control (Figure 7V) and *rn^ts^>egr* (Figure 7 – Figure supplement 2G). Expression of Aspr::HA does not strongly affect regeneration (Figure 7 – Figure supplement 2H), indicating that while it is necessary for regeneration, ectopic expression alone it is not sufficient to improve it, as we have demonstrated with other pro-regeneration genes such as *wg*. Thus, we have used proximity to a DRMS enhancer to identify a novel gene with sequence similarity to Notch ligands as a potential regulator of regenerative capacity.

### Using DUAL Control to screen for chromatin modifiers that influence regeneration

The presence of DRMS enhancers in the vicinity of multiple genes that each contribute to regeneration suggests that restoring expression of a single gene would be insufficient to restore regenerative capacity in mature discs. Indeed, we have previously shown little benefit in restoring *wg* expression alone in mature discs (Harris et al. 2016). One way to restore expression of multiple genes regulated by DRMS elements in late L3 discs would be to manipulate levels of chromatin silencing factors that are involved in inactivating these damage-responsive enhancers at multiple loci. Since reducing levels of these factors throughout the organism could have pleiotropic effects, we therefore used DUAL Control to knock down a panel of epigenetic silencing factors in regenerating tissue of late L3 discs. Most of those tested did not have an obvious effect (Figure 8A). However, inhibition of the Polycomb group gene *extra sex combs* (*esc*), which encodes a component of the PRC2 (reviewed by (Kassis et al., 2017), specifically during the regeneration period using DUAL Control consistently improved regeneration, even in discs ablated in early L3 (Figure 8B). We used the weaker *PDM2*-driven version of DUAL Control for these experiments to avoid the pleotropic effects and lethality associated with strong or widespread knockdown of epigenetic regulators. Under these conditions, we observed no adult wing defects when *esc* was knocked down in unablated wings for this short duration (data not shown). Thus, even a small reduction in *esc* levels is likely to improve regeneration.

**Figure 8:**
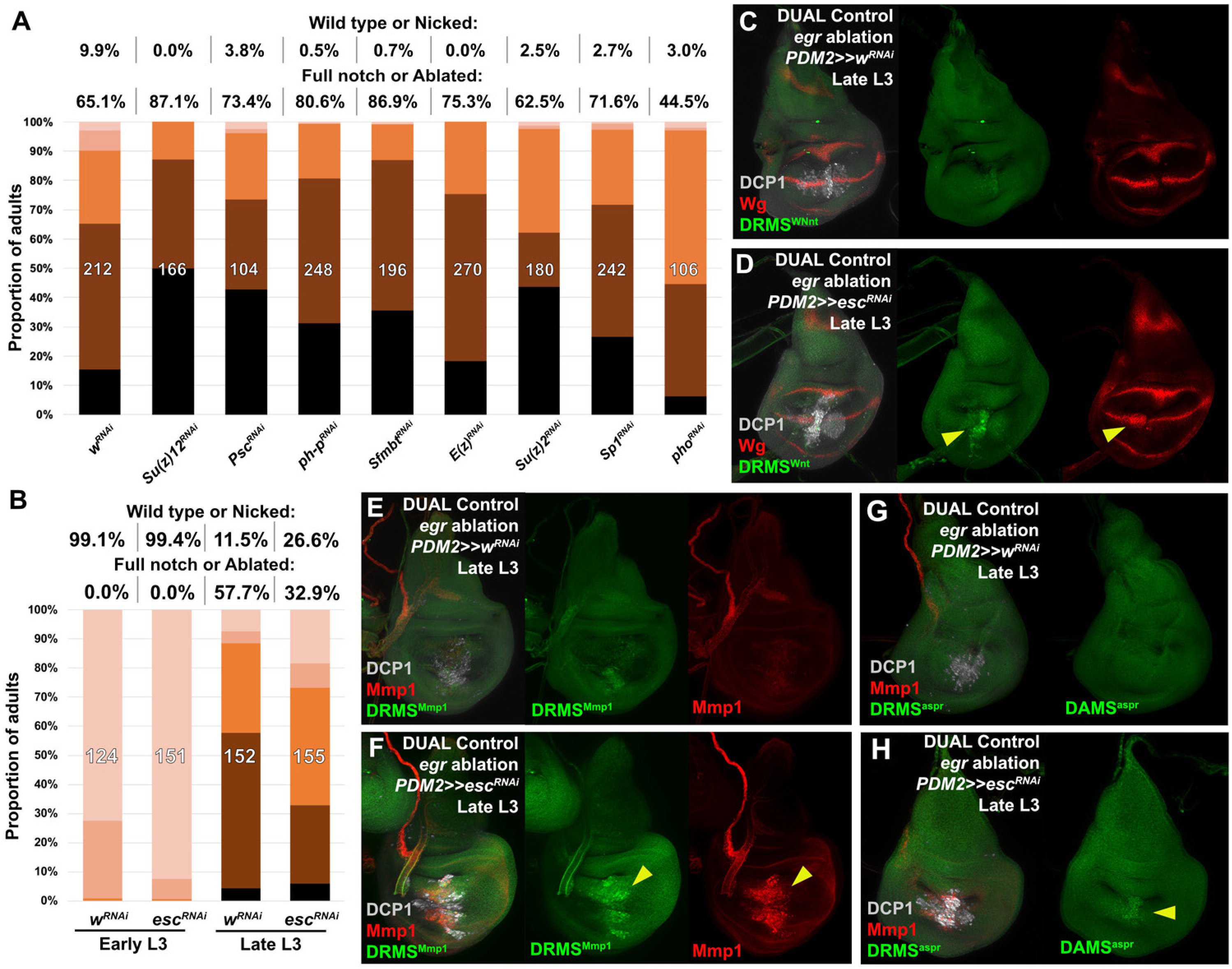
A screen of chromatin silencing factors shows *esc* limits regenerative capacity by silencing DRMS enhancers in mature discs, **(A)** Knockdown of various epigenetic silencing factors in late L3 *rpr* ablated discs using DUAL Control *PDM2>>GAL4*. The loss of most factors negatively impacts regeneration compared to the *w^RNAi^* control, assayed by wing size, **(B)** Knockdown of *esc* as in (A) improves regeneration of both early and late L3 discs, **(C-F)** Late L3 discs bearing DRMS^Wnt^ (C-D) and DRMS^Mmp1^ (E-F) GFP reporters (green) ablated with DUAL Control using *egr*. Control discs (C and E) show little damage-induced reporter activity, or expression of *wg or Mmp1* (red). By comparison, knockdown of *esc* by *PDM2>>GAL4* increases damage-induced reporter expression (D and F arrowheads), as well as *wg* and *Mmp1*. DCP1: gray, **(G-H)** Discs as in (C-F), showing knockdown of *esc* in late L3 discs increases reporter activity of the DRMS^aspr^ enhancer (green).

To test whether the improved regenerative capacity of *esc* knockdown was due to alteration of DRMS activity, we examined the activity of three different DRMS reporters, DRMS^Wnt^, DRMS^Mmp1^, and DRMS^aspr^, in late discs expressing *esc^RNA^*^i^. The level of GFP expression from all of the reporters was increased in ablated discs with *esc* knockdown compared to controls, as was *wg* and *Mmp1* expression (Figure 8C-H), indicating that the loss of *esc* leads to improved regeneration in part via reactivation of multiple DRMS enhancers. Thus, reducing Polycomb-mediated silencing appears capable of overcoming the suppression of a regeneration program that operates simultaneously at multiple loci in the genome.

## Discussion

*Drosophila* imaginal discs lose the ability to regenerate as they progress through the third larval instar (Smith-Bolton et al., 2009; Harris et al., 2016). This correlates with reduced damage-induced expression of a number of genes that have been shown to be important for regeneration. We previously showed that the damage-responsive expression of both *wg* and *Wnt6* requires an enhancer that lies between the two genes (Harris et al., 2016). This enhancer consists of two separable modules: a damage-responsive module and an adjacent module that promotes silencing of the entire enhancer. Importantly, this silencing, as assessed by H3K27 trimethylation, does not spread either to developmentally-regulated enhancers at this locus or to the coding regions of *wg* and *Wnt6*. This mechanism blocks damage-responsive expression of *wg* in mature discs without inactivating *wg* function entirely and suggests a way in which gene expression necessary for regeneration could be blocked without compromising the use of those same genes for developmentally-regulated growth and patterning.

We also found that restoring *wg* expression alone in mature discs was insufficient to preserve regenerative capacity at this stage of development, suggesting that genes other than *wg* that are required for regeneration could be regulated in a similar way. We have now shown that a number of such genes are indeed regulated by a similar mechanism. First, by looking for conserved sequence motifs in DRMS^Wnt^, we identified a comparable enhancer at the *Mmp1* locus. This enhancer has a similar bipartite composition, and its damage-induced activity declines as discs develop through L3. Subsequently, using ATAC-seq to measure chromatin accessibility genome-wide, we found that many genes encoding components of pathways known to function in regeneration, often extracellular ligands or their receptors, are adjacent to regions with localized chromatin changes. We categorized these changes as damage-responsive (DR, opening upon ablation) or maturity silenced (MS, reduced accessibility in old damaged discs versus young damaged) or both, and found that discrete regions bearing these chromatin signatures are associated with genes encoding components of the Wnt, Hippo, JAK/STAT and Toll pathways. Furthermore, when compared to expression data of sorted blastema cells (Khan et al., 2017), we found that both DR and MS peaks are associated with several genes that are upregulated during regeneration. However, the number of regions we identified using ATAC-seq are likely to be an underestimate since our assay examined chromatin changes using whole discs, in which only a small proportion of cells are undergoing regeneration. Thus, the chromatin status of developing cells away from the blastema could potentially mask damage-specific changes, which we showed is likely the case at the *Mmp1* locus. For this reason we expect the number of DR and MS peaks to be higher than we have shown here, and that additional regions of interest might be uncovered with the recent development of single cell ATAC-seq approaches. However, taken together, these observations indicate the presence of a transcriptional program that is likely to be activated in immature discs by damage-responsive enhancers, which becomes muted in more mature discs by widespread silencing of these regulatory elements.

Analysis of both groups of DR and MS regions showed that each is enriched for particular transcription factor binding motifs, suggesting coordinated activation and silencing by distinct pathways that regulate both behaviors in regenerating tissue. These findings parallel those of other groups who have used chromatin profiling in imaginal discs to show that coordinated epigenetic regulation of genomic enhancers is an important mechanism that regulates various behaviors of imaginal discs, including their growth and identity during development (McKay and Lieb, 2013; Ma et al., 2019). In a recent study that investigated the basis of terminal exit from the cell cycle in cells of the *Drosophila* wing disc (Ma et al., 2019) it was found that distal enhancers for key cell cycle regulators such as *cyclin E* and *string* became less accessible as development proceeds. This occurs independently of cell-cycle status, but seems instead to be governed by a temporal program. Cell-cycle exit becomes more and more robust as development proceeds as assessed by the inability of ectopic expression of cell cycle regulators to promote re-entry into the cell cycle.

### Identification of *asperous* as a novel regulator of regeneration

By its proximity to a DRMS enhancer, we identified *aspr* as a gene necessary for full regeneration. While knockdown of *aspr* has no obvious function during normal development, it adversely impacts regenerative growth. Establishment of the wing margin is an early event in the formation of the wing pouch. One view of wing development proposes that the pouch is generated by a wave of recruitment emanating from the margin; undifferentiated cells at the edge of the growing pouch are recruited to express *vestigial*, a marker of the pouch fate (Zecca and Struhl, 2007). We have shown that following *aspr* knockdown, expression of the markers at the margin such as Cut and the Notch reporter is delayed both in normal and regenerating discs suggesting that *aspr* could potentially promote Notch signaling. Our sequence analysis reveals Aspr has sequence similarity to Notch ligands, and is most similar to the vertebrate Notch ligand Jagged (Lindsell et al., 1995), suggesting its influence on Notch signaling could be as a ligand. However, unlike all Notch ligands characterized to date, Aspr lacks a predicted transmembrane domain and is therefore likely to be secreted. Consistent with this notion, we observed punctate staining outside the region of expression when *aspr::HA* was overexpressed. Being anchored to the membrane is thought to be crucial for the mechanism by which Notch ligands function; endocytosis of the ligand engaged to Notch is presumed to generate the mechanical force that alters Notch conformation and renders it susceptible to proteolytic cleavage, which eventually results in nuclear translocation of the intracellular domain (Parks et al., 2000; Langridge and Struhl, 2017). Thus, at least based on this view of Notch signaling, it is unlikely that Aspr could function as a canonical Notch ligand. Moreover, Notch signaling is thought to function as part of double repressive mechanism that controls regenerative growth (Smith-Bolton et al., 2009), and is reduced in blastema cells to permit *Myc*-dependent growth. As such, it is unclear how Aspr might promote Notch signaling, and yet also be required for regeneration. Importantly, overexpression of *aspr* did not appear to increase Notch signaling, at least as assessed by activation of the Notch reporter. Therefore the role of Aspr in promoting regeneration could be independent of a direct influence on Notch signaling, and remains to be explored.

### Identifying manipulations to promote regeneration in mature discs

Since multiple genes necessary for regeneration appear to be silenced as larvae progress through L3, it seems unlikely that restoring the expression of any single factor can restore regenerative capacity. We were surprised to find that knockdown of *apt* appears to improve regeneration in discs ablated in either early or late L3. Knockdown of *apt* increased expression of a STAT reporter, suggesting this improvement might be via increased JAK/STAT signaling. However, we cannot exclude the possibility that other mechanisms might be more important. A previous study has shown that reducing JAK/STAT signaling reduced tissue loss following *egr*-induced ablation, but did not seem to compromise the extent of compensatory proliferation (La Fortezza et al., 2016). Furthermore, the overexpression of *upd* or *upd2* concurrently with ablation did not improve regeneration. However, increasing Stat92E activity in the blastema during the regeneration phase was not tested in that study, as appears to occur when *apt* is knocked down using DUAL Control. It is possible that modulation of Stat92E activity might affect more or different gene targets than the *upd* ligands alone to promote regeneration.

We also found that reducing PcG-mediated repression via knockdown of *esc* improved overall regeneration. Reducing *esc* activity derepressed at least three DRMS enhancers that were tested using reporter gene constructs. Since a reduction in *esc* levels alleviates silencing at multiple DRMS enhancers, it is likely that many of these other enhancers are also regulated by Polycomb-mediated silencing, as for DRMS^Wnt^. However, this knock down is not sufficient to completely restore regeneration to that of early L3 discs. We previously showed that the silencing of the DRMS^Wnt^ enhancer is associated with highly localized accumulation of H3K27 trimethylation at the enhancer locus. Interestingly, disruption of H3K27 or H3K9 trimethylation at cell-cycle enhancers is insufficient to prevent cell-cycle exit in the wing disc during the pupal phase (Ma and Buttitta, 2017). This suggests that other, as yet uncharacterized, mechanisms might function in addition to heterochromatin formation and PcG-mediated silencing to irreversibly make these enhancers inaccessible to transcription factor binding. By analogy, reducing *esc* function may only be able to extend regenerative capacity until these other mechanisms reduce accessibility of DRMS elements, or limit gene expression via other means. Genome-wide screens, potentially involving tools such as DUAL Control, have the potential to uncover some of these mechanisms.

## Materials and Methods

### Fly stocks and genotypes

Stocks and crosses were maintained on yeast food at 25°C, except those for GAL4/UAS based ablation experiments, which were maintained at 18°C. Stocks used in this study: *rn^ts^>egr* (*w1118*;; *rn-GAL4*, *tub-Gal80ts*, *UAS-egr*) and *rn^ts^>rpr/egr* (*w1118*;; *rn-GAL4*, *tub-Gal80ts*, *UAS-rpr* or *UAS-egr*), *rn^ts^>* (*w1118*;; *rn-GAL4*, *tub-Gal80ts*)(Smith-Bolton et al., 2009), *AP-1-RFP* (Chatterjee and Bohmann, 2012), *UAS-his::RFP* (Emery et al., 2005), *UAS-dGFP* and *lexAOp-dGFP* (Lieber et al., 2011), *UAS-dILP8* (Colombani et al., 2012), *DRMS^Wnt^-GFP* (*BRV118-GFP*, (Harris et al., 2016)), *PCNA-GFP* (Thacker et al., 2003), *Hh-GAL4* (Tanimoto et al., 2000), NRE-GFP (Zacharioudaki and Bray, 2014), *UAS-aspr^SIRNAi(M1)^* (labelled as (2) in manuscript) and *UAS-aspr::HA* (a generous gift from David Bilder). Stocks obtained from the Bloomington stock center: *UAS-hep^CA^* (BL6406), *hep^r75^* (BL6761), *pc^15^* (BL24468), *lexAOp-hrGFP* (BL29954), *10xStat92E-GFP* (BL26197) *UAS-w^RNAi^* (BL33613), *UAS-myc* (BL9674), *UAS-myc^RNAi^* (BL51454), *UAS-hep^RNAi^* (BL28710), *UAS-bsk^RNAi^* (BL53310), *UAS-upd^RNAi^* (BL33680), *UAS-upd2^RNAi^* (BL33949), *UAS-upd3^RNAi^* (BL32859), *UAS-dome^RNAi^* (BL53890), *UAS-stat92E^RNAi^* (BL33637), *UAS-apt^RNAi^* (BL26236), *UAS-wg^RNAi^* (BL32994), *UAS-mmp1^RNAi^* (BL31489), *UAS-su(z)2^RNAi^* (BL57466), *UAS-e(pc)^RNAi^* (BL67921), *UAS-e(z)^RNAi^* (BL33659), *UAS-Sfmbt^RNAi^* (BL32473), *UAS-ph-p^RNAi^* (BL33669), *UAS-pc^RNAi^* (BL33622), *UAS-psc^RNAi^* (BL38261), *UAS-pho^RNAi^* (BL42926), *UAS-su(z)12^RNAi^* (BL33402), *UAS-esc^RNAi^* (BL31618) *UAS-Sp1^RNAi^* (BL35777), *UAS-CG9572/aspr^RNAi^* (labelled as (1) in manuscript) (BL58340), *en-GAL4* (BL30564), *Mi*[*MIC*]*CG9572*[*MI02471*] (BL 35863).

### Ablation experiments

GAL4/UAS-based genetic ablation experiments and wing scoring were performed essentially as described in Smith-Bolton et al. (Smith-Bolton et al., 2009), with each experimental condition compared to a suitable control that was ablated and scored in parallel. Unless otherwise indicated, discs were dissected and fixed for immunofluorescence immediately after the ablation period. DUAL Control ablation experiments were also density controlled (50 larvae per vial) and experiments conducted at 25°C, with a 37°C heat shock administered at day 3.5 or day 4.5 in a circulating water bath for 45 minutes unless otherwise stated. A detailed ablation protocol is available upon request. Discs were dissected, fixed and stained at 24 hr PHS unless otherwise stated. Wing scoring experiments were performed on the number of flies per genotype shown, resulting from at least two distinct biological replicates. Physical wounding experiments were performed essentially as described for *ex vivo* culture in Harris et al. (Harris et al., 2016).

### DUAL Control stock generation

The DUAL Control stock genotype is *hsFLP; hs-p65::zip, lexAOp-*ablation */ CyO; salm-zip::LexA-DBD, PDM2 or DVE>>GAL4*. A non-ablating stock was generated without a *lexAop*-ablation transgene. The ablation drivers used were *lexAOp-rpr* and *lexAOp-egr.* Each was generated by replacing the *GFP* coding sequence from *pJFRC19-13XLexAop2-IVS-myr::GFP* (Pfeiffer et al., 2010) with the full *rpr* or *egr* coding sequence from genomic DNA and LP03784 respectively. The resulting transgenes were inserted into landing site *su(Hw)attP5* (BL32231) via PhiC31 recombination. The *hs-p65* construct was built by cloning nucleotides −242 to 0 upstream of the TSS from the *Hsp70* gene into *pAttB* along with the *p65AD::zip* and *Hsp70* 3’UTR from *pBPp65ADZpUw* (Pfeiffer et al., 2010). The transgene was inserted into landing site *attP40* (BL25709) and recombined with *lexAOp-rpr* or *lexAOp-egr*. The *LexA-DBD* was generated by removing the *GAL4-DBD* sequence from *pActPL-zip::GAL4-DBD* (Luan et al., 2006), and replacing it with the codon optimized *LexA-DBD* from *pBPLexA::p65Uw* (Pfeiffer et al., 2010). This *zip::LexA-DBD* cassette was then cloned into *pAttB* along with the *salm* enhancer fragment R85E08 (Flylight), the DSCP sequence (Pfeiffer et al., 2008) and *Hsp70* 3’UTR. This transgene was inserted into landing site *attP2* (BL8622). The *PDM2>>GAL4* and *DVE>>GAL4* constructs were generated by cloning the *PDM2* or *DVE* enhancer fragments R11F02 or R42A07 (Flylight) into *pAttB*, along with an *FRT-PolyA-FRT* cassette, the *GAL4* coding sequence (GenBank: NM_001184062) and *SV40* 3’UTR. The transgene was inserted into landing site VK00027 (BL9744) and recombined with *salm-zip::LexA-DBD*. Both recombined chromosomes were built into a single stock with hsFLP (BL8862) on the X chromosome. Detailed plasmid maps are available on request.

### Transgenic reporter line construction

The *Mmp1-GFP* enhancer reporter was generated by amplifying the *Mmp1* genomic region using primers listed in Supplemental Methods Table 1, and cloning upstream of a minimal *hsp70* promoter and *eGFP* coding sequence into *pAttB* (accession KC896839.1). Related *Mmp1* GFP reporters were generated by replacing the *Mmp1* enhancer DNA with genomic regions amplified from genomic DNA with the primers listed in Supplemental Methods Table 1, as were reporters for the DRMS^ftz-f1^, DRMS^Ptpmeg2^, DRMS^bru2^, DRMS^aspr/CG9572^, and DRMS^apt^ regions. All GFP reporters were inserted into the *AttP40* landing site via PhiC31 recombination ensuring comparability. Transgenic services were provided by BestGene (Chino Hills, CA).

### Immunofluorescence and *in situ* hybridization

Discs were fixed and stained for immunofluorescence essentially as in (Harris et al., 2016), and mounted in ProLong Gold Antifade Reagent (Cell Signaling, Beverly, MA). The following primary antibodies were used in this study: from the DSHB, Iowa City, IA; mouse anti-Wg (1:100, 4D4), mouse anti-Mmp1 (1:100, a combination of 14A3D2, 3A6B4 and 5H7B11), rat anti-Ci (1:10, 2A1). anti-Cut (1:100, 2B10). Other antibodies; rabbit anti-DCP1 (1:1000, Cell Signaling), rabbit anti-TDF/apt (1:1000, (Liu et al., 2003), rabbit anti-HA (1:1000, Cell Signaling). Secondary antibodies used were from Cell Signaling, all at 1:500; donkey anti-mouse 555, donkey anti-rabbit 555, donkey anti-rat 647, donkey anti-rabbit 647, donkey anti-rabbit 488 and donkey anti-mouse 488. Nuclear staining was by DAPI (1:1000, Cell Signaling). Samples were imaged on a Leica TCS SP5 Scanning Confocal, Zeiss LSM 700 Scanning Confocal or Zeiss M2 Imager with Apotome. RNA in situ hybridizations were performed according to established methods for alkaline phosphatase-based dig-labelled probe detection. Discs were dissected and fixed as for immunofluorescence, Digoxigenin labelled probes were generated targeting the *aspr* gene coding sequence using the primer pairs listed in Supplemental methods table 1 to generate templates with T7 sequences at either the 5’ (sense probe) or 3’ (anti-sense probe) ends. Control and experimental discs were stained simultaneously for the same duration, mounted in Permount (Fisher Scientific, Pittsburg, PA) and imaged on a Zeiss Axio Imager M2.

### Semiquantitative PCR

Around 50 discs were dissected from equivalently staged male larvae of genotypes *rn^ts^>egr* (ablated), *rn^ts^>* (unablated), *rn-GAL4/UAS-aspr:HA* or *Mi(MIC)CG9572/Y;; rn^ts^>egr* (ablated) at early L3 (day 7, 18°C), and added to equal volumes of Trizol (Sigma). RNA was extracted according to established protocols, yielding 2-4μg total RNA per sample. cDNA was synthesized using Denville rAmp™ cDNA Synthesis Kit (Thomas Scientific) from 200ng starting RNA of each sample with a polyT primer. Subsequently, 1ul of each cDNA library was used for reduced-cycle semi-quantitative PCR with the primers listed in Supplemental methods table 1 to detect *aspr* and *actin* expression. The entire assay was repeated using discs from a separate ablation experiment for confirmation.

### ATAC-seq and Sequencing analysis

Samples for ATAC-seq library preparation were generated as follows: Larvae of genotype + *; +; rn-GAL4, GAL80ts* (*rn^ts^>*, undamaged) and + *; +; rn-GAL4, GAL80ts, UAS-egr* (*rn^ts^>egr*, damaged) were grown to early L3 (day 7) or late L3 (day 9) and upshifted to 30°C for 40 h, as for other ablation experiments. Discs were dissected in PBS immediately upon downshift and collected as pools of 100 discs for early L3 samples and 50 discs for late L3 samples. The 4 samples were placed in lysis buffer (10mM Tris 7.5, 10mM NaCl, 3mM MgCl2, 0.1% IGEPAL CA-630), pelleted, and exposed to the Tn5 transposase enzyme (Illumina) essentially as in Buenrostro *et al.* (Buenrostro et al., 2013). Three biological repeats were performed, and DNA was sequenced on a HiSeq2500 or HiSeq4000 as single index, multiplexed samples with 50PE or 100PE reads. Reads were trimmed to 50 bp using cutadapt (Martin, 2011) and then aligned to the dm3 reference genome using bowtie 2 (setting: --seed 123, -q –X 2000). Reads with quality scores below 5 were removed. Reads mapping to Chr2L/2R/3L/3R/4/X were employed in subsequent analysis. Peaks were called with MACS2 (setting: -g dm –keep-dup all –shift 9 –nomodel –seed 123), using a sonicated genomic DNA dataset as a control (Zhang et al., 2008). Signal tracks were generated from individual replicates using Deeptools v2.4.1 (Ramirez et al., 2016). Signal in browser shots are represented as z scores of pooled replicates as described previously (Uyehara et al., 2017).

### Differential accessibility analysis

ATAC peak calls from pooled datasets were ranked by MACS2 q-score. The top 11,500 peaks from each dataset were selected and combined into a union peak set of 46,000 peaks which was subsequently reduced by merging any peaks that overlapped by 1 bp or more using the GenomicRanges Bioconductor package (Gentleman et al., 2004; Lawrence et al., 2013). This resulted in a final union peak set of 14,142 peaks. ATAC-seq reads were counted inside union peaks using featureCounts from Rsubread (setting: isPairedEnd = T, requireBothEndsMapped = T, countChimericFragments = F, allowMultiOverlap = T) (Liao et al., 2013). The resulting count matrix was used as input for DESeq2 analysis (Love et al., 2014). Peaks were called as differentially accessible using the criteria of log2FoldChange>0.5 and padj<0.1.

### *In silico* sequence analysis and motif scanning

Comparison of DRMS DNA sequences and identification of the 17bp motif was performed using BLAST (https://blast.ncbi.nlm.nih.gov/) and Gene Palette software (Rebeiz and Posakony, 2004). Analysis of the 17bp motif was performed using Meme Suite with the TOMTOM Motif Comparison Tool (Gupta et al., 2007), which was run using default to compare the motif against the combined *Drosophila* databases. For motif scanning, 28 transcription factors with a previously-associated role in the regeneration response were used for directed motif enrichment analysis (Supplemental methods table 2). DNA sequences from the full length DR and MS peaks were scanned using AME version 5.1.0 (setting: --scoring fisher –hit-lo-fraction 0.25 –evalue-report-threshold 10) from the MEME suite using the union set of 14,142 ATAC-seq peaks as background (McLeay and Bailey, 2010). PWMs were obtained from Fly Factor Survey (Zhu et al., 2011), listed in Supplemental methods file 1. DNA sequences were extracted using the Biostrings and dm3.Org.Db Bioconductor packages (Pagès H, Aboyoun P, Gentleman R, DebRoy S, Biostrings: Efficient manipulation of biological strings. R package version 2.42.1., Carlson M (2019). org.Dm.eg.db: Genome wide annotation for Fly. R package version 3.4.0.). In cases where multiple PWMs for the same transcription factor were detected, the p-value corresponding the best PWM match was reported.

## Acknowledgements

The authors would like to thank David Bilder for his gift of unpublished stocks. We thank current members of the Hariharan and Harris labs for useful feedback, especially Melanie Worley, Jack StPeter and Jacob Klemm. We thank the Bloomington Stock Center, Drosophila Genomics Resource Center, Developmental Studies Hybridoma Bank, and BestGene for stocks, reagents, and services. This work was funded by NIH grant R35 GM122490 and a Research Professor Award from the American Cancer Society (RP-16238–06-COUN) to IKH, and by NIH grant R35 GM128851 and a Research Scholar Award from the American Cancer Society (RSG-17-164-01-DDC) to DJM.

## Funding Information

This paper was supported by the following grants:

National Institute of General Medical Sciences R35 GM122490 to Iswar K Hariharan. American Cancer Society RP-16-238-06-COUN to Iswar K Hariharan.

National Institute of General Medical Sciences R35 GM128851 to Daniel J. McKay. American Cancer Society RSG-17-164-01-DDC to Daniel J. McKay.

---

**Figure 1 – Figure supplement 1:**
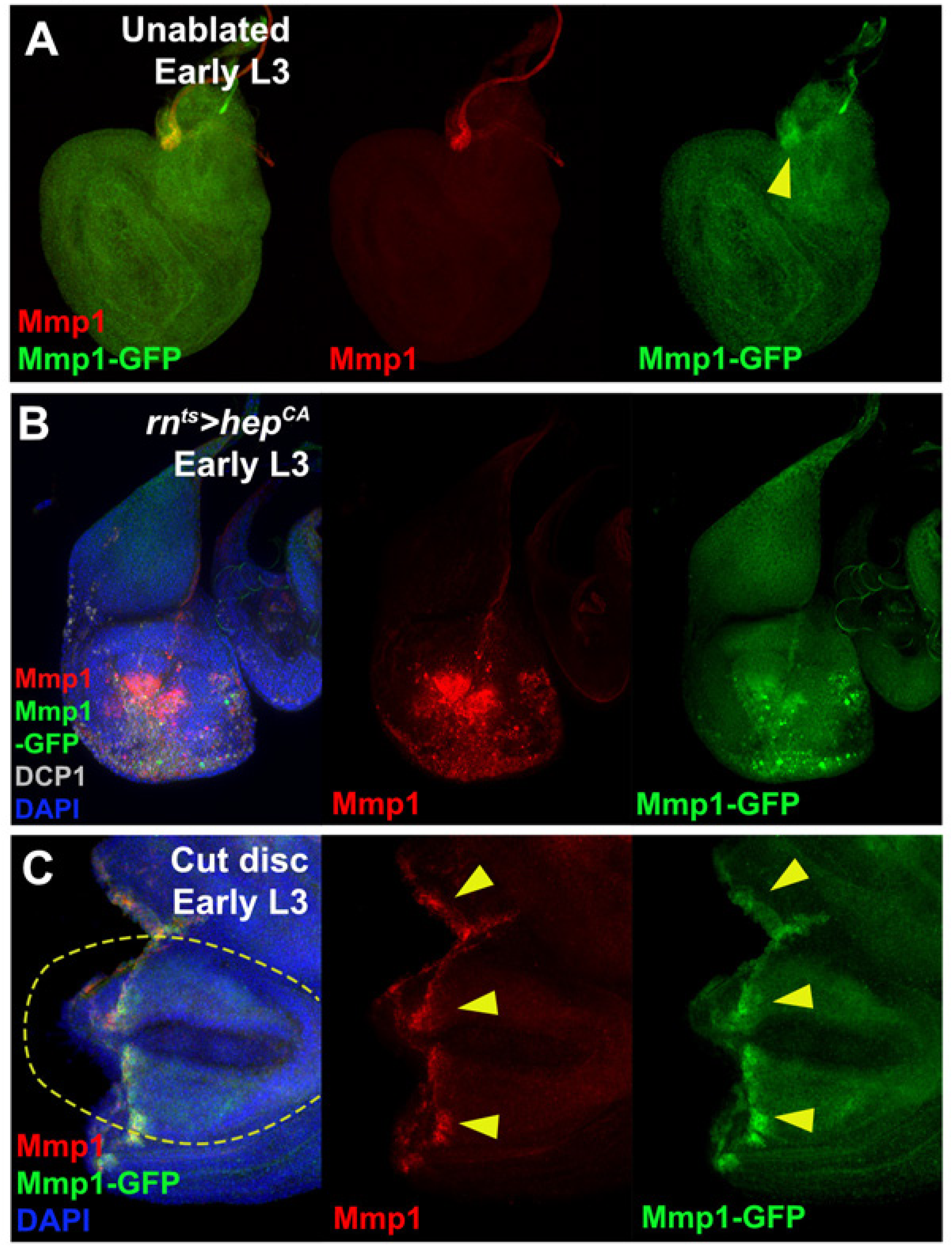
The *Mmp1* enhancer is activated by both ectopic JNK signaling and physical wounding, **(A)** An early L3 disc bearing the *Mmp1-GFP* enhancer reporter (green), showing the reporter weakly recapitulates developmental Mmp1 (red) in the air sac primordium (arrowhead), **(B)** Early L3 disc with ectopic activation of JNK signaling in the pouch via expression of *UAS-hepCA* with *rn^ts^>*. Both Mmp1 (red) and the enhancer reporter (green) are strongly activated. Cleaved caspase DCP1: gray, DAPI: blue, **(C)** Early L3 disc bearing the *Mmp1-GFP* enhancer reporter (green), physically fragmented, cultured for 8 hours and stained for Mmp1 (red). Both GFP and Mmp1 are coincident in cells along the wound edge. Original wing pouch is indicated with yellow dotted line, cut edge is indicated with yellow arrowheads DAPI: blue.

**Figure 1 – Figure supplement 2:**
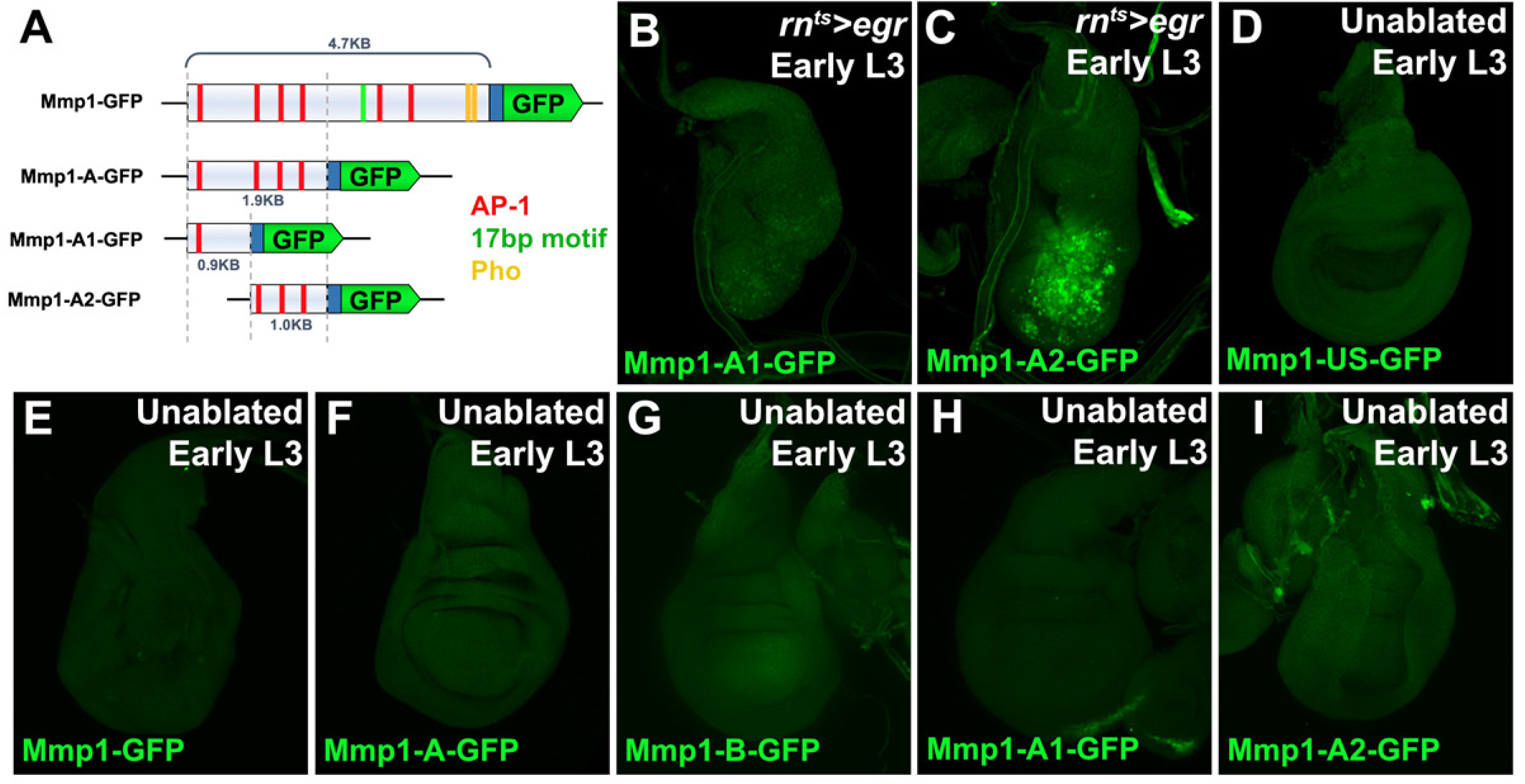
The damage-activated portion of the *Mmp1* enhancer is defined by a 1kb fragment, **(A)** Schematic of the *Mmp1-GFP* reporter and related constructs used to investigate damage-activation of the enhancer. AP-1 binding sites (TGASTMA, red bars), Pho binding sites (AATGGCB, yellow bars) and the 17bp motif (TGGCGATCGGCGGGAGT, green bars) are indicated. Blue box: *hsp70* minimal promoter, **(B-C)** Early L3 discs bearing the *Mmp1-A1-GFP* (B) and *Mmp1-A2-GFP* reporter (C) showing GFP expression (green) following ablation with *egr*. The 1kb fragment of the enhancer with 3 consensus AP-1 binding sites is responsible for the majority of damage-induced activity, **(D-I)** Early L3 undamaged discs bearing the GFP reporters indicated. None of the reporters expresses GFP (green) in developing discs in the absence of ablation, including the *Mmp1-US* reporter (*Mmp1-US* described in Figure 3) (D) or subdivisions of the *Mmp1* enhancer (E-I).

**Figure 2 – Figure supplement 1:**
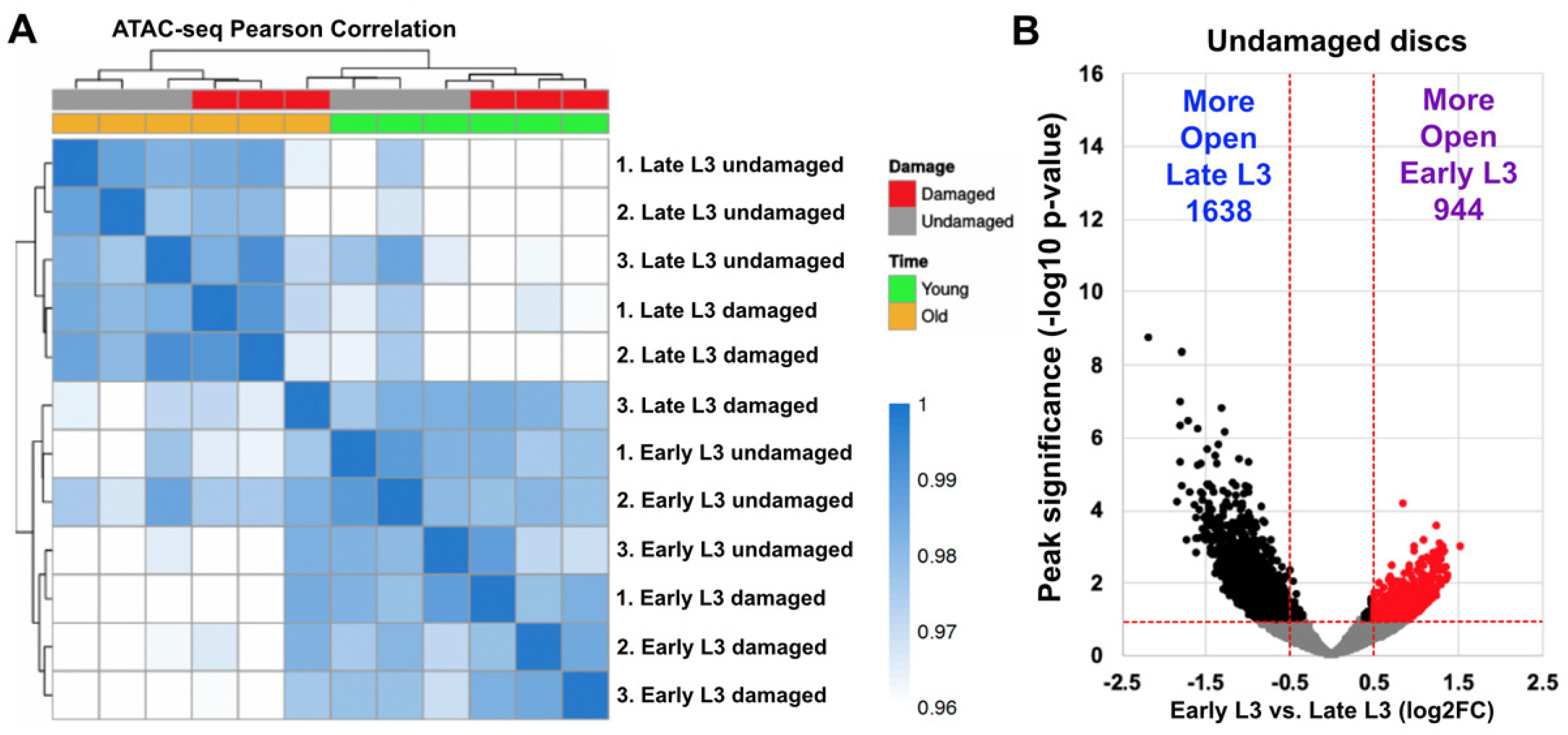
ATAC-seq changes mostly cluster by damage and age, and show chromatin accessibility alters during development, **(A)** Pearson correlation on CPM-normalized counts from DEseq2 of ATAC-seq data shows data cluster by developmental stage, damage status and biological repeat, with the exception of one late L3 damaged sample (repeat 3), **(B)** Volcano plot identifying regions that are more or less accessible in undamaged early L3 discs versus undamaged late L3 discs. Lines indicate significance threshold (p<0.1) and fold change cutoff (log2FC>0.5). The number of peaks in each category above the threshold adjusted p-values and fold change is indicated on the graph.

**Figure 3 – Figure supplement 1:**
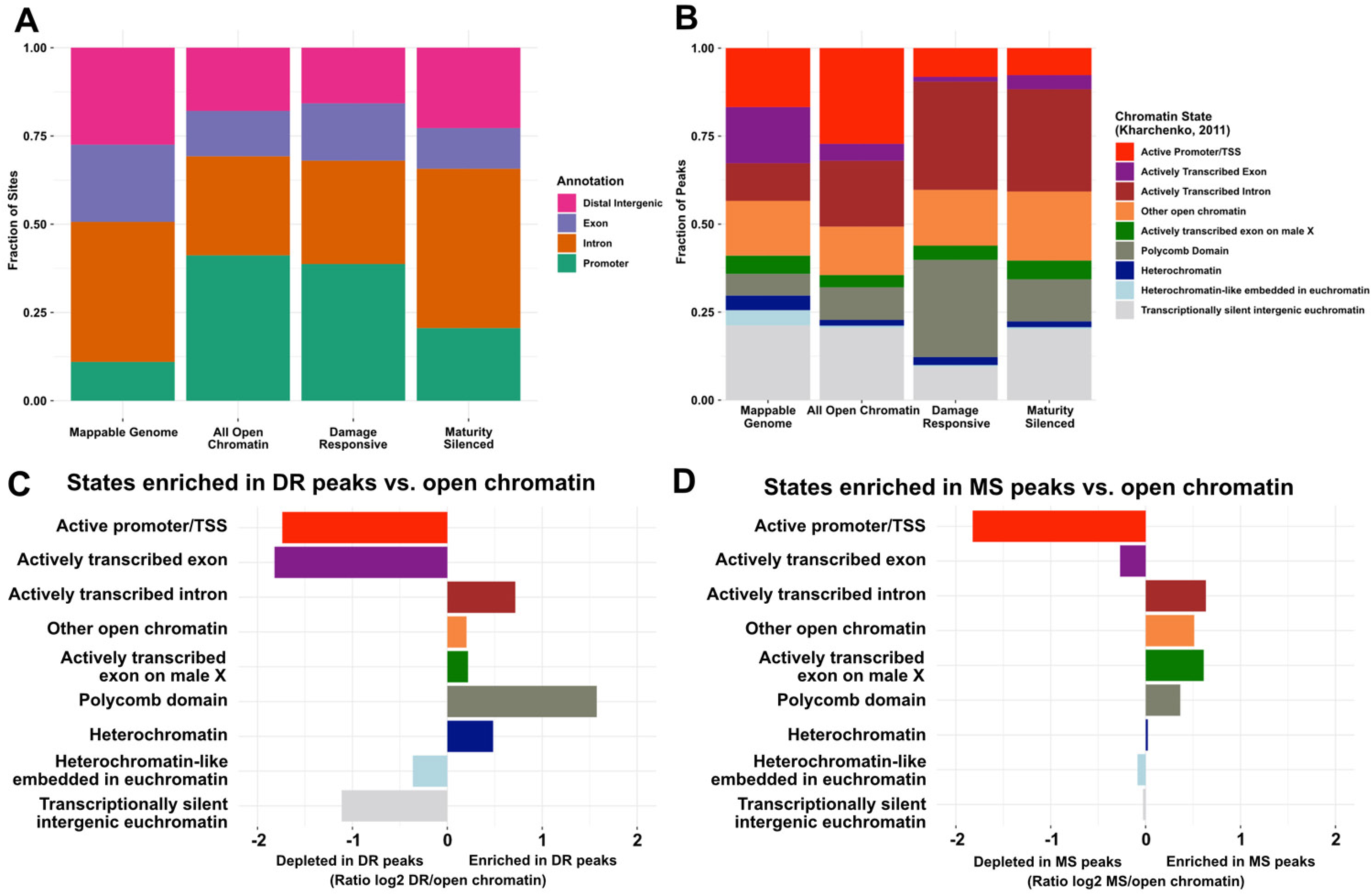
DR and MS peaks are enriched at genomic loci resembling enhancer DNA, **(A)** Stacked bar chart showing fractions of the mappable genome, total open chromatin, DR peaks and MS peaks detected by ATAC-seq that are fall within DNA categorized as distal intergenic, exon, intron and promoter DNA from annotated gene models (Flybase.org). Mappable genome is the total genome with assigned sequence, open chromatin is the total peaks detected by peak calling in all samples from the ATAC-seq analysis. MS peaks are enriched in intronic DNA, versus all open chromatin, but underrepresented at promoters, **(B)** Stacked bar chart showing fractions of the mappable genome, total open chromatin and DR and MS peaks detected by ATAC-seq that fall within DNA categorized into the different chromatin states indicated. Chromatin states were obtained from ChIP-seq data of epigenetic regulators (Kharchenko et al. 2011). Both DR and MS peaks are enriched at DNA representing actively transcribed introns and other open chromatin versus total open chromatin, while DR peaks are also enriched at DNA categorized as Polycomb regulated. Both DR and MS peaks are depleted at promoters and actively transcribed exons, (C-D) Ratiometric graphs showing the distribution of DR (C) and MS (D) peaks across the genome relative to total open chromatin detected by ATAC-seq, categorized into the chromatin states defined by ChIP-seq data of chromatin modifications from Kharchenko et al., 2011.

**Figure 3 – Figure supplement 2:**
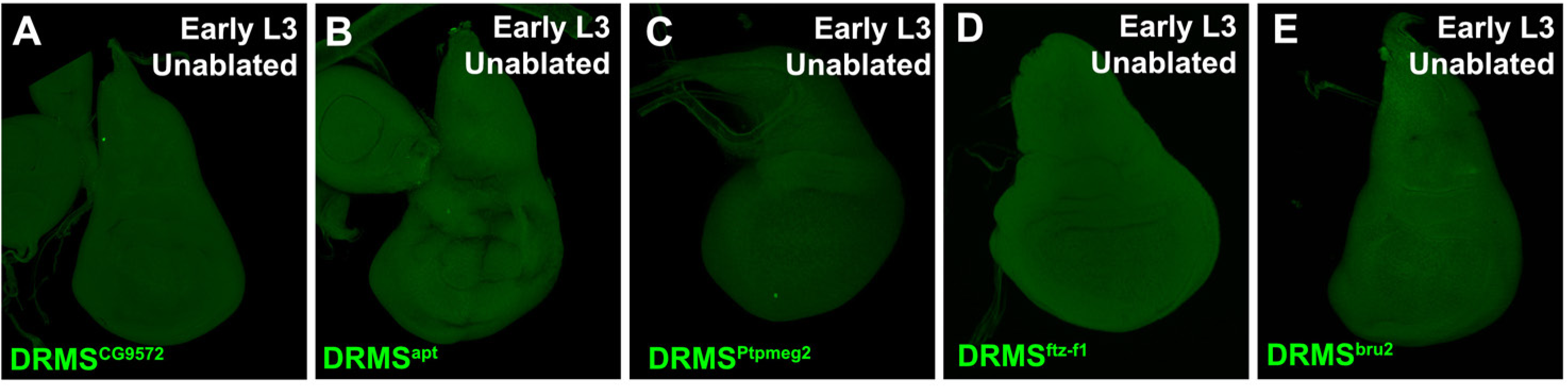
Validated DRMS enhancers are not expressed in the absence of damage, **(A-E)** Early L3 discs bearing GFP reporters of the DRMS enhancer fragments indicated. In all cases expression of GFP (green) was not observed in the absence of damage.

**Figure 4 – Figure supplement 1:**
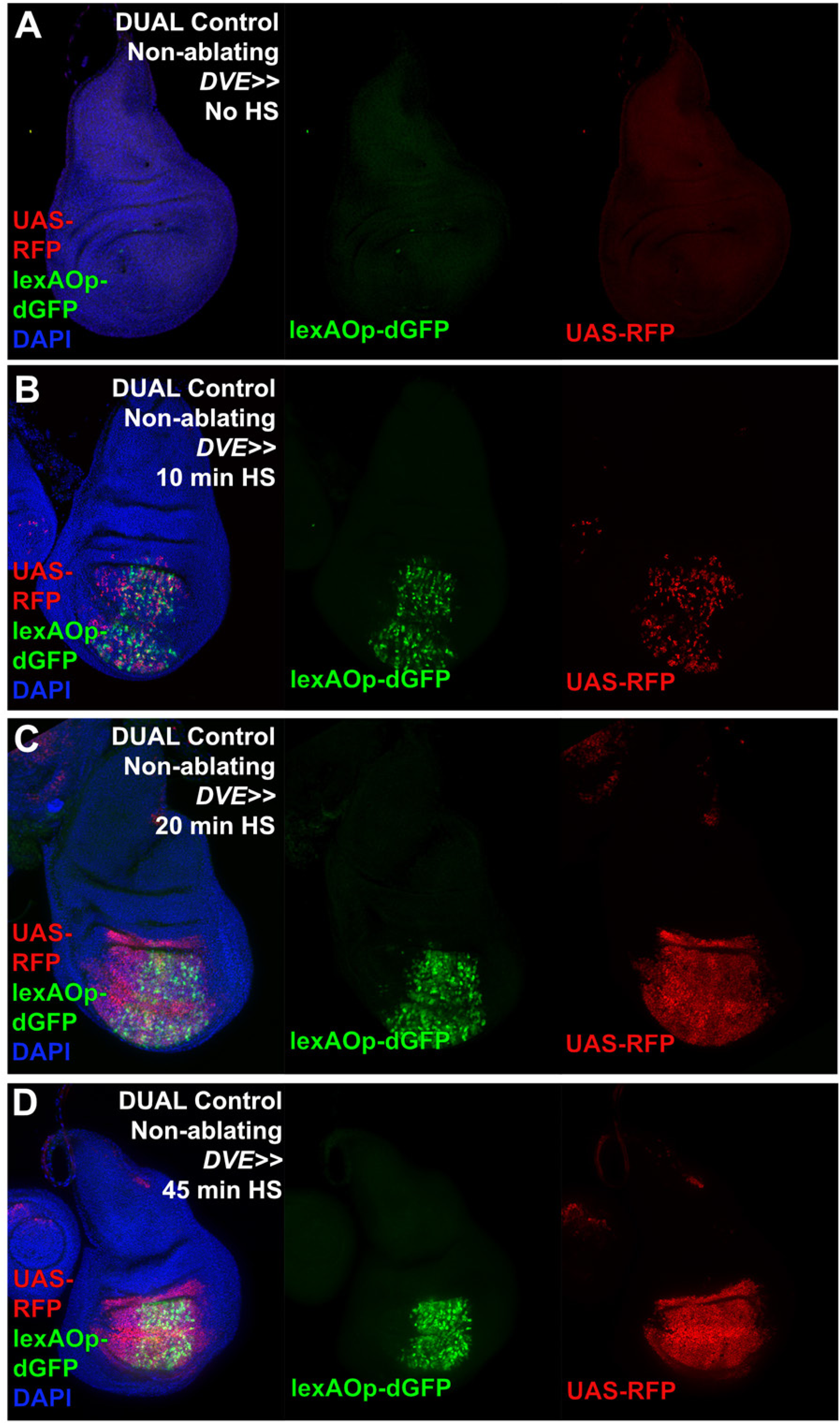
Activation of DUAL Control with different durations of heat shock, **(A-D)** Discs bearing a non-ablating DUAL Control *DVE>>GAL4* crossed to *lexAOp-dGFP; UAS-RFP* fluorescent reporters and imaged at 24 hr. In the absence of a heat shock (HS) discs show no GFP (green) or RFP (red) expression. After a 10-minute HS both reporters are weakly activated in the pouch (B). Longer duration HS of 20 minutes (C) and 45 minutes (D) increase activity of both LexA and GAL4 sides of the DUAL Control system. DAPI: blue.

**Figure 5 – Figure supplement 1:**
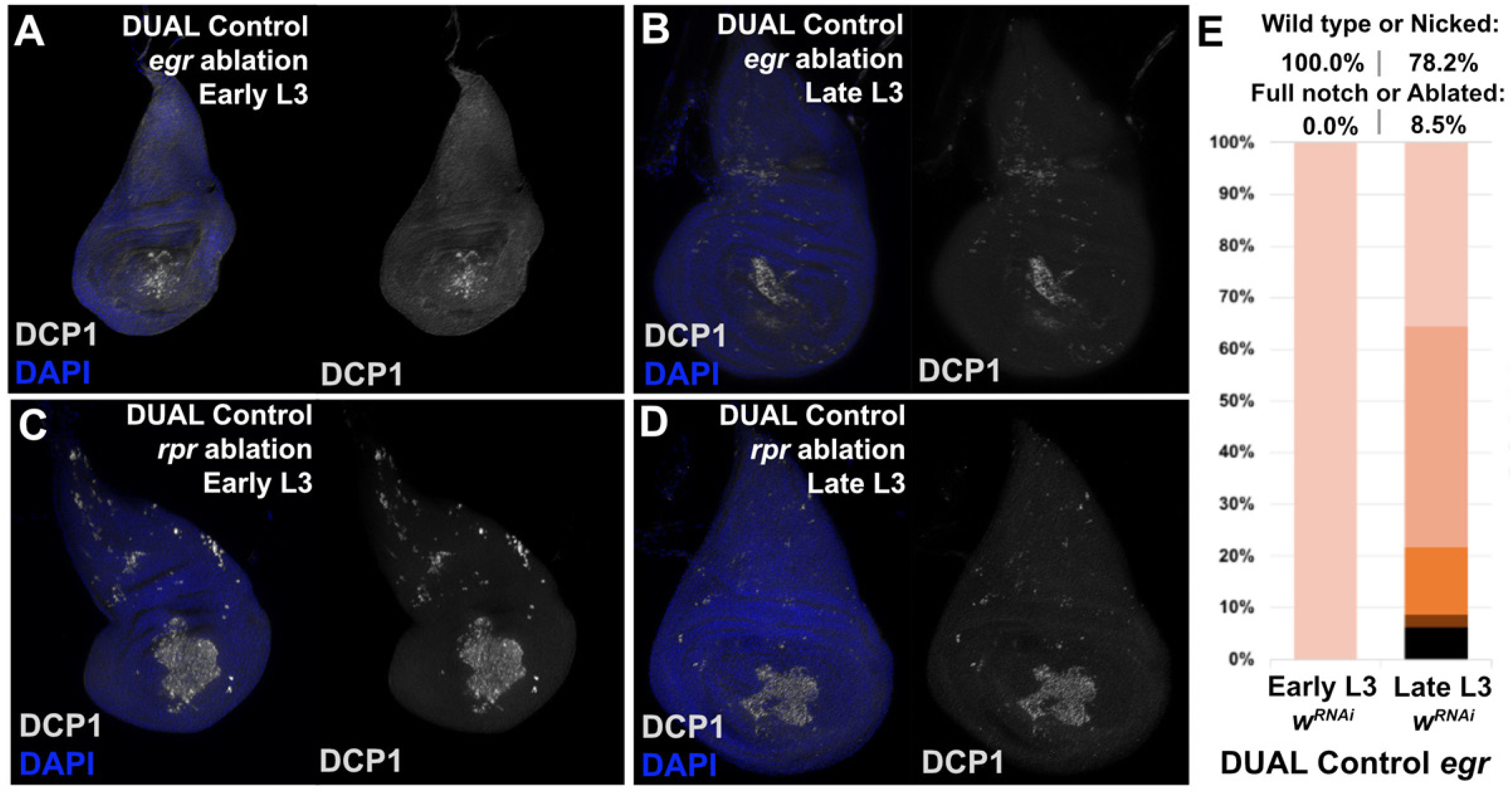
*egr*-induced ablation with DUAL Control is weaker than that of *rpr*, **(A-B)** Discs ablated by DUAL Control with *egr* in early L3 (A) or late L3 (B) stained for DCP1 (gray). Ablation is comparable between timepoints. DAPI: blue, **(C-D)** Discs as in (A-B) ablated with *rpr,* showing comparable levels of ablation at both timepoints, but a greater abundance of DCP1 than in discs damaged with *egr*, **(E)** Early or late L3 discs ablated with *egr* using DUAL control, assayed for regeneration by wing size. Graph illustrates the proportion of adults eclosing with the indicated wing phenotypes, showing overall more complete wings than those of discs ablated with *rpr* (main figure), consistent with reduced levels of ablation by *egr* in this system. *UAS-w^RNAi^* is used as a control for regeneration scoring.

**Figure 5 – Figure supplement 2:**
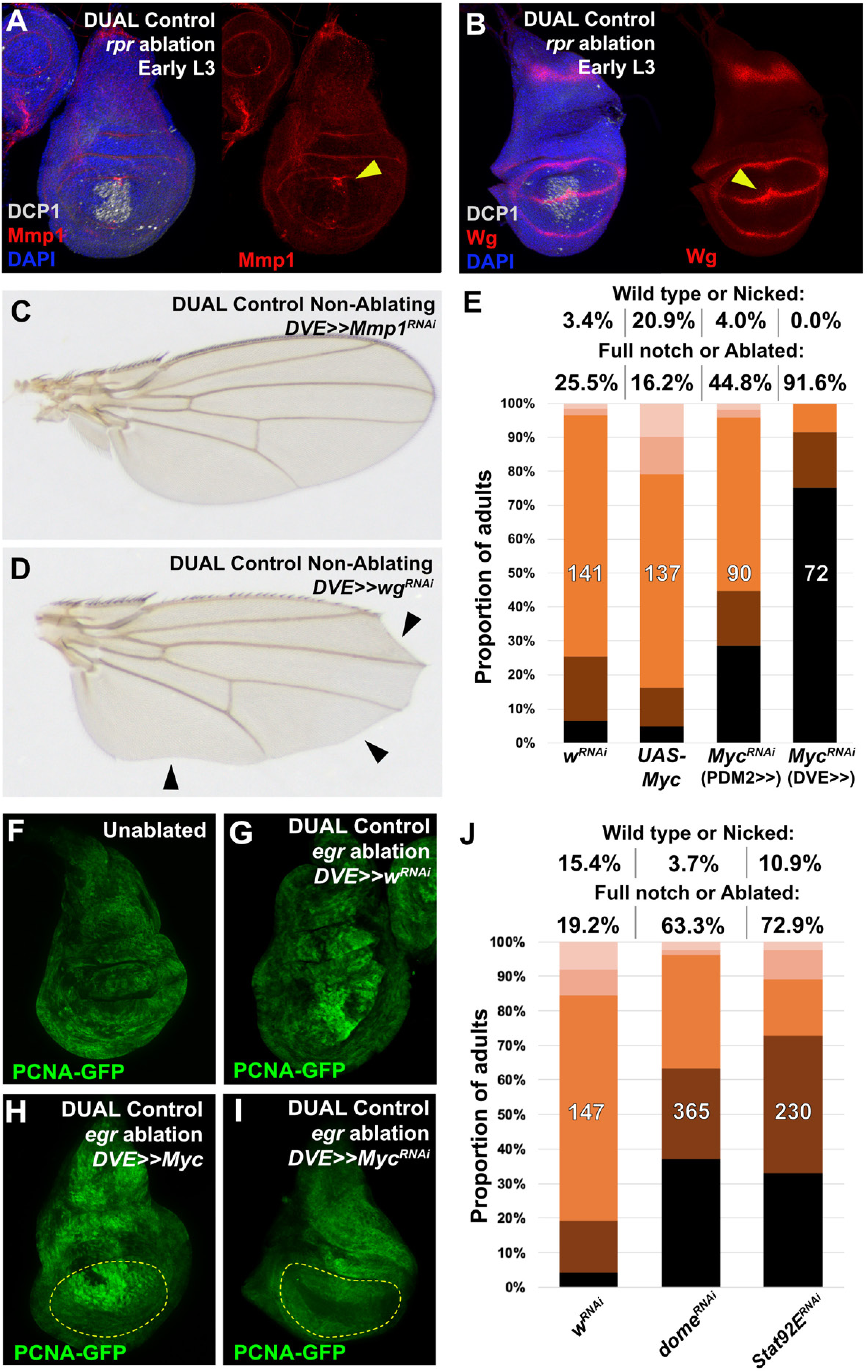
Manipulating *wg*, *Myc* and JAK/STAT signaling during regeneration using DUAL Control, **(A-B)** Early L3 discs ablated with *rpr* using DUAL Control, stained for Mmp1 (A) or Wg (B). Both are expressed at increased levels following ablation by *rpr* (arrowheads), but at lower levels than *egr* (main figure). DCP1: gray, DAPI: blue, **(C)** Adult wing resulting from knockdown of *Mmp1* using DUAL Control *DVE>>GAL4* in the absence of ablation, showing knockdown for this duration does not strongly influence wing development, **(D)** Adult wing resulting from knockdown of *wg* using DUAL Control *DVE>>GAL4* in the absence of ablation, showing loss of the margin around the entire wing typical of reduced Wg (arrowheads). This phenotype is distinguishable from ablation, which occurs only at the distal wing tip in a distinctive pattern, starting between L3 and L4 and extending inwards towards the center of the wing blade, **(E)** Late L3 discs ablated with *rpr* using DUAL control and assayed for regeneration by wing size. Graph illustrates the proportion of adults eclosing with the indicated wing phenotypes with manipulation of *Myc* expression. Ectopic activation with *UAS-Myc* leads to improved wing sizes in late L3 *rpr* ablated discs compared to *w^RNAi^*, while knock down of *Myc* using *PDM2* or *DVE* drivers prevents regrowth, with the *DVE* driver having a stronger effect, **(F-I)** Discs bearing a *PCNA-GFP* reporter of E2F activity, and thus proliferation, in unablated discs (F), discs ablated with *egr* using DUAL Control (G), ablated with *egr* using DUAL Control and ectopically expressing *Myc* in the pouch with *DVE>>GAL4* (H), and ablated with *egr* using DUAL Control and expressing *Myc* RNAi in the pouch with *DVE>>GAL4* (I). Proliferation (GFP, green) is increased in blastema cells as a result of ablation, strongly upregulated throughout the pouch as a result of ectopic *Myc*, and significantly reduced in cells with *Myc* knockdown. Yellow outline indicates the pouch, **(J)** Discs ablated with *rpr* using DUAL control and expressing the indicated RNAi transgenes using *DVE>>GAL4*, assayed for regeneration by adult wing size. Graph illustrates the proportion of adults eclosing with the indicated wing phenotypes. Loss of JAK/STAT signaling via knockdown of *dome* or *Stat92E* during regeneration strongly reduces wing regrowth compared to *w^RNAi^* control.

**Figure 6 – Figure supplement 1:**
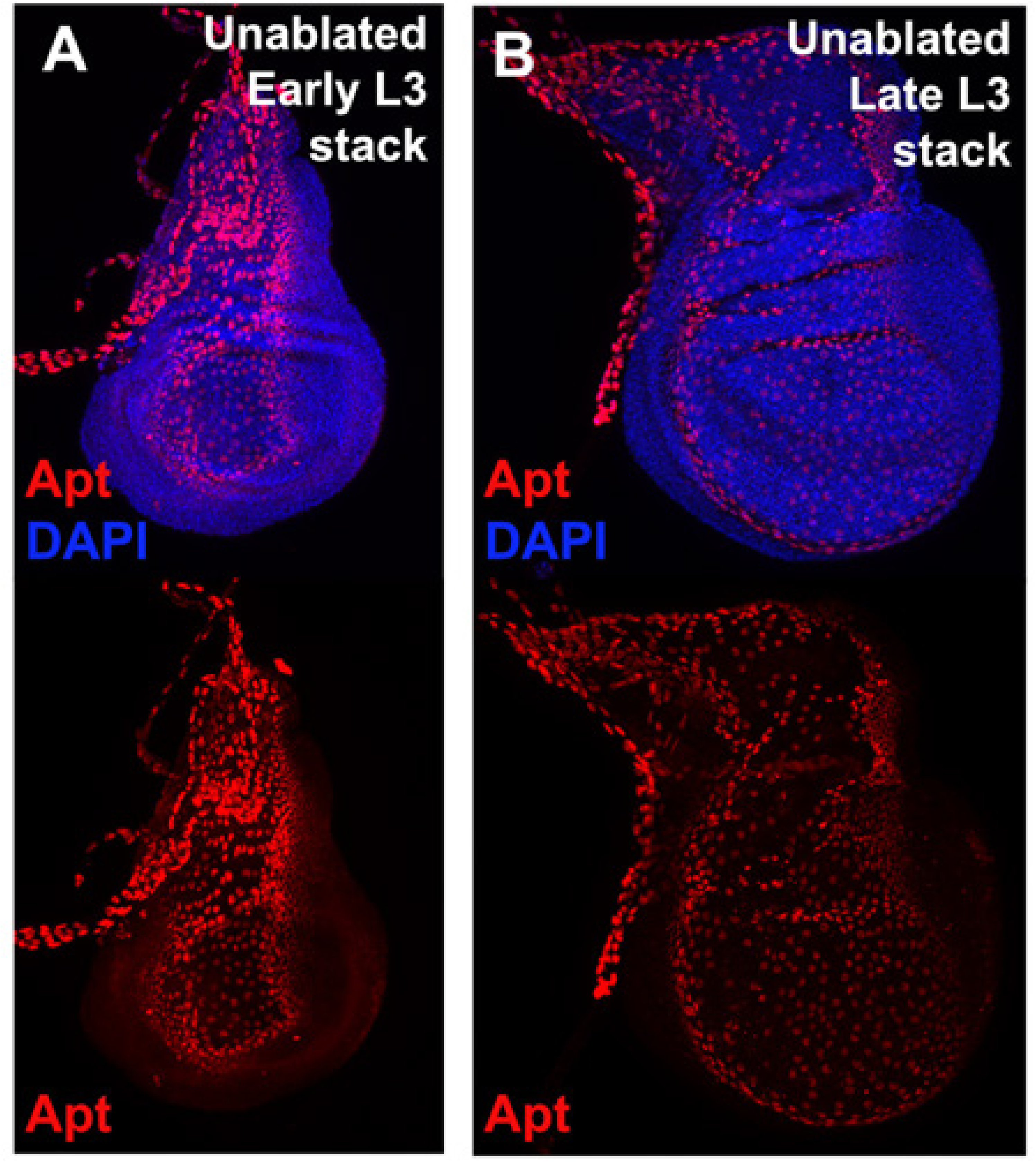
Expression of *apt* in developing discs, **(A-B)** Levels of Apt detected by an anti-Apt antibody (red) in an early L3 disc (A) and late L3 disc (B), showing very strong expression in the trachea and cells of the peripodial membrane overlying the disc proper. Apt expression is not detected in the disc proper in unablated discs. DAPI: blue.

**Figure 7 – Figure supplement 1:**
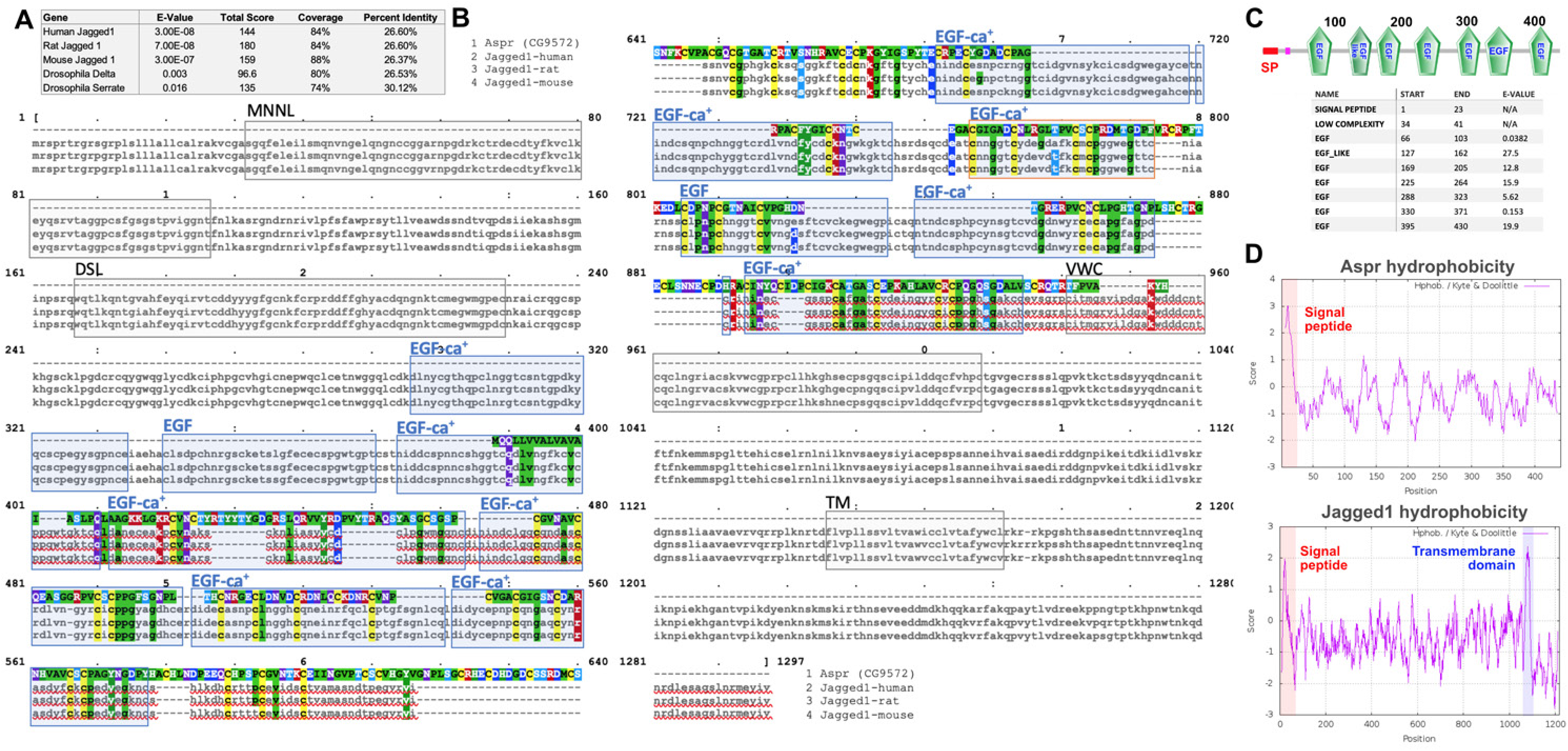
*asperous* (*aspr*, *CG9572*) peptide sequence is similar to Notch ligands but likely lacks a transmembrane domain. **(A)** Table showing protein blast (blastp) results of Aspr against known Notch ligands of *Drosophila* (Delta and Serrate) and Jagged proteins of human, rat and mouse, showing a high similarity to the mammalian Jagged proteins, **(B)** Alignment of Aspr protein sequence with that of Jagged from human, rat and mouse, showing high regions of similarity in predicted EGF repeats. Blue boxes: Predicted EGF or EGF-ca+ domains in all species, Orange box: EGF repeat in mouse and rat. MNNL: N terminus of Notch ligand domain, DSL: Delta serrate ligand domain, VWC: von Willebrand factor type C domain, TM: Transmembrane domain, (Clustal omega, https://www.ebi.ac.uk/services), **(C)** Predicted domain structure of Aspr showing EGF repeats (green) and an N terminal signal peptide (SP, red box), Coordinates of the predicted domains in Aspr, and their expected value is shown, (SMART protein annotation tool, http://smart.embl-heidelberg.de), **(D-E)** Hydropathy plots of the Aspr peptide sequence (D) and mouse Jag1 (E), showing the N-terminal hydrophobic signal peptide sequence (red box), and the transmembrane region in Jag1 (blue box) that is absent from Aspr, (ProtScale, https://web.expasy.org/protscale).

**Figure 7 – Figure supplement 2:**
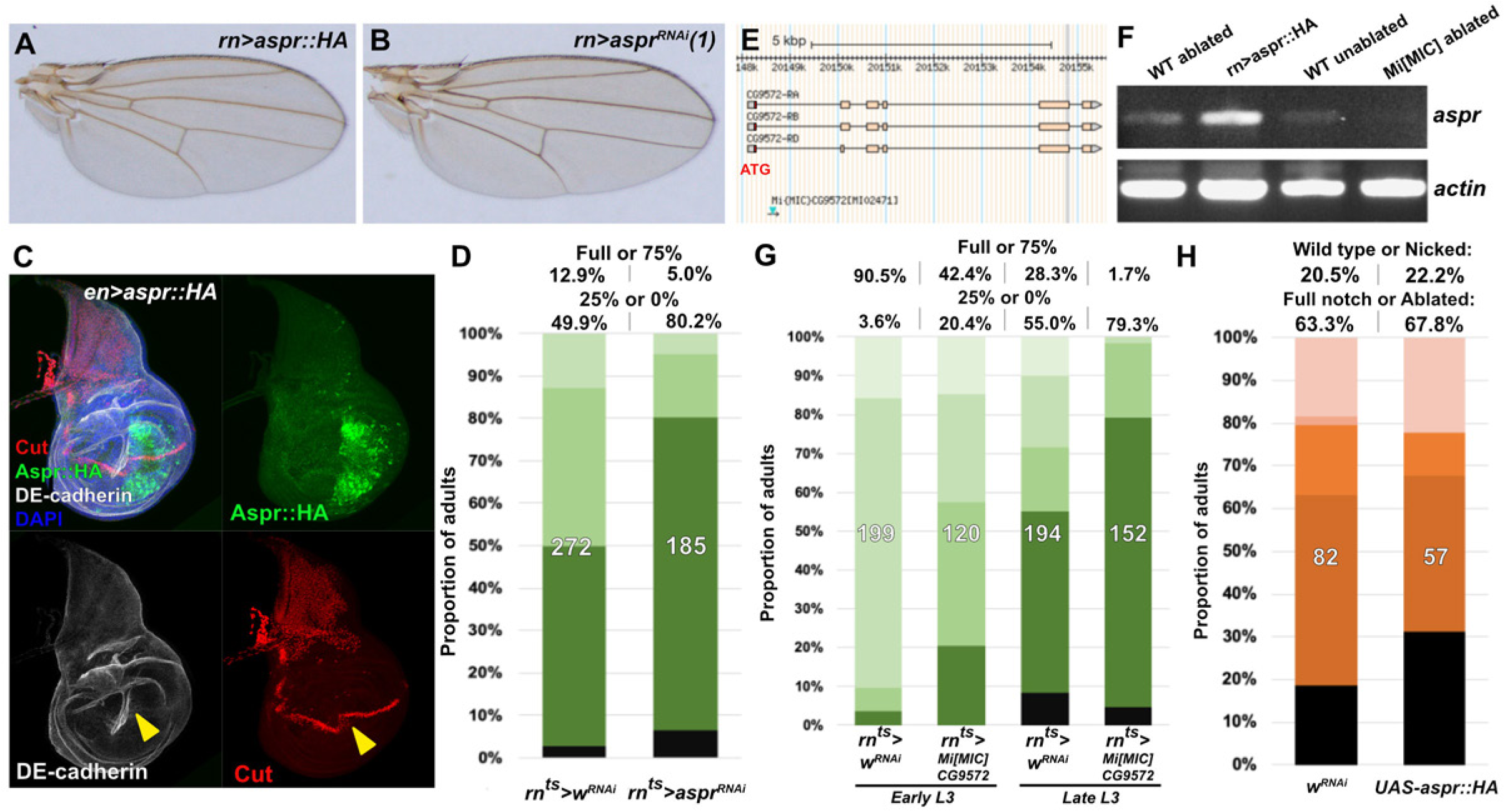
*aspr* is necessary for optimal regeneration, **(A-B)** Adult wings from discs ectopically expressing *aspr* with *rn-GAL4* driving *UAS-aspr::HA* (A), or knockdown of *aspr* using *UAS-aspr^RNAi^*. Wings have no obvious developmental defects, **(C)** Disc expressing Aspr::HA (green) in the posterior compartment using *en-GAL4*. DE-cadherin (gray) shows disruption of the pouch tissue due to unusual folding at the expression boundary of Aspr::HA (arrowhead). This fold is further highlighted by the disrupted Cut pattern (red, arrowhead), although levels and timing of Cut expression are otherwise unaffected, **(D)** Early L3 (day 7) discs ablated with *rn^ts^>egr* and expressing control *w^RNAi^* and *aspr^RNAi^*, assayed for regeneration by adult wing size as in (Smith-Bolton et al. 2009). Graph illustrates the proportion of adults eclosing with the indicated wing phenotypes, with the percentage of wings scored as “full or 75%” wing size and “25% or 0%” wing size shown above, **(E)** Genome browser screenshot (Flybase.org) showing position and orientation of the *Mi[MIC]CG9572* transposon insertion within the first intron (after the coding start site, ATG, red) of *aspr*. The GFP on the transposon shows enhancer trap activity of *aspr*, while the intronic position likely prematurely terminates *aspr* transcription, **(F)** Semiquantitative PCR to detect *aspr* and *act5C* (*actin*) mRNA from ablated discs (*rn^ts^>egr*), discs overexpressing *aspr* (rn>aspr::HA), unablated discs (*rn^ts^>egr* no upshift) and *aspr* hemizygous ablated discs (Mi[MIC]/Y;;*rn^ts^>egr*). *aspr* expression is increased upon damage in wild type discs, but not the Mi[MIC] line, suggesting it is a transcriptional mutant. **(G)** Wing regeneration assay as in (D) of early and late L3 (day 7 and day 9) discs ablated with *rn^ts^>egr,* comparing regeneration in male control flies versus hemizygous males with the *Mi[MIC]CG9572* insertion. The Mi[MIC] line has strongly reduced ability to regrow wing tissue compared to the control at two different developmental time points, **(H)** Late L3 discs ablated with *rpr* using DUAL control and expressing *UAS-aspr::HA* using *DVE>>GAL4*, assayed for regeneration by adult wing size and compared to the standard control *w^RNAi^*. Graph illustrates the proportion of adults eclosing with the indicated wing phenotypes. Overexpression of Aspr::HA does not strongly affect regeneration.

