## Supplemental methods file 1 for "Regenerative capacity in *Drosophila* imaginal discs is controlled by damage-responsive, maturity-silenced enhancers"

```
MEME version 5

ALPHABET= ACGT

strands: + -

Background letter frequencies
A 0.25 C 0.25 G 0.25 T 0.25

MOTIF Dmelanogaster-FlyFactorSurvey-BH2_Cell_FBgn0004854
letter-probability matrix: alength= 4 w= 9 nsites= 100 E= 0
 0.0952381  0.3333333  0.3333333  0.2380952
 0.2857143  0.0952381  0.04761905  0.5714286
 0.000000  0.000000  0.000000  1.000000
 1.000000  0.000000  0.000000  0.000000
 1.000000  0.000000  0.000000  0.000000
 0.2380952  0.000000  0.000000  0.7619048
 0.1428571  0.04761905  0.0952381  0.7142857
 0.000000  0.000000  1.000000  0.000000
 0.2380952  0.1904762  0.5238095  0.04761905

MOTIF Dmelanogaster-FlyFactorSurvey-BH2_SOLEXA_FBgn0004854
letter-probability matrix: alength= 4 w= 8 nsites= 100 E= 0
 0.1832061  0.2311887  0.3151581  0.2704471
 0.1984733  0.2279171  0.0610687  0.5125409
 0.03380589  0.000000  0.000000  0.9661941
 1.000000  0.000000  0.000000  0.000000
 1.000000  0.000000  0.000000  0.000000
 0.2377317  0.000000  0.03707743  0.7251908
 0.08178844  0.2846238  0.1145038  0.519084
 0.2868048  0.1014177  0.5899673  0.02181025

MOTIF Dmelanogaster-FlyFactorSurvey-br_SANGER_10_FBgn0000210
letter-probability matrix: alength= 4 w= 7 nsites= 100 E= 0
 0.08333333  0.000000  0.9166667  0.000000
 1.000000  0.000000  0.000000  0.000000
 0.000000  0.000000  0.000000  1.000000
 0.000000  0.000000  0.000000  1.000000
 1.000000  0.000000  0.000000  0.000000
 0.000000  0.08333333  0.9166667  0.000000
 1.000000  0.000000  0.000000  0.000000

MOTIF Dmelanogaster-FlyFactorSurvey-br_SOLEXA_10_FBgn0000210
letter-probability matrix: alength= 4 w= 11 nsites= 100 E= 0
 0.2633136  0.2573964  0.3491124  0.1301775
 0.2443182  0.2215909  0.1164773  0.4176136
 0.02486188  0.002762431  0.9502762  0.02209945
 0.839779  0.05801105  0.000000  0.1022099
 0.01104972  0.01104972  0.002762431  0.9751381
 0.008287293  0.000000  0.000000  0.9917127
 1.000000  0.000000  0.000000  0.000000
 0.000000  0.000000  1.000000  0.000000
 0.8895028  0.002762431  0.1022099  0.005524862
 0.5888889  0.150000  0.05833333  0.2027778
 0.4333333  0.08888889  0.3472222  0.1305556

MOTIF Dmelanogaster-FlyFactorSurvey-br.PA_SANGER_5_FBgn0000210
letter-probability matrix: alength= 4 w= 7 nsites= 100 E= 0
 0.7272727  0.000000  0.2727273  0.000000
 0.9090909  0.09090909  0.000000  0.000000
 0.000000  0.000000  0.000000  1.000000
 1.000000  0.000000  0.000000  0.000000
 0.000000  0.000000  0.9090909  0.09090909
 0.000000  0.000000  0.000000  1.000000
 0.000000  0.000000  0.3636364  0.6363636

MOTIF Dmelanogaster-FlyFactorSurvey-br.PE_SANGER_5_FBgn0000210
letter-probability matrix: alength= 4 w= 12 nsites= 100 E= 0
 0.500000  0.000000  0.500000  0.000000
 0.200000  0.100000  0.000000  0.700000
 0.200000  0.300000  0.000000  0.500000
 0.000000  0.700000  0.100000  0.200000
 0.300000  0.200000  0.300000  0.200000
 0.300000  0.100000  0.000000  0.600000
 0.000000  1.000000  0.000000  0.000000
 0.100000  0.000000  0.100000  0.800000
 1.000000  0.000000  0.000000  0.000000
 1.000000  0.000000  0.000000  0.000000
 0.100000  0.500000  0.000000  0.400000
 0.200000  0.000000  0.800000  0.000000

MOTIF Dmelanogaster-FlyFactorSurvey-br.PL_SANGER_5_FBgn0000210
letter-probability matrix: alength= 4 w= 9 nsites= 100 E= 0
 0.3636364  0.6363636  0.000000  0.000000
 0.5454545  0.4545455  0.000000  0.000000
 0.1818182  0.000000  0.1818182  0.6363636
 1.000000  0.000000  0.000000  0.000000
 0.000000  0.000000  0.9090909  0.09090909
 1.000000  0.000000  0.000000  0.000000
 0.000000  0.5454545  0.2727273  0.1818182
 0.000000  0.7272727  0.1818182  0.09090909
 0.7272727  0.09090909  0.000000  0.1818182

MOTIF Dmelanogaster-FlyFactorSurvey-br.Z1_FlyReg_FBgn0000210
letter-probability matrix: alength= 4 w= 8 nsites= 100 E= 0
 0.7058824  0.000000  0.1176471  0.1764706
 0.7058824  0.1176471  0.1764706  0.000000
 0.5294118  0.000000  0.000000  0.4705882
 0.000000  0.1764706  0.000000  0.8235294
 0.8235294  0.000000  0.1176471  0.05882353
 0.05882353  0.2352941  0.5294118  0.1764706
 0.6470588  0.1764706  0.1176471  0.05882353
 0.9411765  0.05882353  0.000000  0.000000

MOTIF Dmelanogaster-FlyFactorSurvey-br.Z2_FlyReg_FBgn0000210
letter-probability matrix: alength= 4 w= 8 nsites= 100 E= 0
 0.3333333  0.1428571  0.04761905  0.4761905
 0.000000  0.7142857  0.1904762  0.0952381
 0.000000  0.000000  0.000000  1.000000
 1.000000  0.000000  0.000000  0.000000
 0.04761905  0.000000  0.2380952  0.7142857
 0.0952381  0.2380952  0.2380952  0.4285714
 0.5714286  0.04761905  0.04761905  0.3333333
 0.5238095  0.1904762  0.04761905  0.2380952

MOTIF Dmelanogaster-FlyFactorSurvey-br.Z3_FlyReg_FBgn0000210
letter-probability matrix: alength= 4 w= 8 nsites= 100 E= 0
 0.437500  0.187500  0.187500  0.187500
 0.750000  0.062500  0.000000  0.187500
 0.625000  0.000000  0.375000  0.000000
 0.062500  0.687500  0.000000  0.250000
 0.000000  0.000000  0.062500  0.937500
 0.437500  0.000000  0.187500  0.375000
 0.000000  0.000000  0.625000  0.375000
 0.000000  0.000000  0.062500  0.937500

MOTIF Dmelanogaster-FlyFactorSurvey-br.Z4_FlyReg_FBgn0000210
letter-probability matrix: alength= 4 w= 6 nsites= 100 E= 0
 0.625000  0.000000  0.375000  0.000000
 0.250000  0.625000  0.000000  0.125000
 0.125000  0.000000  0.000000  0.875000
 1.000000  0.000000  0.000000  0.000000
 1.000000  0.000000  0.000000  0.000000
 0.125000  0.000000  0.000000  0.875000

MOTIF Dmelanogaster-FlyFactorSurvey-btd_NAR_FBgn0000233
letter-probability matrix: alength= 4 w= 10 nsites= 100 E= 0
 0.4827586  0.1034483  0.2413793  0.1724138
 0.3448276  0.000000  0.6551724  0.000000
 0.137931  0.000000  0.6206897  0.2413793
 0.06896552  0.000000  0.9310345  0.000000
 0.000000  0.000000  1.000000  0.000000
 0.000000  0.000000  1.000000  0.000000
 0.06896552  0.9310345  0.000000  0.000000
 0.000000  0.000000  1.000000  0.000000
 0.000000  0.000000  0.6896552  0.3103448
 0.5172414  0.03448276  0.3103448  0.137931

MOTIF Dmelanogaster-FlyFactorSurvey-dm_Max_SANGER_10_FBgn0000472
letter-probability matrix: alength= 4 w= 10 nsites= 100 E= 0
 0.375000  0.04166667  0.500000  0.08333333
 0.375000  0.3333333  0.1666667  0.125000
 0.000000  1.000000  0.000000  0.000000
 1.000000  0.000000  0.000000  0.000000
 0.000000  1.000000  0.000000  0.000000
 0.04166667  0.000000  0.9583333  0.000000
 0.000000  0.000000  0.000000  1.000000
 0.000000  0.000000  1.000000  0.000000
 0.04166667  0.125000  0.8333333  0.000000
 0.2083333  0.04166667  0.2083333  0.5416667

MOTIF Dmelanogaster-FlyFactorSurvey-Ets21c_SANGER_5_FBgn0005660
letter-probability matrix: alength= 4 w= 9 nsites= 100 E= 0
 0.6315789  0.05263158  0.3157895  0.000000
 0.000000  0.4736842  0.000000  0.5263158
 0.1578947  0.000000  0.000000  0.8421053
 0.000000  0.000000  0.000000  1.000000
 0.000000  1.000000  0.000000  0.000000
 0.000000  1.000000  0.000000  0.000000
 0.000000  0.000000  1.000000  0.000000
 0.000000  0.000000  0.8947368  0.1052632
 0.2777778  0.1666667  0.000000  0.5555556

MOTIF Dmelanogaster-FlyFactorSurvey-fru_SANGER_10_FBgn0004652
letter-probability matrix: alength= 4 w= 14 nsites= 100 E= 0
 0.255814  0.4651163  0.1860465  0.09302326
 0.250000  0.4318182  0.1818182  0.1363636
 0.2826087  0.4347826  0.1304348  0.1521739
 0.1521739  0.500000  0.1086957  0.2391304
 0.173913  0.3695652  0.2826087  0.173913
 0.5652174  0.1304348  0.2608696  0.04347826
 0.500000  0.4565217  0.02173913  0.02173913
 0.8695652  0.000000  0.1304348  0.000000
 0.000000  0.02173913  0.8695652  0.1086957
 0.000000  0.000000  0.000000  1.000000
 0.9782609  0.000000  0.02173913  0.000000
 1.000000  0.000000  0.000000  0.000000
 0.08695652  0.3695652  0.000000  0.5434783
 0.7555556  0.02222222  0.1333333  0.08888889

MOTIF Dmelanogaster-FlyFactorSurvey-fru_SOLEXA_5_FBgn0004652
letter-probability matrix: alength= 4 w= 24 nsites= 100 E= 0
 0.2461059  0.3831776  0.1993769  0.1713396
 0.260274  0.3589041  0.2438356  0.1369863
 0.2438356  0.3452055  0.2219178  0.1890411
 0.2575342  0.3506849  0.2273973  0.1643836
 0.3150685  0.339726  0.1972603  0.1479452
 0.309589  0.2958904  0.2410959  0.1534247
 0.260274  0.3643836  0.2246575  0.1506849
 0.2219178  0.3506849  0.2082192  0.2191781
 0.260274  0.3589041  0.1945205  0.1863014
 0.1616438  0.4520548  0.2191781  0.1671233
 0.430137  0.1945205  0.2931507  0.08219178
 0.3945205  0.5561644  0.000000  0.04931507
 1.000000  0.000000  0.000000  0.000000
 0.005479452  0.000000  0.9917808  0.002739726
 0.000000  0.002739726  0.002739726  0.9945205
 0.9342466  0.008219178  0.000000  0.05753425
 0.9890411  0.008219178  0.002739726  0.000000
 0.1424658  0.5123288  0.05479452  0.290411
 0.3972603  0.169863  0.2383562  0.1945205
 0.3917808  0.2821918  0.1863014  0.139726
 0.3315068  0.3150685  0.2273973  0.1260274
 0.3780822  0.2739726  0.2438356  0.1041096
 0.3452055  0.2821918  0.200000  0.1726027
 0.280000  0.3371429  0.1914286  0.1914286

MOTIF Dmelanogaster-FlyFactorSurvey-fru.PG_SANGER_5_FBgn0004652
letter-probability matrix: alength= 4 w= 7 nsites= 100 E= 0
 0.000000  0.200000  0.800000  0.000000
 1.000000  0.000000  0.000000  0.000000
 1.000000  0.000000  0.000000  0.000000
 0.900000  0.000000  0.000000  0.100000
 0.800000  0.200000  0.000000  0.000000
 0.800000  0.000000  0.000000  0.200000
 0.500000  0.000000  0.000000  0.500000

MOTIF Dmelanogaster-FlyFactorSurvey-ftz.f1_FlyReg_FBgn0001078
letter-probability matrix: alength= 4 w= 18 nsites= 100 E= 0
 0.000000  0.000000  0.6666667  0.3333333
 0.000000  1.000000  0.000000  0.000000
 0.3333333  0.000000  0.6666667  0.000000
 0.000000  0.000000  1.000000  0.000000
 0.000000  0.3333333  0.000000  0.6666667
 0.000000  0.000000  1.000000  0.000000
 0.6666667  0.000000  0.000000  0.3333333
 0.000000  1.000000  0.000000  0.000000
 0.000000  1.000000  0.000000  0.000000
 0.000000  0.000000  0.000000  1.000000
 0.000000  0.000000  0.000000  1.000000
 0.000000  0.6666667  0.3333333  0.000000
 0.3333333  0.000000  0.6666667  0.000000
 0.3333333  0.000000  0.6666667  0.000000
 0.6666667  0.3333333  0.000000  0.000000
 0.000000  0.6666667  0.000000  0.3333333
 0.000000  0.000000  0.3333333  0.6666667
 0.000000  0.000000  0.6666667  0.3333333

MOTIF Dmelanogaster-FlyFactorSurvey-ftz.f1_SANGER_5_FBgn0001078
letter-probability matrix: alength= 4 w= 7 nsites= 100 E= 0
 1.000000  0.000000  0.000000  0.000000
 1.000000  0.000000  0.000000  0.000000
 0.000000  0.000000  1.000000  0.000000
 0.000000  0.000000  1.000000  0.000000
 0.000000  0.000000  0.000000  1.000000
 0.000000  1.000000  0.000000  0.000000
 1.000000  0.000000  0.000000  0.000000

MOTIF Dmelanogaster-FlyFactorSurvey-kay_Jra_SANGER_5_FBgn0001291
letter-probability matrix: alength= 4 w= 11 nsites= 100 E= 0
 0.3488372  0.06976744  0.3953488  0.1860465
 0.5813953  0.2325581  0.1395349  0.04651163
 0.000000  0.000000  0.000000  1.000000
 0.000000  0.000000  0.7674419  0.2325581
 1.000000  0.000000  0.000000  0.000000
 0.09302326  0.4186047  0.4883721  0.000000
 0.000000  0.000000  0.02325581  0.9767442
 0.000000  1.000000  0.000000  0.000000
 1.000000  0.000000  0.000000  0.000000
 0.000000  0.5581395  0.02325581  0.4186047
 0.06976744  0.6046512  0.2325581  0.09302326

MOTIF Dmelanogaster-FlyFactorSurvey-kay_Jra_SANGER_5_FBgn0001297
letter-probability matrix: alength= 4 w= 11 nsites= 100 E= 0
 0.3488372  0.06976744  0.3953488  0.1860465
 0.5813953  0.2325581  0.1395349  0.04651163
 0.000000  0.000000  0.000000  1.000000
 0.000000  0.000000  0.7674419  0.2325581
 1.000000  0.000000  0.000000  0.000000
 0.09302326  0.4186047  0.4883721  0.000000
 0.000000  0.000000  0.02325581  0.9767442
 0.000000  1.000000  0.000000  0.000000
 1.000000  0.000000  0.000000  0.000000
 0.000000  0.5581395  0.02325581  0.4186047
 0.06976744  0.6046512  0.2325581  0.09302326

MOTIF Dmelanogaster-FlyFactorSurvey-ken_SANGER_10_FBgn0011236
letter-probability matrix: alength= 4 w= 9 nsites= 100 E= 0
 0.375000  0.04166667  0.5833333  0.000000
 0.08333333  0.000000  0.9166667  0.000000
 0.5416667  0.08333333  0.04166667  0.3333333
 0.000000  0.000000  1.000000  0.000000
 0.9583333  0.000000  0.000000  0.04166667
 1.000000  0.000000  0.000000  0.000000
 1.000000  0.000000  0.000000  0.000000
 0.000000  0.000000  1.000000  0.000000
 0.000000  0.1666667  0.125000  0.7083333

MOTIF Dmelanogaster-FlyFactorSurvey-ken_SOLEXA_5_FBgn0011236
letter-probability matrix: alength= 4 w= 15 nsites= 100 E= 0
 0.2543021  0.3632887  0.2351816  0.1472275
 0.2322097  0.3595506  0.2340824  0.1741573
 0.2742537  0.3712687  0.2052239  0.1492537
 0.244403  0.3376866  0.2220149  0.1958955
 0.3880597  0.03731343  0.5018657  0.07276119
 0.08395522  0.001865672  0.8712687  0.04291045
 0.4608209  0.02052239  0.03358209  0.4850746
 0.000000  0.000000  0.988806  0.01119403
 0.983209  0.000000  0.000000  0.01679104
 0.9925373  0.003731343  0.001865672  0.001865672
 0.9981343  0.000000  0.000000  0.001865672
 0.003731343  0.000000  0.9850746  0.01119403
 0.003731343  0.3432836  0.0858209  0.5671642
 0.4033771  0.1669794  0.2270169  0.2026266
 0.2686869  0.2969697  0.3050505  0.1292929

MOTIF Dmelanogaster-FlyFactorSurvey-klu_SANGER_10_FBgn0013469
letter-probability matrix: alength= 4 w= 11 nsites= 100 E= 0
 0.1111111  0.1111111  0.1111111  0.6666667
 0.1578947  0.000000  0.8421053  0.000000
 0.050000  0.500000  0.000000  0.450000
 0.000000  0.000000  1.000000  0.000000
 0.000000  0.000000  0.350000  0.650000
 0.050000  0.000000  0.950000  0.000000
 0.000000  0.000000  1.000000  0.000000
 0.000000  0.000000  1.000000  0.000000
 0.100000  0.000000  0.000000  0.900000
 0.000000  0.000000  1.000000  0.000000
 0.000000  0.100000  0.550000  0.350000

MOTIF Dmelanogaster-FlyFactorSurvey-klu_SOLEXA_5_FBgn0013469
letter-probability matrix: alength= 4 w= 15 nsites= 100 E= 0
 0.177305  0.1276596  0.3262411  0.3687943
 0.07300885  0.01106195  0.7964602  0.119469
 0.07355865  0.4572565  0.01988072  0.4493042
 0.0139165  0.001988072  0.9801193  0.003976143
 0.027833  0.01192843  0.333996  0.6262425
 0.05168986  0.000000  0.9383698  0.009940358
 0.01988072  0.009940358  0.9642147  0.005964215
 0.01789264  0.000000  0.9761431  0.005964215
 0.1411531  0.09542744  0.01192843  0.7514911
 0.005964215  0.000000  0.9761431  0.01789264
 0.05168986  0.04771372  0.5924453  0.3081511
 0.2007952  0.1272366  0.4055666  0.2664016
 0.1888668  0.1689861  0.3737575  0.2683897
 0.1602434  0.1825558  0.3407708  0.316430
 0.1991247  0.1794311  0.3326039  0.2888403

MOTIF Dmelanogaster-FlyFactorSurvey-lim_SOLEXA_2_FBgn0026411
letter-probability matrix: alength= 4 w= 9 nsites= 100 E= 0
 0.1998167  0.2612282  0.3666361  0.172319
 0.2166538  0.228219  0.08404009  0.4710871
 0.04354033  0.0007137759  0.000000  0.9557459
 0.971449  0.000000  0.02855103  0.000000
 1.000000  0.000000  0.000000  0.000000
 0.000000  0.000000  0.000000  1.000000
 0.04354033  0.0235546  0.1063526  0.8265525
 0.608137  0.041399  0.3197716  0.03069236
 0.3133858  0.2905512  0.2937008  0.1023622

MOTIF Dmelanogaster-FlyFactorSurvey-Lim1_Cell_FBgn0026411
letter-probability matrix: alength= 4 w= 7 nsites= 100 E= 0
 0.000000  0.1111111  0.000000  0.8888889
 0.000000  0.000000  0.000000  1.000000
 1.000000  0.000000  0.000000  0.000000
 1.000000  0.000000  0.000000  0.000000
 0.000000  0.000000  0.000000  1.000000
 0.000000  0.000000  0.000000  1.000000
 0.9444444  0.000000  0.05555556  0.000000

MOTIF Dmelanogaster-FlyFactorSurvey-Lim1_SOLEXA_FBgn0026411
letter-probability matrix: alength= 4 w= 7 nsites= 100 E= 0
 0.1479401  0.258427  0.09269663  0.5009363
 0.04400749  0.05149813  0.000000  0.9044944
 0.9578652  0.000000  0.02808989  0.01404494
 1.000000  0.000000  0.000000  0.000000
 0.004681648  0.007490637  0.000000  0.9878277
 0.02902622  0.01217228  0.07865169  0.8801498
 0.7312734  0.001872659  0.2490637  0.01779026

MOTIF Dmelanogaster-FlyFactorSurvey-lmd_SANGER_5_FBgn0039039
letter-probability matrix: alength= 4 w= 14 nsites= 100 E= 0
 0.0952381  0.7142857  0.04761905  0.1428571
 0.1363636  0.1363636  0.3181818  0.4090909
 0.000000  0.1818182  0.8181818  0.000000
 0.000000  0.4545455  0.04545455  0.500000
 0.2272727  0.000000  0.7272727  0.04545455
 0.000000  0.000000  1.000000  0.000000
 0.000000  0.000000  1.000000  0.000000
 0.000000  0.000000  1.000000  0.000000
 0.000000  0.000000  1.000000  0.000000
 0.09090909  0.000000  0.9090909  0.000000
 0.000000  0.1818182  0.2272727  0.5909091
 0.3181818  0.500000  0.1363636  0.04545455
 0.09090909  0.1363636  0.4090909  0.3636364
 0.04761905  0.0952381  0.3809524  0.4761905

MOTIF Dmelanogaster-FlyFactorSurvey-lmd_SOLEXA_5_FBgn0039039
letter-probability matrix: alength= 4 w= 15 nsites= 100 E= 0
 0.1766562  0.1451104  0.2271293  0.4511041
 0.1299639  0.6119134  0.05415162  0.2039711
 0.1490468  0.2807626  0.2253033  0.3448873
 0.03292894  0.2322357  0.729636  0.005199307
 0.02426343  0.4315425  0.05199307  0.492201
 0.2010399  0.000000  0.6655113  0.1334489
 0.02253033  0.000000  0.9532062  0.02426343
 0.000000  0.000000  0.9948007  0.005199307
 0.04852686  0.000000  0.8942808  0.05719237
 0.01733102  0.000000  0.9046794  0.0779896
 0.07105719  0.000000  0.8942808  0.03466205
 0.01213172  0.1213172  0.3154246  0.5511265
 0.202773  0.4194107  0.2218371  0.1559792
 0.1420959  0.1509769  0.3410302  0.365897
 0.1413043  0.1757246  0.3550725  0.3278986

MOTIF Dmelanogaster-FlyFactorSurvey-pan_FlyReg_FBgn0085432
letter-probability matrix: alength= 4 w= 8 nsites= 100 E= 0
 0.160000  0.400000  0.120000  0.320000
 0.080000  0.120000  0.000000  0.800000
 0.080000  0.000000  0.040000  0.880000
 0.080000  0.000000  0.000000  0.920000
 0.000000  0.080000  0.800000  0.120000
 0.680000  0.000000  0.200000  0.120000
 0.320000  0.000000  0.000000  0.680000
 0.280000  0.440000  0.120000  0.160000

MOTIF Dmelanogaster-FlyFactorSurvey-peb.F1.3_SANGER_2.5_FBgn0003053
letter-probability matrix: alength= 4 w= 7 nsites= 100 E= 0
 0.7272727  0.09090909  0.09090909  0.09090909
 0.000000  0.000000  1.000000  0.000000
 0.000000  1.000000  0.000000  0.000000
 1.000000  0.000000  0.000000  0.000000
 0.000000  0.000000  0.000000  1.000000
 0.000000  1.000000  0.000000  0.000000
 0.2727273  0.6363636  0.000000  0.09090909

MOTIF Dmelanogaster-FlyFactorSurvey-PhdP_Cell_FBgn0025334
letter-probability matrix: alength= 4 w= 6 nsites= 100 E= 0
 0.2941176  0.2941176  0.000000  0.4117647
 0.05882353  0.000000  0.000000  0.9411765
 0.8823529  0.000000  0.000000  0.1176471
 0.9411765  0.000000  0.05882353  0.000000
 0.05882353  0.000000  0.000000  0.9411765
 0.05882353  0.000000  0.05882353  0.8823529

MOTIF Dmelanogaster-FlyFactorSurvey-PhdP_SOLEXA_FBgn0025334
letter-probability matrix: alength= 4 w= 6 nsites= 100 E= 0
 0.1328947  0.2986842  0.08815789  0.4802632
 0.02236842  0.000000  0.000000  0.9776316
 0.9802632  0.000000  0.000000  0.01973684
 1.000000  0.000000  0.000000  0.000000
 0.000000  0.000000  0.000000  1.000000
 0.04868421  0.02631579  0.06842105  0.8565789

MOTIF Dmelanogaster-FlyFactorSurvey-pho_FlyReg_FBgn0002521
letter-probability matrix: alength= 4 w= 14 nsites= 100 E= 0
 0.400000  0.000000  0.600000  0.000000
 0.000000  0.600000  0.400000  0.000000
 0.200000  0.400000  0.200000  0.200000
 0.200000  0.000000  0.600000  0.200000
 0.000000  0.000000  0.200000  0.800000
 0.200000  0.200000  0.000000  0.600000
 0.800000  0.000000  0.200000  0.000000
 0.000000  0.000000  0.000000  1.000000
 0.000000  0.000000  1.000000  0.000000
 0.000000  0.000000  1.000000  0.000000
 0.000000  1.000000  0.000000  0.000000
 0.000000  0.200000  0.200000  0.600000
 0.200000  0.200000  0.200000  0.400000
 0.400000  0.600000  0.000000  0.000000

MOTIF Dmelanogaster-FlyFactorSurvey-pho_SANGER_10_FBgn0002521
letter-probability matrix: alength= 4 w= 12 nsites= 100 E= 0
 0.3181818  0.500000  0.04545455  0.1363636
 0.7727273  0.000000  0.2272727  0.000000
 0.6363636  0.1363636  0.04545455  0.1818182
 0.5454545  0.09090909  0.2272727  0.1363636
 1.000000  0.000000  0.000000  0.000000
 0.000000  0.000000  0.000000  1.000000
 0.000000  0.000000  1.000000  0.000000
 0.000000  0.000000  1.000000  0.000000
 0.000000  1.000000  0.000000  0.000000
 0.000000  0.1363636  0.8636364  0.000000
 0.1363636  0.09090909  0.7727273  0.000000
 0.2272727  0.6363636  0.09090909  0.04545455

MOTIF Dmelanogaster-FlyFactorSurvey-pho_SOLEXA_5_FBgn0002521
letter-probability matrix: alength= 4 w= 15 nsites= 100 E= 0
 0.3576826  0.1889169  0.2418136  0.2115869
 0.2630332  0.3767773  0.2085308  0.1516588
 0.3123543  0.4755245  0.1351981  0.07692308
 0.6969697  0.06060606  0.1864802  0.05594406
 0.5687646  0.1375291  0.01398601  0.2797203
 0.5151515  0.1212121  0.2097902  0.1538462
 0.976690  0.000000  0.000000  0.02331002
 0.000000  0.000000  0.000000  1.000000
 0.000000  0.000000  1.000000  0.000000
 0.000000  0.000000  1.000000  0.000000
 0.08624709  0.8927739  0.000000  0.02097902
 0.02797203  0.1025641  0.7925408  0.07692308
 0.1888112  0.1445221  0.5664336  0.1002331
 0.3262911  0.4882629  0.07042254  0.1150235
 0.1905941  0.3613861  0.1633663  0.2846535

MOTIF Dmelanogaster-FlyFactorSurvey-sd_FlyReg_FBgn0003345
letter-probability matrix: alength= 4 w= 12 nsites= 100 E= 0
 0.2142857  0.000000  0.5714286  0.2142857
 0.5714286  0.07142857  0.07142857  0.2857143
 0.2857143  0.5714286  0.07142857  0.07142857
 1.000000  0.000000  0.000000  0.000000
 0.07142857  0.07142857  0.000000  0.8571429
 0.2142857  0.000000  0.07142857  0.7142857
 0.000000  0.5714286  0.000000  0.4285714
 0.2142857  0.3571429  0.2142857  0.2142857
 0.2857143  0.07142857  0.1428571  0.500000
 0.3571429  0.4285714  0.2142857  0.000000
 0.3571429  0.07142857  0.500000  0.07142857
 0.2857143  0.2142857  0.2857143  0.2142857

MOTIF Dmelanogaster-FlyFactorSurvey-Sox15_SANGER_5_FBgn0005613
letter-probability matrix: alength= 4 w= 8 nsites= 100 E= 0
 0.5714286  0.04761905  0.2380952  0.1428571
 0.6190476  0.0952381  0.0952381  0.1904762
 1.000000  0.000000  0.000000  0.000000
 0.000000  1.000000  0.000000  0.000000
 1.000000  0.000000  0.000000  0.000000
 1.000000  0.000000  0.000000  0.000000
 0.1428571  0.000000  0.000000  0.8571429
 0.2857143  0.000000  0.6666667  0.04761905

MOTIF Dmelanogaster-FlyFactorSurvey-sr_SANGER_5_FBgn0003499
letter-probability matrix: alength= 4 w= 14 nsites= 100 E= 0
 0.2105263  0.05263158  0.4736842  0.2631579
 0.1052632  0.05263158  0.2631579  0.5789474
 0.3157895  0.05263158  0.5263158  0.1052632
 0.000000  0.4210526  0.2631579  0.3157895
 0.000000  0.000000  1.000000  0.000000
 0.05263158  0.000000  0.3684211  0.5789474
 0.000000  0.000000  1.000000  0.000000
 0.000000  0.05263158  0.9473684  0.000000
 0.000000  0.000000  1.000000  0.000000
 0.1052632  0.7368421  0.000000  0.1578947
 0.05263158  0.000000  0.8421053  0.1052632
 0.000000  0.05263158  0.4736842  0.4736842
 0.05263158  0.2105263  0.4736842  0.2631579
 0.2777778  0.000000  0.4444444  0.2777778

MOTIF Dmelanogaster-FlyFactorSurvey-sr_SOLEXA_5_FBgn0003499
letter-probability matrix: alength= 4 w= 14 nsites= 100 E= 0
 0.1993599  0.1554641  0.3891175  0.2560585
 0.1335927  0.2088197  0.2542153  0.4033722
 0.1548332  0.09837468  0.5902481  0.1565441
 0.09324209  0.4610778  0.1368691  0.3088109
 0.000000  0.000000  0.9965783  0.003421728
 0.02523524  0.002566296  0.298118  0.6740804
 0.150556  0.000000  0.832763  0.01668092
 0.000000  0.000000  1.000000  0.000000
 0.000000  0.000000  0.9837468  0.01625321
 0.1227545  0.5106929  0.03892216  0.3276305
 0.05731394  0.03849444  0.8190761  0.08511548
 0.08896493  0.06971771  0.4764756  0.3648417
 0.2360992  0.1501283  0.3738238  0.2399487
 0.1922031  0.2089755  0.3245694  0.274252

MOTIF Dmelanogaster-FlyFactorSurvey-ss_tgo_SANGER_10_FBgn0003513
letter-probability matrix: alength= 4 w= 11 nsites= 100 E= 0
 0.2307692  0.000000  0.7692308  0.000000
 0.000000  0.000000  0.000000  1.000000
 0.000000  1.000000  0.000000  0.000000
 1.000000  0.000000  0.000000  0.000000
 0.000000  1.000000  0.000000  0.000000
 0.000000  0.000000  1.000000  0.000000
 0.000000  1.000000  0.000000  0.000000
 0.7692308  0.000000  0.2307692  0.000000
 1.000000  0.000000  0.000000  0.000000
 0.1538462  0.1538462  0.07692308  0.6153846
 0.000000  0.3846154  0.6153846  0.000000

MOTIF Dmelanogaster-FlyFactorSurvey-tgo_ss_SANGER_5_FBgn0003513
letter-probability matrix: alength= 4 w= 8 nsites= 100 E= 0
 0.000000  0.100000  0.000000  0.900000
 0.000000  0.000000  1.000000  0.000000
 0.000000  0.900000  0.000000  0.100000
 0.000000  0.100000  0.900000  0.000000
 0.000000  0.000000  0.000000  1.000000
 0.000000  0.000000  1.000000  0.000000
 1.000000  0.000000  0.000000  0.000000
 0.000000  0.800000  0.000000  0.200000

MOTIF Dmelanogaster-FlyFactorSurvey-Trl_FlyReg_FBgn0013263
letter-probability matrix: alength= 4 w= 10 nsites= 100 E= 0
 0.05633803  0.3098592  0.1267606  0.5070423
 0.08450704  0.3239437  0.1549296  0.4366197
 0.1690141  0.08450704  0.4225352  0.3239437
 0.000000  0.9014085  0.04225352  0.05633803
 0.04225352  0.000000  0.1971831  0.7605634
 0.01408451  0.9859155  0.000000  0.000000
 0.1830986  0.000000  0.02816901  0.7887324
 0.07042254  0.7605634  0.09859155  0.07042254
 0.09859155  0.1830986  0.1549296  0.5633803
 0.1267606  0.4647887  0.1408451  0.2676056

MOTIF Dmelanogaster-FlyFactorSurvey-Tup_Cell_FBgn0003896
letter-probability matrix: alength= 4 w= 8 nsites= 100 E= 0
 0.062500  0.562500  0.312500  0.062500
 0.312500  0.062500  0.062500  0.562500
 0.062500  0.062500  0.000000  0.875000
 0.937500  0.000000  0.000000  0.062500
 1.000000  0.000000  0.000000  0.000000
 0.062500  0.000000  0.125000  0.812500
 0.062500  0.000000  0.375000  0.562500
 0.125000  0.000000  0.875000  0.000000

MOTIF Dmelanogaster-FlyFactorSurvey-tup_SOLEXA_10_FBgn0003896
letter-probability matrix: alength= 4 w= 9 nsites= 100 E= 0
 0.2174452  0.2662257  0.2620918  0.2542373
 0.1154781  0.3573744  0.1146677  0.4124797
 0.05226904  0.000000  0.000000  0.947731
 0.9894652  0.000000  0.01053485  0.000000
 1.000000  0.000000  0.000000  0.000000
 0.000000  0.000000  0.1511345  0.8488655
 0.06320908  0.01215559  0.4371961  0.4874392
 0.301190  0.04718917  0.5954042  0.05621666
 0.3457607  0.2693927  0.2633794  0.1214672

MOTIF Dmelanogaster-FlyFactorSurvey-Tup_SOLEXA_FBgn0003896
letter-probability matrix: alength= 4 w= 7 nsites= 100 E= 0
 0.08136483  0.3700787  0.04724409  0.5013123
 0.02887139  0.000000  0.000000  0.9711286
 1.000000  0.000000  0.000000  0.000000
 1.000000  0.000000  0.000000  0.000000
 0.000000  0.000000  0.09448819  0.9055118
 0.03412073  0.000000  0.4304462  0.5354331
 0.335958  0.0183727  0.5958005  0.04986877

MOTIF Dmelanogaster-FlyFactorSurvey-twi_da_SANGER_5_FBgn0003900
letter-probability matrix: alength= 4 w= 10 nsites= 100 E= 0
 0.5217391  0.173913  0.3043478  0.000000
 0.04347826  0.826087  0.08695652  0.04347826
 0.000000  0.9565217  0.000000  0.04347826
 1.000000  0.000000  0.000000  0.000000
 0.000000  0.1304348  0.6956522  0.173913
 0.9565217  0.000000  0.04347826  0.000000
 0.000000  0.000000  0.000000  1.000000
 0.000000  0.04347826  0.9565217  0.000000
 0.000000  0.08695652  0.173913  0.7391304
 0.000000  0.3181818  0.3181818  0.3636364

MOTIF Dmelanogaster-FlyFactorSurvey-twi_FlyReg_FBgn0003900
letter-probability matrix: alength= 4 w= 12 nsites= 100 E= 0
 0.06666667  0.2666667  0.200000  0.4666667
 0.200000  0.4666667  0.200000  0.1333333
 0.06666667  0.200000  0.6666667  0.06666667
 0.000000  0.9333333  0.06666667  0.000000
 0.7333333  0.200000  0.06666667  0.000000
 0.000000  0.2666667  0.06666667  0.6666667
 0.4666667  0.06666667  0.3333333  0.1333333
 0.000000  0.000000  0.000000  1.000000
 0.000000  0.000000  0.800000  0.200000
 0.06666667  0.000000  0.1333333  0.800000
 0.06666667  0.1333333  0.1333333  0.6666667
 0.1333333  0.200000  0.5333333  0.1333333

MOTIF Dmelanogaster-FlyFactorSurvey-Ubx_Cell_FBgn0003944
letter-probability matrix: alength= 4 w= 8 nsites= 100 E= 0
 0.150000  0.250000  0.150000  0.450000
 0.000000  0.000000  0.000000  1.000000
 0.000000  0.000000  0.000000  1.000000
 0.850000  0.000000  0.000000  0.150000
 1.000000  0.000000  0.000000  0.000000
 0.000000  0.000000  0.000000  1.000000
 0.000000  0.000000  0.300000  0.700000
 0.700000  0.000000  0.300000  0.000000

MOTIF Dmelanogaster-FlyFactorSurvey-Ubx_FlyReg_FBgn0003944
letter-probability matrix: alength= 4 w= 6 nsites= 100 E= 0
 0.2321429  0.4642857  0.08928571  0.2142857
 0.5535714  0.3392857  0.000000  0.1071429
 0.9642857  0.000000  0.000000  0.03571429
 0.000000  0.000000  0.000000  1.000000
 0.05357143  0.03571429  0.000000  0.9107143
 0.875000  0.000000  0.000000  0.125000

MOTIF Dmelanogaster-FlyFactorSurvey-Ubx_SOLEXA_FBgn0003944
letter-probability matrix: alength= 4 w= 8 nsites= 100 E= 0
 0.1992481  0.1691729  0.1842105  0.4473684
 0.1090226  0.06766917  0.03007519  0.7932331
 0.02255639  0.000000  0.007518797  0.9699248
 0.7481203  0.000000  0.000000  0.2518797
 0.9887218  0.000000  0.0112782  0.000000
 0.000000  0.08646617  0.000000  0.9135338
 0.05639098  0.01503759  0.4887218  0.4398496
 0.7142857  0.000000  0.2631579  0.02255639

MOTIF Dmelanogaster-FlyFactorSurvey-z_FlyReg_FBgn0004050
letter-probability matrix: alength= 4 w= 10 nsites= 100 E= 0
 0.2682927  0.07317073  0.1219512  0.5365854
 0.02439024  0.1707317  0.07317073  0.7317073
 0.02439024  0.000000  0.9756098  0.000000
 1.000000  0.000000  0.000000  0.000000
 0.02439024  0.000000  0.9756098  0.000000
 0.195122  0.2439024  0.000000  0.5609756
 0.04878049  0.1463415  0.7073171  0.09756098
 0.4390244  0.1707317  0.2682927  0.1219512
 0.2439024  0.1219512  0.2926829  0.3414634
 0.3414634  0.1463415  0.04878049  0.4634146

MOTIF Dmelanogaster-FlyFactorSurvey-chinmo_SOLEXA_FBgn0086758
letter-probability matrix: alength= 4 w= 11 nsites= 100 E= 0
 0.404894  0.034483  0.490545  0.070078
 0.918799  0.051168  0.008899  0.021135
 0.000000  0.000000  0.000000  1.000000
 0.006674  0.000000  0.993326  0.000000
 0.000000  0.991101  0.000000  0.008899
 0.992214  0.000000  0.007786  0.000000
 0.105673  0.708565  0.078977  0.106785
 0.052280  0.420467  0.014461  0.512792
 0.132369  0.206897  0.153504  0.507230
 0.242492  0.293660  0.226919  0.236930
 0.296997  0.199110  0.311457  0.192436

MOTIF Dmelanogaster-FlyFactorSurvey-ab_SANGER_10_FBgn0259750
letter-probability matrix: alength= 4 w= 21 nsites= 100 E= 0
 0.000000  0.300000  0.400000  0.300000
 0.500000  0.150000  0.050000  0.300000
 0.200000  0.250000  0.500000  0.050000
 0.350000  0.000000  0.650000  0.000000
 0.000000  1.000000  0.000000  0.000000
 0.000000  1.000000  0.000000  0.000000
 1.000000  0.000000  0.000000  0.000000
 0.000000  0.000000  1.000000  0.000000
 0.000000  0.000000  1.000000  0.000000
 0.550000  0.100000  0.000000  0.350000
 0.350000  0.650000  0.000000  0.000000
 0.050000  0.700000  0.050000  0.200000
 0.200000  0.450000  0.050000  0.300000
 0.450000  0.250000  0.150000  0.150000
 0.200000  0.100000  0.050000  0.650000
 0.100000  0.250000  0.200000  0.450000
 0.400000  0.250000  0.050000  0.300000
 0.400000  0.100000  0.150000  0.350000
 0.250000  0.100000  0.550000  0.100000
 0.500000  0.250000  0.150000  0.100000
 0.300000  0.250000  0.450000  0.000000

MOTIF Dmelanogaster-FlyFactorSurvey-ab_SOLEXA_5_FBgn0259750
letter-probability matrix: alength= 4 w= 26 nsites= 100 E= 0
 0.2860262  0.3449782  0.1768559  0.1921397
 0.2885033  0.4078091  0.1995662  0.1041215
 0.2429501  0.3644252  0.1908894  0.2017354
 0.1952278  0.3752711  0.2299349  0.1995662
 0.3210412  0.2603037  0.1149675  0.3036876
 0.2212581  0.4229935  0.2624729  0.09327549
 0.4121475  0.04121475  0.4490239  0.09761388
 0.002169197  0.9826464  0.000000  0.01518438
 0.000000  0.9869848  0.000000  0.01301518
 0.9739696  0.002169197  0.000000  0.02386117
 0.000000  0.000000  0.989154  0.01084599
 0.000000  0.000000  0.9978308  0.002169197
 0.648590  0.06290672  0.01735358  0.2711497
 0.2819957  0.7071584  0.004338395  0.006507592
 0.05422993  0.8112798  0.03470716  0.09978308
 0.1171367  0.4273319  0.0867679  0.3687636
 0.373102  0.2776573  0.1518438  0.197397
 0.3058568  0.2494577  0.07809111  0.3665944
 0.2255965  0.2819957  0.1518438  0.340564
 0.3123644  0.308026  0.1301518  0.2494577
 0.340564  0.2581345  0.2212581  0.1800434
 0.362256  0.2342733  0.2472885  0.1561822
 0.4663774  0.2776573  0.1626898  0.09327549
 0.4251627  0.2689805  0.1800434  0.1258134
 0.4368132  0.2115385  0.1813187  0.1703297
 0.4742857  0.1942857  0.1428571  0.1885714

MOTIF Dmelanogaster-FlyFactorSurvey-pdm3_SOLEXA_5_FBgn0033288
letter-probability matrix: alength= 4 w= 25 nsites= 100 E= 0
 0.1443649  0.2536751  0.237844  0.3641161
 0.05126272  0.00414625  0.000000  0.944591
 0.9932152  0.000000  0.00414625  0.002638522
 0.9962307  0.003769318  0.000000  0.000000
 0.001884659  0.000000  0.04221636  0.955899
 0.06068602  0.09423294  0.3339616  0.5111195
 0.525066  0.1372032  0.2039201  0.1338108
 0.08217113  0.171504  0.5876366  0.1586883
 0.121749  0.2340746  0.4063325  0.237844
 0.1752733  0.2080663  0.2027893  0.4138711
 0.2589521  0.1858274  0.2698832  0.2853374
 0.1858274  0.1741425  0.3490388  0.2909913
 0.1119487  0.2076894  0.3520543  0.3283076
 0.1062948  0.2261591  0.2751602  0.392386
 0.2427441  0.1858274  0.2423671  0.3290614
 0.1360724  0.2125895  0.3663777  0.2849604
 0.1929891  0.2103279  0.3071994  0.2894836
 0.1948737  0.2367132  0.2879759  0.2804372
 0.1696193  0.2189974  0.3022993  0.3090841
 0.1307953  0.204674  0.4055786  0.2589521
 0.154919  0.2303053  0.3584621  0.2563136
 0.1907275  0.2261591  0.3113456  0.2717678
 0.1684885  0.2091971  0.3467772  0.2755371
 0.1605729  0.2091971  0.3660008  0.2642292
 0.1853253  0.2155825  0.316944  0.2821483

MOTIF Dmelanogaster-FlyFactorSurvey-ss_tgo_SANGER_10_FBgn0015014
letter-probability matrix: alength= 4 w= 11 nsites= 100 E= 0
 0.2307692  0.000000  0.7692308  0.000000
 0.000000  0.000000  0.000000  1.000000
 0.000000  1.000000  0.000000  0.000000
 1.000000  0.000000  0.000000  0.000000
 0.000000  1.000000  0.000000  0.000000
 0.000000  0.000000  1.000000  0.000000
 0.000000  1.000000  0.000000  0.000000
 0.7692308  0.000000  0.2307692  0.000000
 1.000000  0.000000  0.000000  0.000000
 0.1538462  0.1538462  0.07692308  0.6153846
 0.000000  0.3846154  0.6153846  0.000000

MOTIF Dmelanogaster-FlyFactorSurvey-twi_da_SANGER_5_FBgn0000413
letter-probability matrix: alength= 4 w= 10 nsites= 100 E= 0
 0.5217391  0.173913  0.3043478  0.000000
 0.04347826  0.826087  0.08695652  0.04347826
 0.000000  0.9565217  0.000000  0.04347826
 1.000000  0.000000  0.000000  0.000000
 0.000000  0.1304348  0.6956522  0.173913
 0.9565217  0.000000  0.04347826  0.000000
 0.000000  0.000000  0.000000  1.000000
 0.000000  0.04347826  0.9565217  0.000000
 0.000000  0.08695652  0.173913  0.7391304
 0.000000  0.3181818  0.3181818  0.3636364
```
